# Profiling of epithelial functional states and fibroblast phenotypes in hormone therapy-naïve localised prostate cancer

**DOI:** 10.1101/2024.10.23.619925

**Authors:** Eva Apostolov, Daniel L Roden, Holly Holliday, Aurélie Cazet, Kate Harvey, Hanyun Zhang, Sunny Z Wu, Sophie van der Leij, Luke A Selth, Nenad Bartonicek, Ghamdan Al-Eryani, Mengxiao He, Joakim Lundeberg, John Reeves, James G Kench, Alison J Potter, Phillip Stricker, Anthony M Joshua, Lisa Horvath, Alexander Swarbrick

**Affiliations:** Cancer Ecosystems Program, Garvan Institute of Medical Research, Darlinghurst, NSW Australia; School of Clinical Medicine, Faculty of Medicine & Health, UNSW Sydney, Sydney, NSW Australia; Children’s Cancer Institute, Lowy Cancer Research Centre, UNSW Sydney, Sydney, NSW Australia; Flinders University, College of Medicine and Public Health, Flinders Health and Medical Research Institute, Bedford Park, SA Australia; Flinders University, Freemasons Centre for Male Health and Wellbeing, Bedford Park, SA Australia; Faculty of Health and Medical Sciences, The University of Adelaide, Adelaide, SA Australia; Department of Gene Technology, KTH Royal Institute of Technology, Science for Life Laboratory, Solna, Sweden; Tissue Pathology & Diagnostic Oncology, NSW Health Pathology, Royal Prince Alfred Hospital, Camperdown, NSW Australia; Sydney Medical School, University of Sydney, Camperdown, NSW Australia; Kinghorn Cancer Centre, Garvan Institute of Medical Research, Darlinghurst, NSW Australia; Faculty of Medicine & Health, University of Sydney, Camperdown, NSW Australia; St Vincent’s Prostate Cancer Research Centre, Sydney, NSW Australia; Department of Urology, St Vincent’s Private Clinic, Sydney, NSW Australia; School of Clinical Medicine, St Vincent’s Healthcare Clinical Campus, Faculty of Medicine & Health, UNSW Sydney, Sydney, NSW Australia; Chris O’Brien Lifehouse, Camperdown, NSW Australia; University of Sydney, Camperdown, NSW, Australia

## Abstract

Localised prostate cancers (PCa) are heterogeneous and multifocal, with diverse outcomes. Current prognostic methods are epithelium-centric, overlooking the complex cellular landscape within the tumour microenvironment (TME), which remains incompletely characterised. We performed a comprehensive analysis of cancerous and adjacent-benign cores from 24 patients with hormone therapy-naïve localised PCa using single-cell RNA-sequencing. By integrating copy number variation and transcriptional signatures, we classified epithelial cells across a malignant spectrum, revealing widespread molecular perturbation. We found an expansion of Club cell phenotypes, suggestive of Luminal dedifferentiation. We also performed a detailed annotation of stromal phenotypes, focusing on fibroblasts, and identified a novel peri-neural fibroblast population. Spatial transcriptomics elucidated the precise anatomical distribution of CAFs within the PCa TME. This study provides a valuable foundation for advancing our understanding of PCa pathobiology and developing a comprehensive cellular model of the disease.

**Statement of Significance:** Our study leverages single-cell RNA-sequencing and spatial transcriptomics to provide a comprehensive cellular annotation of hormone therapy-naïve localised PCa. We reveal widespread molecular perturbations in epithelial cells and map distinct fibroblast populations to specific anatomical niches. Notably, we identify a novel peri-neural phenotype associated with nerves, which merits further functional characterisation and exploration as a potential therapeutic target.

## Introduction

Prostate cancer (PCa) is a heterogeneous disease with clinical outcomes ranging from indolent to lethal metastatic disease (Rebello et al. 2021). The main growth pathway of PCa is androgen receptor (AR) signalling, which plays a critical role in the treatment of localised and metastatic disease. Current prognostic methods primarily focus on epithelial cells (Eastham et al. 2022), often overlooking the crucial role of the diverse cells in the tumour microenvironment (TME) in shaping disease progression and treatment response (Cazet et al. 2018; Gonzalez et al. 2018; Frankenstein et al. 2020). To optimise treatment and enhance response rates, a comprehensive understanding of the PCa cellular landscape in its native, hormone-sensitive state is essential.

The prostate epithelium comprises basal, androgen-responsive luminal cells and intermediate phenotypes - club and hillock cells (Henry et al. 2018). While the roles of these intermediate phenotypes in the human prostate are not fully understood, they may include immunomodulatory functions (Henry et al. 2018). Insights from mouse models suggest that these cells have increased progenitor potential and resistance to androgen deprivation (Joseph et al. 2020; Kwon et al. 2020; Guo et al. 2020; Karthaus et al. 2020; Germanos et al. 2022), indicating that intermediate phenotypes may play significant roles in PCa due to their less-differentiated and less androgen-dependent nature. The prostate epithelium of many men exhibits a spectrum of transformation, ranging from homeostasis to malignancy, as exemplified by copy number alterations in regions of seemingly benign epithelium (hereafter ‘altered benign’ epithelium) (Yu et al. 2012; Cooper et al. 2015; Erickson et al. 2022). However, current analytical approaches have not yet distinguished these epithelial functional states (normal, altered benign, malignant) and their lineages (basal, luminal, intermediate) at high resolution.

Disruptions in stromal homeostasis are also linked to disease progression (Pascal et al. 2009; Tyekucheva et al. 2017), biochemical recurrence-free survival (Ayala et al. 2003; Yanagisawa et al. 2007; McKenney et al. 2016), and PCa-specific mortality (Saeter et al. 2015). However, the diversity of stromal populations in PCa remains underexplored. Stromal phenotyping has historically employed a narrow set of protein markers - such as alpha-smooth muscle actin (*ACTA2*), vimentin (*VIM*), and CD90 (*THY1*) (Tuxhorn et al. 2002; Pascal et al. 2009) - which are expressed by multiple stromal lineages (Wu et al. 2020). This overlap complicates distinguishing even major lineages like cancer-associated fibroblasts (CAFs) from smooth muscle cells (SMCs), limiting the detection of novel and rare subpopulations that could be pivotal in the disease course. Therefore, precise delineation of epithelial and stromal phenotypes is crucial for advancing our understanding of PCa biology.

Single-cell RNA-sequencing (scRNA-seq) has offered unprecedented insights into the cellular landscape of solid tumours (Patel et al. 2014; Tirosh et al. 2016; Lambrechts et al. 2018; Wu, Al-Eryani, et al. 2021). In localised PCa, several scRNA-seq studies have compared cancer with adjacent benign samples, primarily focusing on immune composition (Tuong et al. 2021; Masetti et al. 2022; Adorno Febles et al. 2023) or assessment of epithelial states across different hormonal contexts, from hormone-naïve to castration-resistant and metastatic PCa (Cheng et al. 2022). Others have performed relatively comprehensive, ‘atlas-style’ profiling of the PCa TME (S. Chen et al. 2021; Song et al. 2022), but in all cases, sample sizes have been small: <10 patients (Tuong et al. 2021; Masetti et al. 2022; Adorno Febles et al. 2023; Song et al. 2022; Cheng et al. 2022), and 13 patients (S. Chen et al. 2021). Furthermore, detailed annotation of epithelial and stromal functional states is lacking. Equally significant is the paucity of spatial data (Marklund et al. 2022; Hirz et al. 2023), particularly when it comes to the spatial organisation of cellular phenotypes.

In this study, we used scRNA-seq and spatial transcriptomics to dissect epithelial functional states and CAF/SMC phenotypes in cancerous and adjacent-benign tissues from 24 hormone therapy-naïve localised PCa patients. This work has yielded a comprehensive transcriptional atlas of the cellular architecture of PCa, uncovering an expanded spectrum of epithelial and CAF cell states.

## Results

### A scRNA-seq atlas of hormone therapy-naïve localised PCa

Two PCa core biopsies were collected and viably cryopreserved (Wu, Roden, et al. 2021) from 24 patients undergoing radical prostatectomy (ST_1), with 7 patients providing two cancer samples and 17 patients providing one cancer and one adjacent-benign sample. The samples were pooled and processed for scRNA-seq profiling on the 10x Genomics platform using a Single Nucleotide Polymorphism (SNP)-based sample (de)multiplexing strategy (ST_2), aimed at maximising cellular yield while minimising sample handling time and batch effects (Fig. 1A).

**Figure 1:**
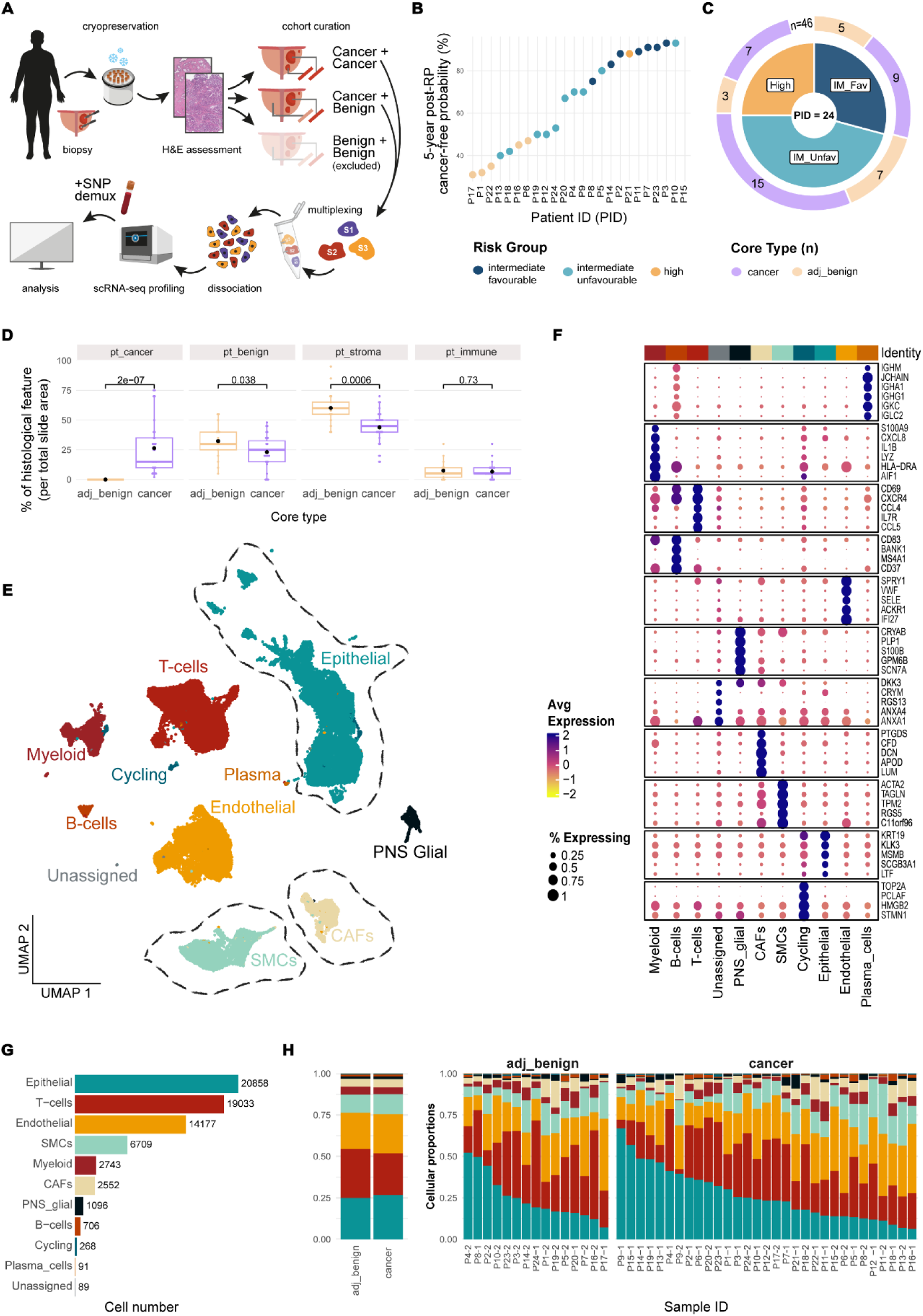
Overview of the experimental workflow, clinico-pathogical characteristics of the cohort and the major cell classes recovered in the scRNA-seq data. **A)** Schematic of the study design and sample processing for single-cell profiling on the 10x Chromium Platform. **B)** 5-year post-radical prostatectomy (RP) probability of remaining PCa-free, calculated using the MSKCC post-RP nomogram (Stephenson et al. 2005). **C)** Cohort composition. The inner pie represents the distribution of clinical risk groups (ISUP 2: intermediate favourable, ISUP 3: intermediate unfavourable, ISUP 4 and ISUP 5: high risk). The outer donut shows the number of ‘adjacent-benign’ and ‘cancer’ cores (samples) within each risk group. **D)** Histological feature percentages (‘pt’) of total slide area per core type. Dots indicate mean values, boxlines represent medians; *p*-values are from independent samples t-tests. **E)** UMAP visualisation of ***major*** cell types (Epithelial, T-cells, Endothelial, SMCs, Myeloid, CAFs, PNS_glial, B-cells, Cycling, Plasma cells, Unassigned). Dotted lines indicate the cell types of focus. **F)** Average expression of the top DEGs per ***major*** category. Negative values indicate expression below the mean across the dataset. **G)** Total cell number per ***major*** category. **H)** Left: ***major*** cell type abundance by core type (adj_benign, cancer). Right: ***major*** cell type abundance per sample ID, faceted by core type.

The patient cohort exhibited a broad spectrum of clinical risk (Intermediate Favourable, *n*=7; Intermediate Unfavourable *n*=11; and High, *n*=6) (Fig. 1B; (Stephenson et al. 2005)). Intermediate-risk patients dominated the cohort (Fig. 1C), reflecting the standard clinical incidence (Preisser et al. 2020). Of the 48 samples, two adjacent-benign samples were not recovered post demultiplexing, resulting in 31 cancer and 15 adjacent-benign samples (Fig. 1C; Supp. Fig. S1). For each sample, morphological assessment was done to assign the relative proportions of different histological features, including cancer, benign epithelium, stromal, and immune cells (Fig. 1D, ST_1). The median cancer cellularity in the cancer samples was 15% (mean = 26%), whereas morphologically benign epithelium was present in both adjacent-benign and cancer samples (Fig. 1D). Both cancer and adjacent-benign cores had substantial stromal and minimal immune components (Fig. 1D), consistent with previous reports on the cellular composition of PCa RP samples (Strand et al. 2016).

68,322 cells passed QC and filtering and were classified into eleven ***major*** cell classes based on cell type-specific markers (ST_3), including Myeloid, T-cells, B-cells, Plasma cells, Endothelial, CAFs, SMCs, Peripheral Nervous System (PNS) glial cells (including Schwann and Satellite cells), Epithelial, Cycling cells, and an Unassigned category (Fig. 1E). Notably, this is the first report of PNS glial cells in the PCa tumour microenvironment (TME) using scRNA-seq. The top differentially expressed genes (DEGs) per ***major*** cell type are shown (Fig. 1F). Epithelial cells were the most abundant, followed by T-cells and Endothelial cells (Fig. 1G). Consistent with other scRNA-Seq studies, adipocytes and neurons were not detected.

The ***major*** cell classes were variably represented across different core types and individual tumour samples (Fig. 1H). Grouping the data by core type (Supp. Fig. S2A) and clinical grade (Supp. Fig. S2B) did not reveal significant differences in ***major*** cellular composition, suggesting that interpatient heterogeneity in cellular composition extends beyond these two factors. Next, all ***major*** cell types were annotated at a higher resolution (i.e. ***minor***), reflecting established lineages (e.g., Luminal, Basal in Epithelial cells). Finally, cells were further subdivided based on clustering into ***subsets***, which typically correspond to cellular states (e.g., Luminal_MSMB+/c7, Basal_S100A2+/c2) (Supp. Fig. S3-S5). While this study primarily focuses on Epithelial cells, SMCs, and CAFs, a detailed cellular taxonomy with annotations to the ***subset*** level is available for all cell types (Supp. Fig. S6, ST_4) and will serve as a useful resource.

### Characterisation of the epithelial lineage reveals patient-unique and shared Luminal states and an abundance of antigen-presenting Club cells

We extracted, re-clustered and annotated epithelial cells using established marker genes and signatures of prostatic epithelial lineages (Henry et al. 2018) (Supp. Fig. S7 A, B). The resulting 20,858 cells were classified into seven ***minor*** categories: Luminal, Luminal-like, Club, Club-IFN, Basal, Neuroendocrine (NE), and Ciliated (Fig. 2A).

**Figure 2:**
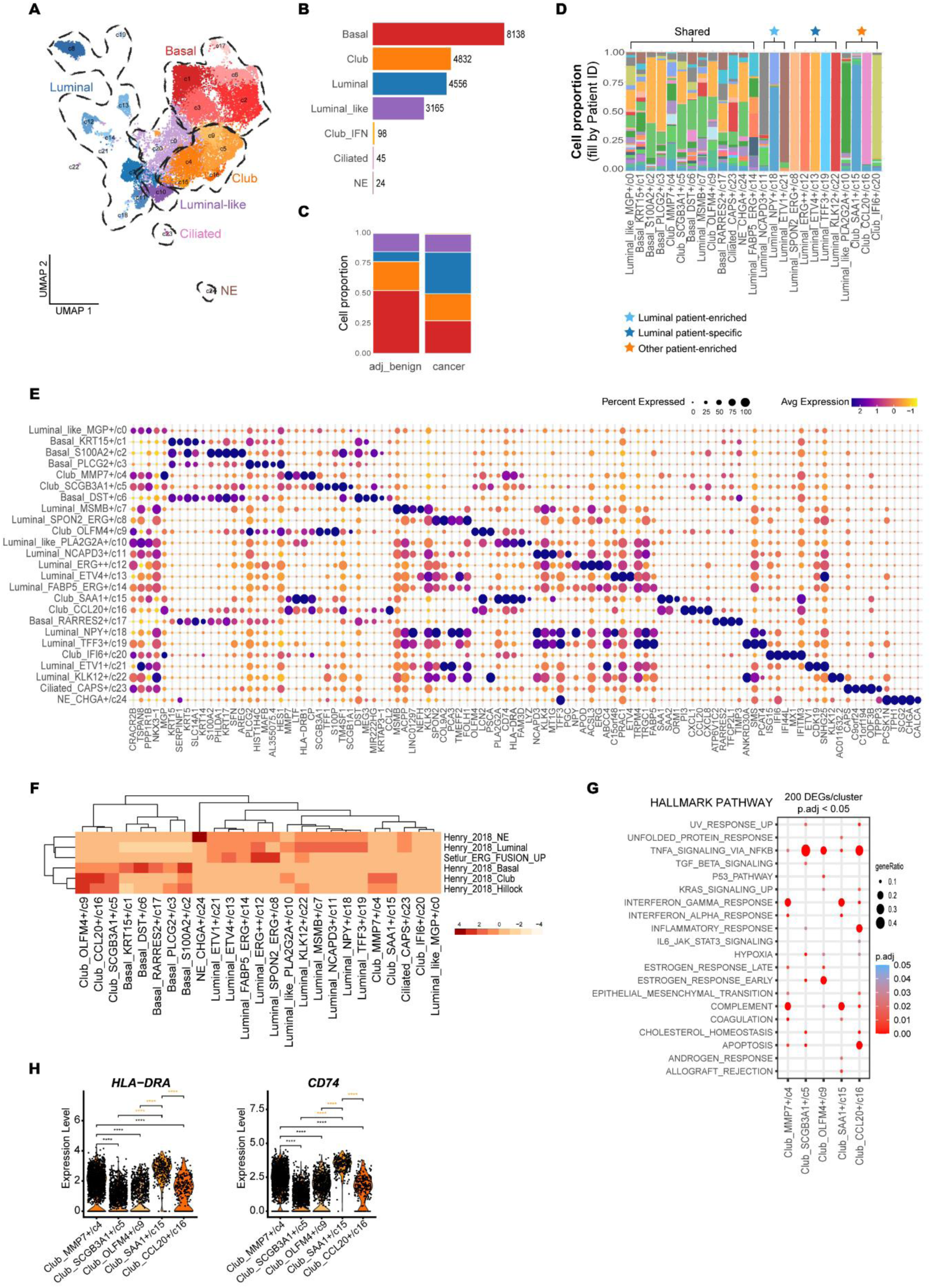
Characterisation of the epithelial lineage reveals patient-unique and shared Luminal states and an abundance of antigen-presenting Club cells. **A)** UMAP visualisation of ***subset*** annotation; dashed circles indicate ***minor*** lineages. **B) *minor*** cell type abundance. **C) *minor*** cell type abundance by core type (adjacent-benign, cancer). **D) *subset*** cluster composition, filled by 24 unique patients; clusters are labelled as ‘Shared’, ‘Luminal patient-enriched’ (“NCAPD3+/c11”, “NPY+/c18”, “ETV1+/c21”), ‘Luminal patient-specific’ (“SPON2_ERG+/c8”, ERG++/c12”, “ETV4+/c13”, “TFF3+c19”, KLK12+/c22”), and ‘Other patient-enriched’ (“Luminal_like_PLA2G2A+/c10”, “Club_SAA1+/c15”, “Club_CCL20+/c16”, “IFI6+/c20”). **E)** Average expression of the top 5 differentially expressed genes (DEGs) per ***subset*** category. **F)** Average AUCell enrichment scores per ***subset*** category for signatures of prostatic epithelial lineages (Henry et al. 2018) and of *TMPRSS2-ERG* fusion (Setlur et al. 2008). **G)** Over-representation analysis of the Hallmark Gene Set Collection (MSigDB) using the top 200 DEGs per Club cell ***subset***; pathways visualised with a 0.05 p-adj value cut-off. **H)** Expression of antigen-presenting machinery molecules, *HLA-DRA* (MHC II) and *CD74*, per Club cell ***subset***. Means compared with Wilcoxon test (unadjusted *p*-values; *p* ≤ 0.001 (****)). Black and orange asterisks indicate comparisons relating to “Club_MMP7+/c4” and “Club_SAA1+/c15”, respectively.

The epithelial landscape was dominated by Basal, followed by Club and Luminal cells (Fig. 2B). Club cells have been reported to make up ∼5% of epithelial cells in the peri-urethral regions of the normal prostate (Henry et al. 2018; Guo et al. 2020; Karthaus et al. 2020; Joseph et al. 2020). However, we found that they constituted ∼25% of epithelial cells in both cancer and adjacent-benign cores (Fig. 2C), suggesting a Club cell expansion in PCa. Hillock cells were not identified in this analysis, similar to previous reports in PCa (Song et al. 2022).

Luminal-like cells expressed both luminal (*KLK3*) and club (*PIGR*) cell markers, and were enriched for both Luminal and Club signatures (Supp. Fig. S7 A, B), therefore likely representing a state between Luminal and Club cells. Club-IFN cells represented a patient-specific (exclusive to a single patient) interferon (IFN)-enriched phenotype (Supp. Fig. S7 A, B).

The rare NE phenotype (*n* = 24 cells) was *CHGA^+^CALCA^+^* and enriched for the NE signature (Supp. Fig. S7 A, B). Ciliated cells (*n =* 45 cells) were *CAPS^+^* (Dinh et al. 2021) (Supp. Fig. S7B). While the role of Ciliated cells is not well understood, they have been implicated in regulating Hedgehog and WNT signalling pathways (Eguether & Hahne 2018), and are suggested to become dysfunctional in PCa (Hassounah et al. 2013).

At a higher resolution (***subset*** level), we detected twenty-five clusters: ten Luminal (*KLK3, NKX3-1, MSMB*), two Luminal-like (*KLK3, PIGR*), five Club (*SCGB3A1, LCN2, PIGR*), five Basal (*KRT5, KRT15, DST, TP63*) clusters, and one cluster each of NE, Ciliated, and Club-IFN cells (Fig. 2A, E, F; Supp. Fig. S7C). Epithelial cluster diversity was more pronounced in the cancer cores (Supp. Fig. S7 D, E), with eight Luminal clusters and four non-luminal clusters (“Luminal_like_PLA2G2A+/c10”, “Club_SAA1+/c15”, “Club_CCL20+/c16”, “Club_IFI6+/c20”) exhibiting patient-enriched (enriched in, but not exclusive to, a single patient) or patient-specific patterns (Fig. 2D). This likely reflects intertumoural genomic and/or transcriptional heterogeneity, typically observed as patient-specific clusters in tumour scRNA-seq data (Wu, Al-Eryani, et al. 2021), suggesting the potential presence of malignant and/or genomically-perturbed cells within these clusters. The remaining clusters were shared among patients (Fig. 2D), indicative of commonly occurring epithelial phenotypes.

Among the Basal clusters, clusters “KRT15+/c1” and “DST+/c6” exhibited a ‘classical’ basal phenotype (*KRT5, KRT15^+;^* Fig. 2E; Supp. Fig. S7C). In contrast, cluster “RARRES2+/c17” resembled ionocytes (*KRT5, KRT15^+^* and *RARRES2, FOXI1^+^*; Fig. 2E; Supp. Fig. S7C), a rare pulmonary basal epithelial cell involved in ion balance (Plasschaert et al. 2018). Analogous profiles have been observed in scRNA-seq of mouse prostate and IHC studies of human prostate (Karthaus et al. 2020). Additionally, clusters “S100A2+/c2” and “PLCG2+/c3” were *KRT5^+^KRT13^+^* and enriched in Basal and Hillock signatures (Fig. 2E, F), indicating a possible enrichment of Hillock features in these cells.

In the Luminal cells, we detected members of the ETS transcription factor (TF) family (*ERG, ETV4, ETV1*) among the top differentially expressed genes (DEGs) for several Luminal clusters (“SPON2_ERG+/c8”, “ERG++/c12”, “ETV4+/c13”, “FABP5_ERG+/c14”, “ETV1+/c21”; Fig. 2E). ETS family transcription factors are not typically expressed by Luminal cells (Qian et al. 2022), except when ETS fusion events occur in PCa (Cancer Genome Atlas Research Network 2015). This suggested that these clusters likely comprise ETS fusion positive PCa cells, consistent with their high average activity of a *TMPRSS2-ERG* fusion signature (Fig. 2F; (Setlur et al. 2008)). The remaining Luminal clusters likely represented various ETS-negative PCa phenotypes, except for “Luminal_MSMB+/c7”, which resembled canonical acinar luminal epithelium characterised by high *MSMB* expression (Fig. 2E) - a gene often downregulated in PCa and cancer-adjacent tissues (Bergström et al. 2018; Crowley et al. 2020).

Club cells are well-characterised in the healthy lung and lung cancer (Yuanyuan Chen et al. 2022; Blackburn et al. 2023), but their functional significance in the healthy prostate and PCa remains unclear (Henry et al. 2018; Song et al. 2022). To better understand their characteristics and given their prevalence in our dataset, we further analysed the five Club clusters using the Hallmark Gene Set Collection. Club cluster “CCL20+/c16” showed signs of inflammation and apoptosis (Fig. 2G), potentially indicating its origin from ‘injured’/atrophic glands. Clusters “SCGB3A1+/c5” and “OLFM4+/c9” were *SCGB3A1^high^* (Fig. 2E), a marker previously suggested to indicate a ‘resting-state’ Club cell phenotype in healthy prostates (Song et al. 2022). These clusters were also enriched in TNFα signalling (Fig. 2G), potentially indicative of an anti-inflammatory role (Rawlins et al. 2009). Clusters “MMP7+/c4” and “SAA1+/c15” were enriched in IFN-gamma (IFNγ) response and complement signalling (Fig. 2G). IFNγ enhances antigen presentation (Mootz et al. 2022), and the high expression of antigen presentation machinery in these clusters (*HLA-DRA* (MHC II); *CD74*; Fig. 2H; (Wosen et al. 2018)) suggested they might modulate immune responses in the PCa TME as non-professional antigen-presenting cells.

Overall, we highlight patient-specific and shared Luminal states and report an ionocyte-like cell in human PCa scRNA-seq data. Additionally, we uncover previously unrecognised Club cell heterogeneity and their prevalence in PCa, along with their expression of antigen-presentation machinery.

### The malignant spectrum in PCa enables comparison of AR signalling within luminal states

An important challenge in PCa is the differentiation between benign, altered benign, and malignant epithelium (Yu et al. 2012; Erickson et al. 2022). To address this, we first used single-cell copy number variation (CNV) analysis using *CopyKAT (Gao et al. 2021)* to identify CNV-transformed cells in the prostatic epithelium. We refer to these cells as CNV^pos^ rather than malignant, to account for the potential presence of altered benign epithelium.

Cells were initially classified as ‘CNV^pos^’ or ‘CNV^neg^’, with CNV^neg^ cells found in both adjacent-benign and cancer cores (Supp. Fig. S8A), consistent with histopathological observations of ‘benign’ epithelium in both core types (Fig. 1D). Despite challenges in classifying cells due to lower CNV burden in localised PCa compared to other solid tumours (Cancer Genome Atlas Research Network 2015), such as breast cancer (Wu, Al-Eryani, et al. 2021) or melanoma (S. Chen et al. 2021), CNVs commonly associated with PCa were detected in both core types. For instance, chr8q gain (including *MYC*) was detected in a cancer core (Fig. 3A), and chr3q gain (including *PI3K*) in an adjacent-benign core (Fig. 3B).

**Figure 3:**
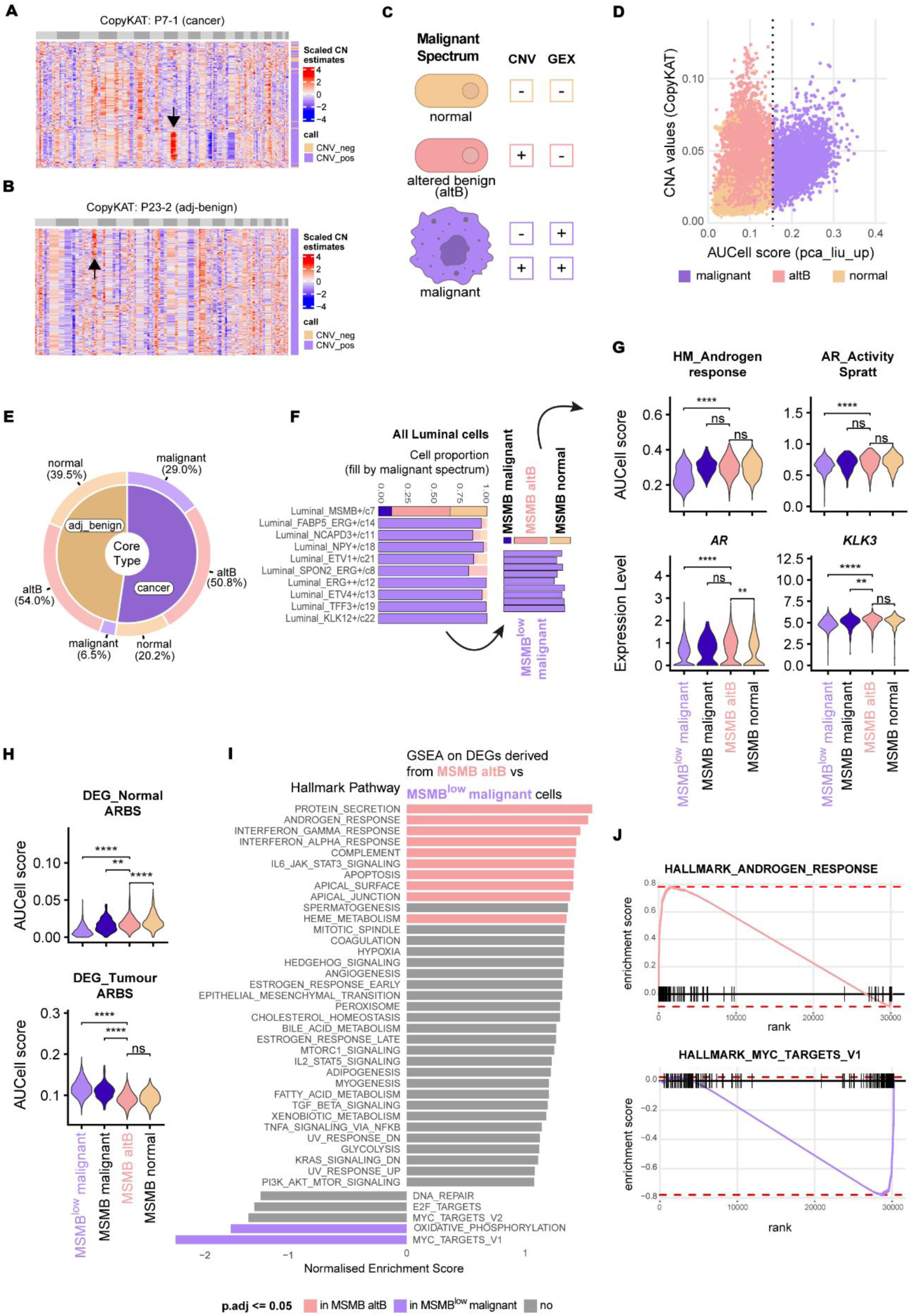
Androgen receptor signalling across the malignant spectrum in PCa A-B) Heatmaps of copy number estimates (A: cancer sample P7-1; B: adj-benign sample P23-2); Black arrows show chr8q (A) and chr3q gain (B). Values are *z*-score scaled CN signals. Cells (rows) are clustered by CNV signal. Row dendrograms are not shown; chromosomes 1-22 are marked with alternating colours (odd: light grey; even: dark grey). **C)** Malignant spectrum states: ‘malignant’ (CNV^pos^/Sig^pos^ or CNV^neg^/Sig^pos^), ‘altered benign’ (altB; CNV^pos^/Sig^neg^), and ‘normal’ (CNV^neg^/Sig^neg^), reflecting differences between gene expression (PCa signatures and AUCell) and CNV status. **D)** Correlation of CNA values (y-axis; *CopyKAT*) versus AUCell scores for the PCa-defining signature (x-axis: “pca_liu_up” (Liu et al. 2006)); dashed line indicates AUCell threshold for “pca_liu_up” (0.15). Cells are coloured by malignant spectrum state. **E)** Cumulative abundance (%) of malignant spectrum states (donut) per core type (pie). **F)** Malignant spectrum classification across Luminal ***subset*** clusters; MSMB+ states (malignant, altB, normal) and MSMB^low^ malignant cells are highlighted in a darker tint. **G)** AUCell scores for “HM.Androgen_response” and “AR_Activity_Spratt”, and *AR* and *KLK3* expression across MSMB^low^ malignant cells and the three MSMB+ states. **H)** AUCell scores for “DEG_Normal_ARBS” and “DEG_Tumour_ARBS” (Pomerantz et al. 2015) across the same epithelial states as in (G). In (G) and (H), gene expression means are compared with the Wilcoxon test, and AUCell scores with a t-test (unadjusted *p*-values; *p* ≤ 0.001 (****)). **I)** Gene set enrichment analysis on avg_logFC-ranked DEGs between altB MSMB+ and MSMB^low^ malignant cells, highlighting significantly enriched pathways (*p-ad*j ≤ 0.05). 40 out of 50 Hallmark pathways with NES values < -1 (enriched in MSMB^low^ malignant) or >1 (enriched in altB MSMB+ cells) are shown. **J)** Enrichment plots, with DEGs ranked by avg_logFC (x-axis). altB = altered benign; avg_logFC = average log fold change; DEGs = differentially expressed genes; NES = Normalised Enrichment Score(s).

To identify malignant cells, we next assessed the enrichment of published PCa signatures, including “pca_liu_up” (Liu et al. 2006) and “pca_wallace_up” (Wallace et al. 2008). We expected these signatures, which were derived from comparisons of morphologically malignant to benign epithelium, to be enriched in malignant but not in altered benign or normal epithelium. Both signatures were PCa-specific, with minimal enrichment in young-adult normal prostatic epithelium (Supp. Fig. S8 C, D). Epithelial cells were assigned as positive or negative for the cancer signatures (‘Sig^pos^’ or ‘Sig^neg^’) based on AUCell-assigned signature thresholds. We then integrated this transcriptomic information with CNV status to classify cells along a malignant spectrum, from normal (Sig^neg^ /CNV^neg^) to altered benign (Sig^neg^ /CNV^pos^) to malignant (Sig^pos^ /CNV^pos^ or Sig^pos^ /CNV^neg^) (Fig. 3C, Supp. Fig. S9A). Altered benign cells had high CNA values but scored below the PCa signature thresholds (Fig. 3D), indicating benign gene expression features despite their CNV^pos^ status.

The three cell states were present in both core types (Fig. 3E). However, malignant cells were heavily enriched in cancer cores, accounting for 29% of the epithelium compared to 6.5% in adjacent-benign cores (Fig. 3E). A small percentage of malignant cells were also detected in non-luminal phenotypes, with 1.35% in Club and 0.04% in Basal cells (Supp. Fig. S9 B, C).

Altered benign cells comprised 50.8% of the epithelium in cancer and 54% in adjacent-benign cores (Fig. 3E), indicating that cells harbouring CNVs but with gene expression profiles distinct from malignant cells are frequent in both core types. Altered benign cells were detected throughout the epithelial lineage (Supp. Fig. S9 B, C), including in Basal (Supp. Fig. S10 A, B) and Club (Supp. Fig. S10 C, D) cells from individual patients, suggesting the presence of CNVs in non-luminal cells. Specifically, altered benign Club cells mapped to the patient-enriched Club cluster “Club_CCL20+/c16” (Fig. 2D). The presence of CNVs and the enrichment in inflammatory and atrophic processes in this state (Fig. 2G), suggested that this cluster may represent luminal cells that have acquired Club-like properties in response to copy number change. Overall, this data indicates widespread genomic and gene expression perturbation of the prostatic epithelium.

Previous scRNA-seq studies have only distinguished malignant and normal epithelium, without addressing the altered benign state (S. Chen et al. 2021; Song et al. 2022; Graham et al. 2024). To address this gap, we focused on the altered benign state in luminal cells - the primary androgen-responsive lineage. We identified this state primarily within the acinar luminal population “Luminal_MSMB+/c7” (hereafter ‘MSMB+ cells’) across multiple patients (Fig. 3F, Supp. Fig. S11A). Given that PCa predominantly exhibits an acinar luminal phenotype (Li & Wang 2016), MSMB+ cells served as an ideal reference for examining how clinically relevant features, such as AR signalling, change across luminal functional states, particularly during the transition from altered benign to malignant epithelium.

We categorised luminal cells into four groups: three functional MSMB+ states (normal MSMB+, altered benign MSMB+, malignant MSMB+), and a fourth group, MSMB^low^ malignant cells, representing malignant cells from other luminal clusters (Fig. 3F;Supp. Fig. S11B). This allowed comparison of **(1)** altered benign MSMB+ cells to their normal and malignant MSMB+ counterparts, and **(2)** altered benign MSMB+ cells to MSMB^low^ malignant cells, the predominant malignant phenotype (Fig. 3F). Higher proportions of MSMB+ malignant compared to MSMB^low^ malignant cells have been associated with improved biochemical recurrence-free survival (Dahlman et al. 2011), highlighting significant biological differences between these phenotypes, which our grouping aimed to capture.

To assess differences in AR activity and androgen response across the four groups, we used: **i)** “AR_activity_Spratt”, comprising nine canonical AR target genes (Spratt et al. 2019); and **ii)** “HM.androgen response” - the hallmark androgen response (Liberzon et al. 2015). Altered benign MSMB+ cells had similar levels of androgen signalling, *AR*, and *KLK3* expression as normal and malignant MSMB+ cells (Fig. 3G). In contrast, altered benign MSMB+ cells had significantly higher androgen signalling than MSMB^low^ malignant cells, including *AR* and *KLK3* expression (Fig. 3G), with similar results observed at the intratumoural level in two patients (Supp. Fig. S11 C-J). This indicated a marked downregulation of canonical androgen signalling in malignancy that becomes particularly pronounced with the loss of the MSMB+ phenotype.

Epithelial AR signalling is context dependent, driving different transcriptional programs in homeostasis and PCa, with accompanying alterations in AR chromatin binding patterns (Pomerantz et al. 2015; Copeland et al. 2019). Consequently, we expanded our analysis with two additional gene signatures: **i)** “DEG_Normal_ARBS” and **ii)** “DEG_Tumour_ARBS”, representing genes from DGEA of normal prostate versus PCa in TCGA data. “DEG_Normal_ARBS” includes direct AR target genes upregulated in normal prostate, whereas “DEG_Tumour_ARBS” includes direct AR target genes upregulated in primary PCa tumours (Pomerantz et al. 2015). Altered benign MSMB+ cells were enriched in normal AR activity, akin to normal MSMB+ cells, while malignant cells were enriched in cancer-associated AR activity (Fig. 3H). Additionally, altered benign MSMB+ cells were enriched in protein secretion and immune responses compared to MSMB^low^ malignant cells, which exhibited upregulation of MYC activity and oxidative phosphorylation (Fig. 3I, J). *In vitro* studies have suggested that MYC activity could antagonise the androgen-induced AR transcriptional program (Barfeld et al. 2017; Guo et al. 2021), suggesting coordinated reciprocal dysregulation of these pathways in malignancy.

In summary, altered benign MSMB+ cells resemble their normal MSMB+ counterparts with respect to androgen signalling, despite harbouring CNVs. It is the loss of the MSMB phenotype that is associated with marked downregulation of canonical androgen signalling and coordinated upregulation of MYC activity.

### Characterisation of SMCs and CAFs in the PCa TME

The prostatic stromal landscape is poorly characterised, with common CAF immunohistochemical markers also expressed by other stromal cells (Supp. Fig. S12). Since ∼50% of the cores in our dataset comprised stroma (Fig. 1D), we focused on delineating its cellular constituents, with an emphasis on SMCs and CAFs.

#### SMCs

SMCs were the fourth most abundant cell type (*n* = 6,709) and clustered into four ***minor*** populations (Supp. Fig. S13A). All were characterised by high expression of *ACTA2*, with three also expressing the mural cell marker *PDGFRB (Bondjers et al. 2003)* (Supp. Fig. S13D). This suggested the presence of two distinct SMC classes - contractile prostatic SMCs (pSMCs) and mural cells, as has been reported in the normal prostate (Joseph et al. 2021). The presence of these distinct SMC classes, each with unique transcriptomic profiles, distinguishes the prostate from other glandular tissues.

Mural cells were the most abundant SMC (Supp. Fig. S13B), comprising vascular SMCs (vSMCs; *RERGL*+/*BCAM*+/*PLN+*) and Pericytes (*THY1+*/ *COL6A3+*/*GGT5+*), followed by pSMCs (*ACTG2+*/*CNN1+* and *PDGFRB^low^*), and a patient-specific IFN^high^ phenotype, SMCs_IFN (*ISG15+*/*IFI6*+; Supp. Fig. S13 C, D). Gene ontologies for vSMCs were enriched in muscle contraction and development (Supp. Fig. S13E), reflecting the association of vSMCs with major blood vessels (Joseph et al. 2021). Pericytes were enriched in wound healing, ECM organisation, and chemotaxis (Supp. Fig. S13E), aligning with their association with minor vessels, angiogenesis, and immune trafficking (Duan et al. 2018; Kang et al. 2019; Joseph et al. 2021). pSMCs were enriched in muscle contraction, negative regulation of locomotion, and focal adhesion (Supp. Fig. S13E). vSMCs and Pericytes have been referred to interchangeably or considered a homogenous group (Teuwen et al. 2021; Hirz et al. 2023) due to a lack of distinguishing markers. For instance, *CSPG4* (encoding NG2), sometimes reported as a pericyte marker (Hosaka et al. 2016), was enriched in vSMCs in our data (Supp. Fig. S13D). Additionally, markers previously reported for Pericytes (*RGS5*; (Mitchell et al. 2008)) and vSMCs (*NDUFA4L2*; (Yang Chen et al. 2022)) were variably expressed in both mural cell types (Supp. Fig. S13D). Overall, this suggested a potential PCa-specific mural gene expression. We identified *RERGL* as a marker of vSMCs in the prostate (Supp. Fig. S13 C, D), which could help differentiate these cell types in future *in situ* and functional studies.

#### CAFs

Fibroblasts were identified by the expression of the canonical markers *DCN* and *PDGFRA* (Fig. 4C), and their functional characteristics were inferred through DGEA and gene ontology (GO) analysis. This was complemented by assessing the expression of canonical markers of **i)** fibroblast activation (e.g., *ACTA2, FAP*) (Pederzoli et al. 2023), **ii)** TGFb signalling (*TGFB1* and receptors *TGFBR1/2*) (Franco et al. 2011), and **iii)** fibroblast specialisation (inversely correlated with *DPT* expression) (Buechler et al. 2021) (Fig. 4C). Ligand-receptor (L-R) interactions were also analysed with CellChat (Jin et al. 2021) (Supp. Fig. S14).

**Figure 4:**
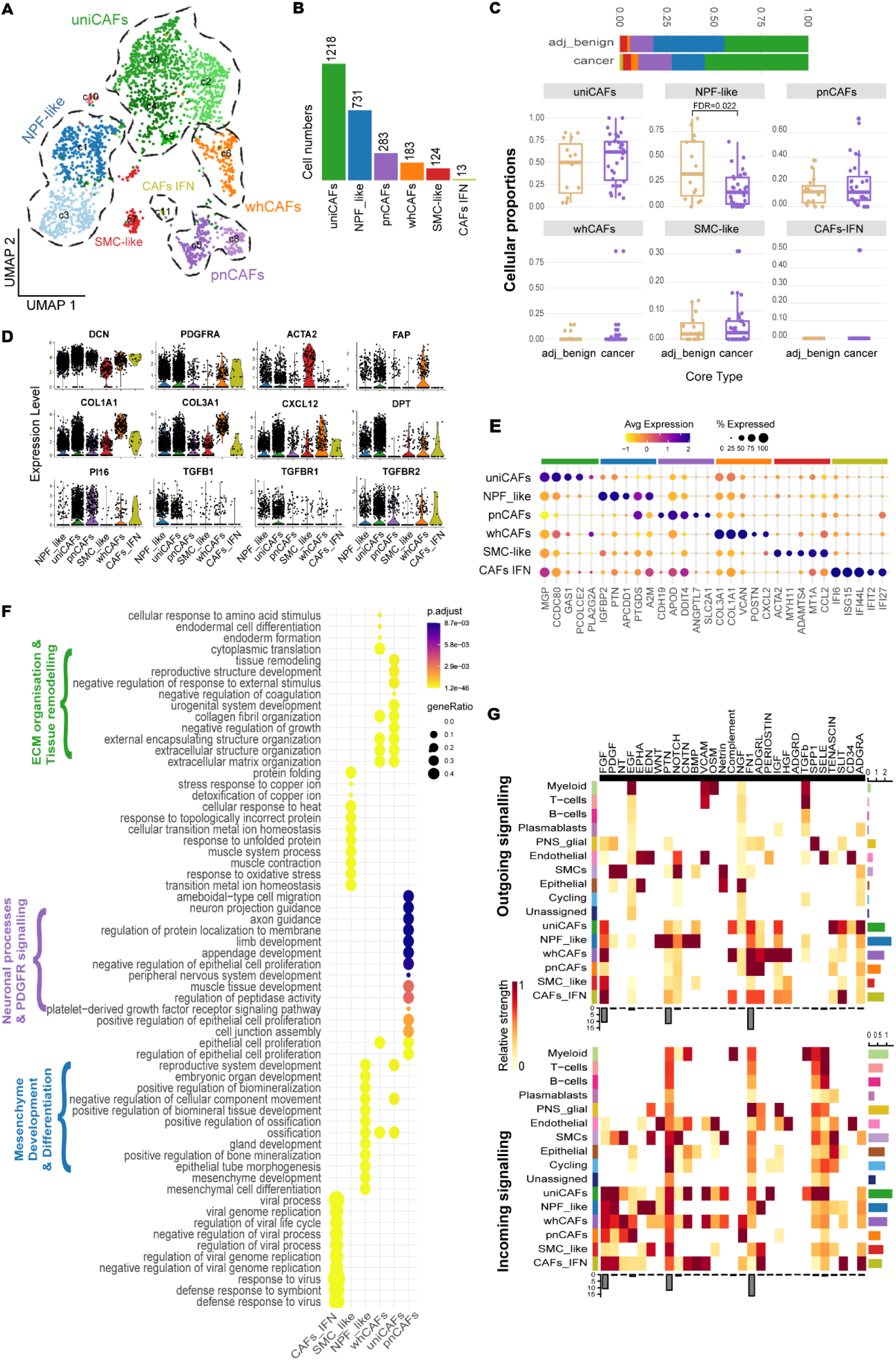
Characterisation of fibroblast populations in the PCa TME reveals unique and novel phenotypes. **A)** UMAP representation of ***minor*** CAF annotation. **B) *minor*** cell type abundance. **C)** Cumulative ***minor*** cellular proportion by core type (top), and ***minor*** cellular proportions by core type; denominator is all fibroblasts from all patients; individual data points represent samples. NPF-like cells are significantly enriched in adjacent-benign cores (Asin FDR = 0.046; Logit FDR = 0.023; *speckle (Phipson et al. 2022)*). **D)** Expression of canonical fibroblast (*DCN, PDGFRA*), myofibroblasts-associated (myCAFs; *COL1A1, COL3A1, ACTA2, FAP*), immunomodulatory CAFs (iCAFs; *CXCL12*); universal fibroblasts (*DPT, PI16*), and TGFb signalling markers (*TGFB1, TGFBR1, TGFBR2*). **E)** Average expression of the top 5 differentially expressed genes (DEGs) per ***minor*** category. **F)** Over-representation analysis (*clusterProfiler*) using the top 150 DEGs per ***minor*** category, showing enrichment of the top 15 biological processes. **G)** CellChat heatmaps displaying selected outgoing (top) and incoming (bottom) signals (x-axis) across all cell types. Cell types are grouped by ***major*** category, except for CAFs, which are grouped by ***minor*** category (y-axis). Colour values indicate relative strength of each signalling pathway, scaled column-wise per heatmap. Grey barplot (bottom) indicates overall signalling strength for each pathway; coloured barplot (right) shows total signalling strength per cell type (all pathways summarised). Full list of signalling pathways is shown in Supp. Fig. S14.

Our analyses reveal an unexpected degree of CAF heterogeneity, with a total of 2,552 fibroblasts assigned to six ***minor*** categories: universal-like CAFs (uniCAFs), wound-healing CAFs (whCAFs), normal prostate fibroblast-like (NPF-like), peri-neural CAFs (pnCAFs), smooth muscle cell-like (SMC-like), and an IFN-enriched phenotype (CAFs-IFN; Fig. 4A). uniCAFs and NPF-like were the most prevalent (Fig. 4B). The use of ‘CAFs’ rather than ‘fibroblasts’ in our nomenclature is due to their distribution in both adjacent-benign and cancer cores (Fig. 4C), and their gene expression features (explained below). NPF-like cells were significantly more abundant in adjacent-benign cores (FDR = 0.023, Fig. 4C), suggesting they resemble normal fibroblasts, while the remaining populations were present in both core types (Fig. 4C). Of these, uniCAFs trended towards prevalence in cancer cores (Fig. 4C), suggesting a potential inverse correlation or spatial isolation between them and NPF-like cells. CAFs-IFN were only detected in cancer cores (Fig. 4C).

uniCAFs, the predominant CAF population, displayed high *DPT* and *PI16* expression (Fig. 4D). This is consistent with a universal fibroblast phenotype found across many tissues, proposed to serve as a reservoir for fibroblast functional specialisation in both steady-state and perturbed conditions (Buechler et al. 2021). They also expressed *PCOLCE2, MGP,* and *CCDC80* (Fig. 4E), which encode factors involved in ECM organisation and tissue remodelling (Fig. 4F), in line with ECM remodelling being one of the primary roles of universal fibroblasts (Buechler et al. 2021). uniCAFs were primary producers of CD34 and Tenascin signalling molecules (Fig. 4G, Supp. Fig. S15A), which are linked to progenitor-like properties and microvessel density, respectively (Buechler et al. 2021; Lu et al. 2023; Ni et al. 2017). Alongside whCAFs, they were predicted by CellChat to be major sources of complement signalling to Myeloid cells (Fig. 4G), implying involvement in immunomodulatory processes (K. Chen et al. 2021). uniCAFs had the highest responder score out of the CAF populations to TGFb signalling (Fig. 4G), suggesting enriched TGFb activation, consistent with their ability to accept specialisation cues. They are also predicted to respond to SELE signalling from Endothelial cells, which is involved in vascular immunoadhesion during inflammation (Guitton et al. 2011) (Fig. 4G), suggesting that uniCAFs may also modulate immune responses associated with vascular features.

The whCAFs population exhibited a *FAP^high^*/*ACTA2^low^* phenotype (Fig. 4D), expressed collagens (*COL3A1, COL1A1*; Fig. 4E), and was enriched in ECM deposition and organisation processes (Fig. 4F), indicative of a wound healing rather than a ‘classical’ myofibroblast-like CAF (myCAF) state (*ACTA2^high^*/*FAP^high^*/*COL+*) (Wu et al. 2020; Werba et al. 2023). A clear myCAF phenotype was not identified in this analysis. whCAFs were the main producers of growth factors, such as HGF, IGF and Periostin signalling (Fig. 4G, Supp. Fig. S15B), suggesting their involvement in growth, repair, and vascularisation processes. whCAFs were predicted to respond to incoming signalling cues such as Netrin, OSM, and VCAM signalling (Fig. 4G). Netrin acts as a vascular and axonal mitogen (Park et al. 2004); OSM, a myeloid cell-derived cytokine, promotes inflammatory gene expression in pancreatic cancer CAFs (Lee et al. 2021); and VCAM regulates trans-endothelial migration of leukocytes (Carman & Springer 2004). Additionally, whCAFs and uniCAFs exhibited similar response patterns (Supp. Fig. S14), suggesting functional similarity and potential co-localisation within similar spatial niches. Overall, the CellChat analyses suggested that both populations may associate with blood vessels, potentially influencing immune cell migration.

NPF-like cells expressed growth factors and prostaglandin-metabolism genes (*IGFBP2, PTGDS;* Fig. 4E), and showed enrichment in mesenchyme development, differentiation, and epithelial tube morphogenesis (Fig. 4F). Within the fibroblast populations, NPF-like cells were the major source of *TGFB1*, the TGFb ligand (Fig. 4D), and the primary source of BMP and WNT signalling (Fig. 4G, Supp. Fig. S15C). BMP signalling is involved in prostatic epithelial differentiation (Omori et al. 2014), whereas WNT signalling suppresses prostatic epithelial proliferation (Wei et al. 2019), suggestive of a role for NPF-like cells in normal tissue development and epithelial homeostasis. Consistent with an interaction with epithelial cells, NPF-like cells were also the main responders to EDN and EPHA signals from Epithelial cells (Fig. 4G).

pnCAFs were a novel state and the only *DPT^low^* population (Fig. 4D) - a phenotype associated with fibroblast functional specialisation (Buechler et al. 2021). High expression of *PI16* (Fig. 4D), which is associated with pain sensitisation (Singhmar et al. 2020); glucose uptake and lipoprotein molecules (*SLC2A1, APOD, CDH19*; Fig. 4E); and enrichment in axon and neuron projection guidance processes (Fig. 4F), suggested pnCAFs might be a specialised state that associates with neurons. Accordingly, pnCAFs represented sources of ADGRL and FN1 signalling molecules, with Glial cells among the primary responders (Fig. 4G; Supp. Fig. S15D). pnCAFs were also the primary responders to FGF, PDGF, EGF signalling, including from Glial cells (Fig. 4G; Supp. Fig. S14B), suggesting this specialised phenotype may be recruited to neuronal regions.

In summary, uniCAFs exhibited a universal fibroblast-like phenotype with roles in ECM organisation and immune response coordination. L-R signalling predicted uniCAFs and whCAFs share niches linked to blood vessels, while uniCAFs and NPF-like cells show spatial separation, with NPF-like cells enriched in adjacent-benign cores. Lastly, pnCAFs represented a novel phenotype possibly closely associated with neurons.

### Spatial mapping of data reveals distinct anatomical organisation of cell types

To test our hypotheses about the spatial distributions of CAF and SMC populations, as well as the localisation of the previously detected Luminal-like and Club cells, we generated Visium FFPE data from 10 samples (3 adjacent-benign, 7 cancer, with 9 matching the scRNA-seq data, ST_5). A specialist prostate anatomical pathologist, blinded to the gene expression data, annotated the H&E images (Fig. 5A), with the histopathological annotations closely aligning with gene expression clusters (Supp. Fig. S16 A, B). We estimated cell numbers and proportions (per 50 µm Visium spot) with Cell2Location (Kleshchevnikov et al. 2022), using our scRNA-seq data as a reference (Fig. 5B).

**Figure 5:**
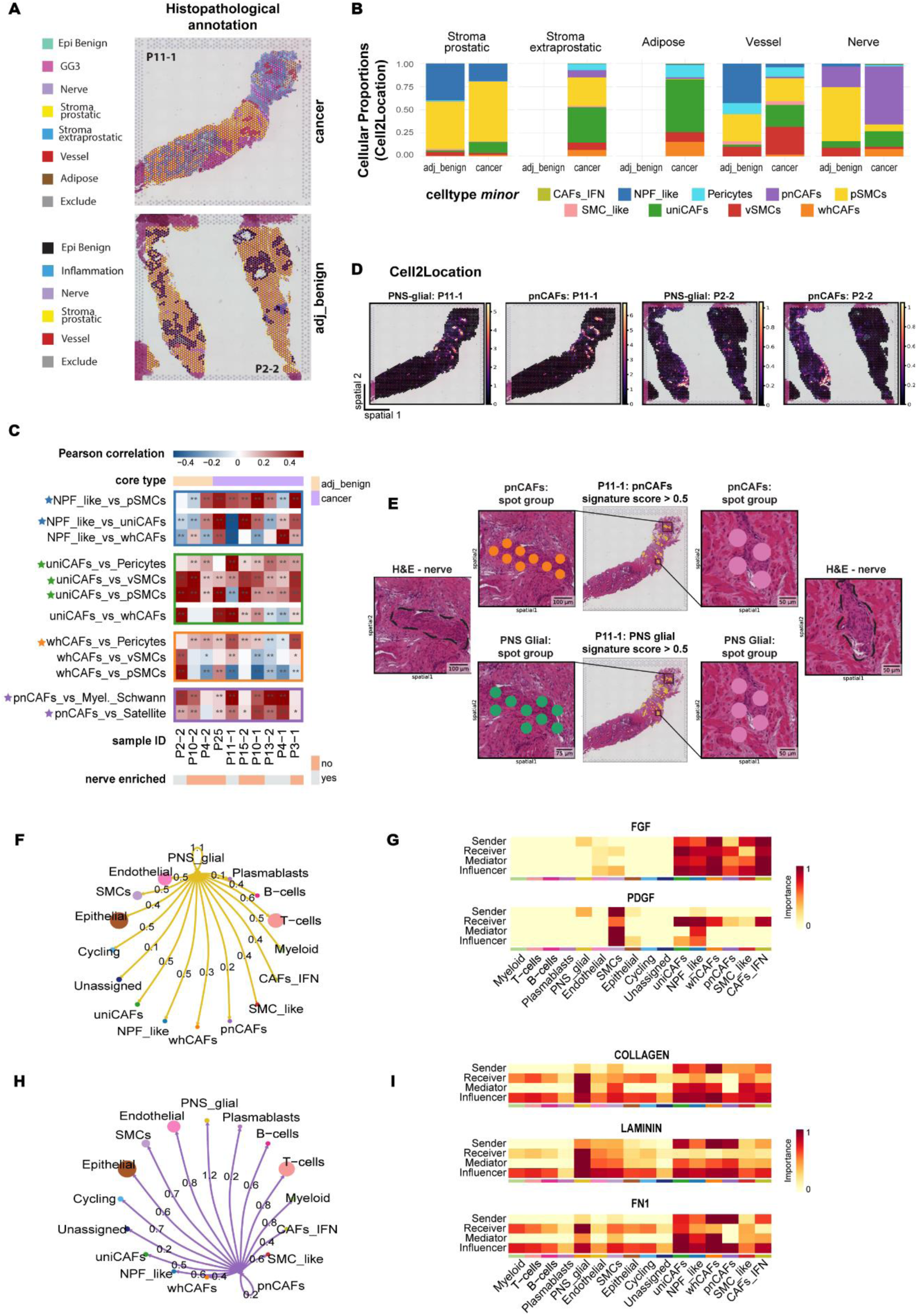
Spatially resolved analysis of Visium FFPE data reveals distinct anatomical distributions and interactions of CAF subsets with SMCs and Glial cells. **A)** Overlay of per-spot pathological annotations for cancer (top) and adjacent-benign (bottom) sections, indicating Epithelium (Epi Benign, Gleason Grade 3), Stroma (Prostatic, Extraprostatic), Nerves, Vessels, Adipose Tissue, and Inflammation. “Exclude” marks areas outside the tissue or within gland lumens. Additional 8 samples are shown in Supp. Fig. S16. **B)** Cell2Location deconvolution proportions for CAF and SMC ***minor*** cell types across histological regions, for all 10 samples, split by core type (x-axis). “Stroma Extraprostatic” and “Adipose” tissue are not found in adjacent-benign cores. **C)** Pearson correlation values between ***minor*** CAF, SMC, and PNS_glial subtypes across 10 samples (3 adjacent-benign, 7 cancer). Full comparison shown in Supp. Fig. S19D. Asterisks are Benjamini-Hochberg adjusted *p*-values: * < 0.01, ** < 0.001, with stars highlighting key interactions. **D)** Cell2Location per-spot deconvolution values for pnCAFs (***minor***) and PNS_glial cells (***major***), overlaid on H&E images of two nerve-enriched samples. **E)** Overlap of high-scoring spots for Cepo-derived PNS_glial and pnCAF signatures with nerves in a nerve-enriched sample. Scores are calculated using ‘scanpy.tl.score_genes()’, with spots scoring > 0.5 highlighted in yellow (spatial plots in the middle; pnCAFs score: top, PNS_glial score: bottom). Black rectangles indicate spot groups. H&E images show detailed regions with nerves outlined by dashed lines. Scale bars (50, 75, and 100μm) are indicated on each image. **F)** Circle plot of aggregated outgoing ligand-receptor interaction probabilities from PNS_glial to ***minor*** CAFs and other ***major*** cell types. Line thickness and numbers indicate the total interaction strength (weight) between cell types; edge weights are comparable across cell types; circle sizes reflect cell numbers in each group. **G)** CellChat-derived centrality scores for the FGF and PDGF signalling pathways, highlighting dominant senders, receivers, mediators, and influencers (Jin et al. 2021) among ***minor*** CAF cell types and other ***major*** cell types. PNS_glial cells are among dominant senders; pnCAFs and other CAFs are dominant receivers. Scores are scaled row-wise. **H)** As in (F), showing interaction probabilities from pnCAFs to ***minor*** CAFs and other ***major*** cell types. **I)** As in (G), showing centrality scores for the COLLAGEN, LAMININ, and FN1 signalling pathways. CAFs are dominant senders; PNS_glial are dominant receivers.

The deconvolution analysis revealed distinct spatial localisation of Club, Luminal-like, and Luminal cells. Club cells, typically found in benign epithelium in the transitional prostatic zone (Henry et al. 2018) (P4-2, Supp. Fig. S16-S17), were also detected in the peripheral zone of adjacent-benign tissues and, though in lower numbers, in some cancer samples (Supp. Fig. S17-S18). Luminal-like and Luminal cells were dispersed throughout the benign cores (Supp. Fig S17). In cancer cores, Luminal-like cells were distinct from Luminal cells and localised to both histologically benign (P11-1, P10-1, P3-1, P15-2; Supp. Fig. S16-S17) and cancerous (P25, P13-2, Supp. Fig. S16, S18) regions. Overall, this suggests that both morphologically benign and malignant epithelium may exhibit signs of dedifferentiation, potentially losing Luminal features and acquiring Club-like properties.

Next, in our histopathological annotation of the samples, we distinguished between prostatic stroma, primarily comprising SMCs and fibroblasts within the prostate (Joseph et al. 2021), and extraprostatic stroma, rich in adipose tissue, blood vessels, and nerves surrounding the prostate. Although the anatomical separation between these stroma types can be ambiguous (Ayala et al. 1989; Sakr et al. 1996), extraprostatic stroma is an important structure, as malignant cells must penetrate it to invade the local organ bed.

Comparing CAF and SMC proportions between adjacent-benign and cancer cores in the Visium data reflected trends consistent with the scRNA-seq data. NPF-like cells were enriched in adjacent-benign cores, specifically in the prostatic stroma (Fig. 5B; Supp. Fig. S19 A, B). They also correlated with pSMCs on a per-spot level in both core types, suggesting close association and highlighting both cell types as major components of the prostatic stroma (Fig. 5C; Supp. Fig. S19C). However, the relationship between NPF-like cells with uniCAFs and whCAFs was unclear, with positive and negative correlations observed in both core types (Fig. 5C; Supp. Fig. S19C).

In 6 out of 10 samples, uniCAFs and whCAFs correlated with each other (Fig. 5C; Supp. Fig. S19C) and were enriched in cancer cores (Supp. Fig. S19C). Both cell types were predominantly found in the extraprostatic stroma (Fig. 5B; Supp. Fig. S19B), which explains the negative correlation we observed with NPF-like cells, confined to the prostatic stroma. uniCAFs, specifically, were enriched in vessel-associated and adipose regions (Fig. 5B). Importantly, they correlated with vSMCs and Pericytes (Fig. 5C; Supp. Fig. S19 B, C), suggesting their association with major and minor blood vessels, providing tissue-level validation of the signalling predictions identified by scRNA-seq. Additionally, uniCAFs positively correlated with pSMCs, particularly in the fibromuscular stroma at the prostate periphery (Fig. 5C; Supp. Fig. S19 B, C), highlighting their role as significant components of the outermost layer of the prostatic stroma (“capsule”). Moreover, this suggested that uniCAFs and whCAFs may not closely interact with epithelial cells and might influence localised PCa biology more indirectly through roles in extraprostatic extension or by affecting immune cell trafficking.

pnCAFs were enriched in nerve regions (Fig. 5B) and were strongly correlated with Glial subtypes, such as myelinating Schwann and Satellite cells, particularly in nerve-enriched samples (Fig. 5C, D; Supp. Fig. S19C). To ensure neurons in the Visium data, which were absent from the scRNA-seq reference, were not misidentified as pnCAFs in the deconvolution analysis, we leveraged Cepo (Kim et al. 2021) to generate a ranked list of cell type-specific genes (ST_6) and identify stably expressed pnCAF markers using the scRNA-seq data. The pnCAF list was refined by excluding genes expressed in Glial cells (Supp. Fig. S20A), resulting in a 9-gene pnCAF-specific signature (ST_7). All genes are known fibroblast markers and none are classical neuron markers, though some are co-expressed at varying levels by peripheral nervous system cells (Supp. Fig. S20B). Rescoring each Visium spot with these curated gene signatures revealed their enrichment in nerve-enriched samples, specifically in regions morphologically annotated as nerves (Fig. 5E; Supp. Fig. S20 C, D). We also used published sub-cellular resolution Xenium spatial data to demonstrate that distinct cell types within nerve bundles are enriched in pnCAF and PNS_glial signatures (Supp. Fig. S21, ST_9), further supporting the association of pnCAFs with nerves.

This association was noteworthy, as it might have implications for regulating perineural invasion, which occurs in ∼50% of patients and is considered a preferred metastatic pathway for PCa cells (Zareba et al. 2017; Kraus et al. 2019). We extended our CellChat L-R analysis to explore interactions between Glial cells and pnCAFs, aiming to understand whether pnCAFs recruit Glial cells - and potentially neurons - or vice versa. Glial cells showed the strongest autocrine interactions (Fig. 5F), and represented a source of PDGF and FGF signalling to CAFs, including pnCAFs (Fig. 5G). Conversely, pnCAFs exhibited the strongest putative outgoing interactions with Glial cells, indicated by the weight of aggregated interactions (Fig. 5H). Glial cells primarily responded to ECM signalling from CAFs, including pnCAFs, through the collagen, laminin, and fibronectin pathways (Fig. 5I). Notably, they exhibited high influencer and mediator scores (Jin et al. 2021) for these pathways (Fig. 5I), suggesting they may serve as gatekeepers for cell-cell communication between CAFs and other cell types. CAF ligands, such as Collagen VI (Supp. Fig. S22 A, B), and other ECM components are known to promote differentiation of Glial cells, neuronal myelination, and coordination of nerve regeneration post injury (Vitale et al. 2001; Lam et al. 2019). This underscores the potential role of pnCAFs as regulators of Glial cell and potentially neuronal function.

## Discussion

Our cohort from predominantly intermediate-risk patients is representative of the natural occurrence of prostate cancer and constitutes the largest scRNA-seq PCa cohort to date. We provide the most comprehensive cellular annotation of PCa, identifying 11 ***major*** cell types, 50 ***minor*** subtypes, and numerous cell state classifications. Integrated with published bulk gene expression studies, this detailed reference atlas will serve as a powerful resource for exploring PCa as a cellular system using deconvolution methods like ecotyping (Wu, Al-Eryani, et al. 2021; Steen et al. 2023). This is particularly relevant given the significant interpatient heterogeneity we observed in cellular composition, which may have both biological and clinical implications.

Our scRNA-seq analyses highlighted an abundance of Club cells and mapped epithelial cells across a malignant spectrum, enabling precise investigation of baseline AR signalling across various functional states. We also provide the most extensive annotation of CAFs in PCa and identified a novel CAF population closely associated with the peripheral nervous system. Using Visium spatial transcriptomics, we uncovered distinct spatial distributions of epithelial phenotypes and CAF phenotypes within the PCa TME.

We identified a novel antigen-presenting phenotype (*HLA-DRA* (MHC II) */CD74*+) in Club cells (Blackburn et al. 2023). Depletion of MHC II^high^ epithelial cells has been linked to dysregulated barrier immunity in the lung (Shenoy et al. 2021), suggesting a role in mediating and maintaining local immune responses. Although rare in the peri-urethral regions of the normal prostate (Henry et al. 2018) and in other PCa scRNA-seq studies (Song et al. 2022), Club cells were the second most prevalent epithelial subtype in our dataset. Their signatures were detected in the transitional zone of the prostate, as previously reported (Henry et al. 2018), but also in association with benign epithelium in the peripheral zone of both adjacent-benign and cancer cores.

Additionally, Luminal-like cells, potentially representing an intermediate state between Luminal and Club cells, were detected in both adjacent-benign and cancerous cores, associated with both morphologically benign and malignant epithelium in the spatial transcriptomics data. Our results overall suggest that Luminal epithelium, whether benign or malignant, may lose its typical features and acquire Club-like characteristics.

Supporting these findings, an expansion of Club cells (PIGR/LCN2/MMP7/CP+) has been observed by immunohistochemistry in atrophic lesions in the prostatic peripheral zone, including prostatic inflammatory atrophy, a precursor to PCa and prostatic intraepithelial neoplasia (Huang et al. 2023). These cells were localised toward the glandular lumen, suggesting that Luminal to Club cell differentiation may occur under inflammatory conditions (Huang et al. 2023).

Genomic alterations such as chr8p loss (e.g., *NKX3-1*) and chr8q gain (e.g., *MYC*) have been identified in atrophic regions (Yildiz-Sezer et al. 2006). In our study, two of three Club cell states (“SAA1+/c15”, “CCL20+/c16”) were patient-enriched, possibly indicating patient-specific genomic profiles. Although clonal relationships between these cells and their malignant counterparts could not be assessed due to technical constraints, our findings suggest that the abundance of Club cells and presence of Luminal-like cells may be linked to ongoing dedifferentiation in a tumor-promoting microenvironment, consistent with a transcriptional field cancerisation effect.

We observed widespread CNV changes across all epithelial lineages, consistent with the growing recognition of somatic events in functionally and histologically normal tissues (Yizhak et al. 2019; Grossmann et al. 2021). Whether altered benign cells are precursors to PCa and clonally related to malignant cells remains unclear. These molecular changes may represent pre-cancerous lesions or alternatively benign histopathologies that are not on the path to malignant progression. Higher-resolution methods, ideally incorporating direct dual measurements of genomic copy number plus gene expression are necessary to validate the presence or absence of PCa-associated driver events and define the progression of disease.

Our analysis of AR signalling dynamics across Luminal functional states revealed that altered benign MSMB+ cells, despite the presence of CNVs, functionally resemble their normal counterparts with respect to AR signalling, with both states enriched in a normal AR repertoire. The MSMB+ states exhibited higher levels of androgen response compared to MSMB^low^ malignant cells, which were enriched in PCa-specific AR activity. This indicated that the loss of the MSMB+ phenotype is associated with gradual suppression of AR differentiation and secretory programs and a shift towards oncogenic AR signalling.

Comparisons between altered benign MSMB+ cells and MSMB^low^ malignant cells showed that downregulation of the canonical androgen response was associated with increased MYC activity, consistent with recent findings that MYC upregulation is a universal feature in PCa malignant versus normal luminal cells (Graham et al. 2024). We found that altered benign and malignant MSMB+ cells also exhibited a higher canonical androgen response than MSMB^low^ malignant cells, suggesting that the loss of the MSMB+ phenotype is a critical checkpoint in the transition to malignancy. This underscores the need to identify regulatory pathways driving full malignant transformation and AR reprogramming, which may offer opportunities for therapeutic prevention of PCa progression or inducing differentiation of the prostatic epithelium.

Although MYC is overexpressed in >80% of localised PCa (Koh et al. 2010), its role and timing in the transcriptional reprogramming of the prostate remains unclear. Consistent with our findings, MYC over-expression in PCa cell lines antagonises AR activity and androgen-induced gene expression *in vitro (Barfeld et al. 2017)*, including *KLK3*, which encodes the clinically relevant prostate specific antigen (PSA) protein. *In vivo,* MYC overexpression dampens the canonical AR transcriptional program by pausing RNA polymerase II at promoters of AR-dependent genes, including those in the hallmark androgen response signature (Qiu et al. 2022). This suggests that the downregulation of androgen response in MSMB^low^ malignant cells could result from MYC antagonism. Clinically, divergent MYC and AR activity is relevant, as a MYC high/AR low phenotype in malignant cells is associated with faster biochemical recurrence (Qiu et al. 2022). Future studies should investigate the heterogeneity of these TFs at the intrapatient level. We reported four Fibroblast phenotypes: uniCAFs, NPF_like, whCAFs, and a novel pnCAF phenotype, along with distinguishing features for the SMC lineage (pSMCs, vSMCs, and Pericytes). Analysis of our scRNA-seq and Visium datasets revealed diverse distributions of CAFs in the stroma, linked to different anatomical niches. Notably, the NPF-like population, characterised by *PTGS2* (COX2), and *PTGDS* (PGD2) expression, was the only population found interstitially, near epithelial cells. It was also the only cell type in our dataset significantly enriched in adjacent-benign cores, suggesting a resting or tumour-suppressive phenotype. Stromally-derived prostaglandin metabolism is known to maintain prostate epithelial homeostasis (Kim et al. 2021). Urinary PGD2 concentration is significantly higher in non-inflamed PCa compared to inflamed PCa (Bellei et al. 2023). Since inflammation in PCa has been associated with adverse prognosis and clinicopathological features (Sciarra et al. 2016), this implies a protective immunomodulatory role for *PTGDS+* NPF-like cells in PCa. The identification of NPF-like cells as mediators of prostaglandin metabolism is novel, highlighting their potential role in maintaining homeostasis and shaping the immune phenotype (Kickler et al. 2012; Knoblich et al. 2018) of PCa.

We report a novel, specialised pnCAF phenotype (*DPT^low^ and PI16^high^*), enriched in axonal and neuronal projection guidance processes. Spatial deconvolution analyses and correlation of cell type-specific signatures indicated frequent co-localisation of pnCAFs and Glial cells. pnCAFs may respond to FGF/PDGF signalling from Glial cells, while Glial cells may respond to ECM-associated signalling from pnCAFs, suggesting there may be a reciprocal interaction between pnCAFs with Glial cells and/or neurons in the recruitment or maintenance of each other’s states. Whether these interactions are homeostatic roles or involved in perineural invasion (PNI) associated with PCa remains to be determined.

PNI is considered a preferred metastatic pathway for PCa and a poor prognostic factor in radical prostatectomy pathology (Zareba et al. 2017). Investigating pnCAF interactions with malignant cells in PNI-positive tumours and exploring strategies to disrupt these interactions could reveal new avenues to impair PCa growth and metastatic spread (Magnon et al. 2013; Xue et al. 2023). Furthermore, pnCAFs and their interactions with Glial cells and/or neurons may be important in cancer-associated neuropathic pain (Singhmar et al. 2020; Shi et al. 2023). Overall, these analyses lay the groundwork for further exploration and validation of the identified cell states using higher-sensitivity methods and functional characterisation.

Crucially, even when considering our cohort alongside recently published PCa scRNA-seq datasets (S. Chen et al. 2021; Tuong et al. 2021; Masetti et al. 2022; Siefert et al. 2021; Song et al. 2022; Wong et al. 2022; Graham et al. 2024), the total number of patients profiled is only ∼70 patients. In future research, studying a larger cohort of patients will be essential. Specifically, two key studies are needed: (1) paired transcriptome and copy number analysis to better characterise epithelial diversity and clonal relationships, and (2) high-resolution (single-cell or sub-cellular) spatial analysis to definitively localise stromal, epithelial, and immune populations.

In summary, our findings reveal distinct molecular features and spatial distributions of key cell populations and offer a critical reference for exploring PCa pathobiology.

## Methods

### Patient material and ethics approval

Patient material collection was conducted with approval from the St Vincent’s Hospital Human Research Ethics Committee, under protocol SVH 12/231. Written consent was obtained from all participants, and their clinical data was de-identified.

### Risk stratification

For analytical purposes, patients were categorised into three risk groups (Intermediate Favourable, Intermediate Unfavourable, and High) based on pre-operative PSA serum levels, Gleason grade, and clinical stage. Clinical metadata, including patient age, are available in ST_9.

### Tissue Collection and Storage

Two 5 mm core biopsies were obtained from 70 hormone therapy-naïve radical prostatectomy specimens at St Vincent’s Hospital, targeting either the largest (index) tumour focus or two distinct tumour foci based on pre-surgery MRI assessments. Cores were placed into a collection medium (RPMI 1640, no phenol red (Gibco, cat# 11835030) and 10% FBS (Hyclone, cat# SFBS-AU) and stored on ice or at 4℃. All samples were processed within 2-4 hours. Each core was split into a scRNA-seq and a Formalin-Fixed Paraffin-Embedded (FFPE) section. The scRNA-seq section was diced (0.5-1 mm³), cryopreserved (95% FBS, 5% DMSO (Sigma-Aldrich, cat# 20-139)), and stored in liquid nitrogen. The FFPE section was fixed in 10% neutral buffered formalin for 24 hours, dehydrated in 70% v/v ethanol, and processed for H&E staining using 4 μm sections.

H&E slides were imaged on the Aperio CS2 Digital Pathology Slide Scanner (Leica Biosystems). An expert prostate pathologist (Prof James Kench, Royal Prince Alfred

Hospital) assessed the images to confirm the presence of tumour and estimate epithelial, stromal, and immune cellular composition. Samples were included in the study cohort only if a tumour focus was sampled, resulting in either two tumour cores or one tumour and one adjacent-benign core per patient (ST_1). Due to insufficient tumour representation, 66% (46/70) of patients were excluded. No statistical method was used to predetermine sample size.

### Buffy Coat Isolation and Storage

Whole blood samples were collected in EDTA-coated Vacutainers (BD, cat# 367863). Buffy coat was isolated by centrifugation at 300xg for 10 minutes at 4°C. 100-200 μL of buffy coat was snap-frozen and stored at -80°C. The remaining buffy coat was mixed with cryopreservation medium (40% RPMI 1640, 50% FBS, 10% DMSO), and frozen at a rate of 1℃ per minute using a Mr. Frosty Freezing Container (Thermo Scientific, cat# 5100-0001) to -80°C, before being transferred to liquid nitrogen within 1-2 weeks for long-term storage.

### Tissue processing for scRNA-seq

Samples were transferred from liquid nitrogen to -80°C the day before processing. After thawing in a 37°C water bath for 2 minutes, they were transferred to a 15mL LoBind tube (Eppendorf, cat #0030122216) and washed twice with cold DPBS (Gibco, cat# 14190144). Floating cells were pelleted by brief centrifugation (350xg; 30 seconds; 4°C) between washes. The supernatant was removed, and samples were transferred to a 5mL tube (Eppendorf, cat# 0030122348) for multiplexing before dissociation, following a pre-designed strategy (ST_2).

Tissue dissociation followed a modified protocol of the Human Tumour Dissociation Kit (Miltenyi Biotec, cat# 130-095-929). Tissues were finely dissected to a “slurry”, mixed with tumour dissociation enzyme mix (ST_10), sealed with Parafilm, and incubated in a heated shaking incubator (37°C; ∼180rpm) for 60-80 minutes. Dissociation progress was monitored every 15 minutes until only a few tissue chunks remained. Cell suspensions were filtered through a 100μm MACS strainer (Miltenyi Biotec, #130-098-463) into chilled 15mL LoBind tubes with neutralisation buffer (RPMI 1640, 50% FBS). Remaining tissue was further dissociated with 2mL of pre-warmed TrypLE Express (ThermoFisher Scientific, cat# 12563029) for 3 minutes at 37°C, then gently triturated. Cell suspensions were combined, centrifuged (350xg; 7 minutes; 4°C), and filtered again if necessary, through a 70μm MACS strainer (Miltenyi Biotec, cat# 130-098-462).

After aspirating the supernatant, the cell pellet was gently resuspended in 100μL of 10x capture buffer (DPBS, 10% FBS). Viability, cell count, and purity were assessed with a haemocytometer (RRID:SCR_025846) using Trypan Blue (0.4% solution, Gibco, cat# 15250061). Samples with <70% viability and >100K cells were enriched for viable cells using the EasySep Dead Cell Removal (Annexin V) Kit (STEMCELL Technologies, cat# 17899). The live cell-enriched population was pelleted (350xg; 5 minutes; 4°C) and resuspended in 10x capture buffer for loading on the 10x Chromium platform. A detailed list of reagents and materials is provided in ST_11.

### Single-cell RNA-sequencing using 10x Chromium

ScRNA-seq was conducted using the Chromium Single Cell 3’ Gene Expression v3 Kit (10x Genomics) on the 10x Chromium Platform, per the manufacturer’s protocol. Each reaction (ST_2) targeted 18,000 cells. Libraries were sequenced on an S4 flow cell on the NovaSeq 6000 System (Illumina; RRID:SCR_016387), with pair-end sequencing and single indexing (Read 1 – 28 cycles; i7 index – 8 cycles; Read 2 – 91 cycles).

### DNA extraction from buffy coat and genotype profiling

DNA was extracted from snap-frozen buffy coat using the QIAamp DNA Blood Mini Kit (Qiagen, cat# 51104) following the manufacturer’s protocol. DNA concentration was quantified with the dsDNA High Sensitivity assay on a Qubit® 3.0 Fluorometer (ThermoFisher Scientific; RRID:SCR_020311). Purity was assessed by measuring absorbance at 260 nm and 280 nm using a NanoDrop 2000 Spectrophotometer (ThermoFisher Scientific; RRID:SCR_018042). For genotype profiling, ∼60μL of DNA at 10ng/μL was aliquoted into a 96-well plate and processed on the Applied Biosystems Axiom v2.0 UK Biobank Array (ThermoFisher Scientific, cat# 902502). All samples passed QC thresholds (Axiom Analysis Suite, v5.0.1.38), and the resulting. CEL files were used for genotype (SNP)-based demultiplexing.

### Tissue processing for Visium spatial transcriptomics

The FFPE blocks were cut into 4 μm sections and processed using the Visium Spatial Gene Expression Kit (FFPE 10x Genomics; RRID:SCR_023571), following the manufacturer’s protocol. Six samples were processed using FFPE V1, and four samples were processed using FFPE V2 (ST_5).

Samples processed with FFPE V1 were rehydrated using xylene and ethanol gradients. Rehydrated tissue sections were H&E stained and imaged using a Leica DM6000 microscope (Leica Microsystems) at 20x magnification. The imaged tissues were then processed for antigen retrieval, followed by probe hybridisation, and ligation. Ligation products were released from the tissues and captured by poly-dT probes on the surface of the Visium Spatial Gene Expression slides.

Samples processed with FFPE V2 were sectioned onto a SuperfrostPlus Gold Adhesion slide (Epredia, cat# K5800AMNZ72), stained with an optimised Visium H&E staining protocol (ST_12), and imaged using a NanoZoomer S210 microscope (Hamamatsu Photonics, cat# C13239-01), under a 40x lens magnification. The samples were then transferred to a 10x Visium slide using the CytAssist platform (10x Genomics; RRID:SCR_024570) and processed according to the manufacturer’s protocol.

Libraries were sequenced on the NovaSeq 6000 System (Illumina), with pair-end sequencing and dual indexing (Read 1 – 28 cycles; i7 index – 10 cycles; i5 index – 19 cycles; Read 2 – 50 cycles). At least 40,000 reads were obtained on average for each Visium spot.

### Data mapping and pre-processing

#### scRNA-seq data pre-processing

Cell Ranger (10x Genomics, v3.1.0; RRID:SCR_017344) was used for demultiplexing raw BCL files, genome alignment (GRCh38), filtering with EmptyDrops, and counting barcodes and Unique Molecular Identifiers (UMI). scRNA-seq data normalisation, dimensionality reduction, and clustering were performed with *Seurat* (v3.2.2; RRID:SCR_016341). Samples with <50 cells, cells with <250 genes (nFeature_RNA), <500 UMIs (nCount_RNA), and percent mitochondrial reads (pt.mito) >20% were filtered out.

#### Visium data pre-processing

Space Ranger (10x Genomics, v1.3.1 for FFPE V1; v3.0.1 for FFPE V2; RRID:SCR_025848) was used for demultiplexing raw BCL files, genome alignment (10x Genomics, refdata-gex-GRCh38-2020-A), and counting barcodes and UMIs. The analysis used the Visium Human Transcriptome Probe Set (10x Genomics, v2.0 GRCh38-2020-A).

Visium data was processed with *scanpy* (v1.9.1; RRID:SCR_018139) and *anndata* (v0.8.0; RRID:SCR_018209). Deduplicated count matrices were used to construct sample-specific objects. Spots with <1 UMI were excluded. Dimensionality reduction was performed with 50 principal components, followed by Leiden clustering. Histology of Visium spots was annotated by a prostate pathologist (Dr Alison Potter), blinded to gene expression clusters, using *Loupe* (10x Genomics, v7.0.1; RRID:SCR_018555). Spots outside tissue areas were manually annotated as “Exclude”.

### SNP-based demultiplexing

Cell Ranger *possorted_genome_bam.bam* files were used for SNP-based clustering of individual cells with Souporcell (SNP-based demultiplexing; built from commit “790b281dbe”) (Heaton et al. 2020). Souporcell-derived SNP profiles were matched to patient-matched genotypes (.CEL files) and assigned to patient IDs using VireoSNP (v0.5.6) (Huang et al. 2019). Default parameters were used for all analyses.

### Cell type annotation

#### Preliminary cell type annotation

Preliminary cell type annotation was performed using SingleCellNet (SCN; v0.1.0) (Tan & Cahan 2019). The SCN classifier, trained on our breast cancer scRNA-seq dataset (Wu, Al-Eryani, et al. 2021), annotated major cell types (*Epithelial, CAFs, SMCs, Endothelial, Myeloid, T-cells, B-cells, Plasma cells, rand*), reflecting similarities between the human breast and prostate cellular compositions (Henry et al. 2018; Kumar et al. 2023). This annotation guided cell type reclustering and the specification of query/reference cells for CNV analysis. It was then reviewed and manually refined due to the absence of myoepithelial cells in the prostate (Henry et al. 2018), pSMCs in the breast (Kumar et al. 2023), and the presence of a ‘*rand*’ cluster in the PCa scRNA-seq data (Supp. Fig. S23).

#### Cell type–specific reclustering analyses

Barcodes for major cell types (from SCN-annotation) were subset for reprocessing and reclustering using Seurat (v3.2.2). The data was log-normalised (’NormalizeData()’), and the top 2,000 variable features were selected from each cell type-specific object (’FindVariableFeatures()’). The data was scaled using linear transformation (’ScaleData()’), and mitochondrial contamination (“pt.mito”) was regressed out. Principal Component Analysis (PCA) was conducted on the scaled data using 100 principal components (PCs). Significant PCs were identified (’JackStraw()’ and ‘ScoreJackStraw()’, overall.p.value < 0.01), and used to construct local neighbourhoods (’FindNeighbors()’) and cluster cells at various resolutions (’FindClusters(algorithm=1, resolution = c(seq(0.1, 0.8, by=0.1))’), followed by UMAP visualisation (’RunUMAP()’). Plasma and Unassigned cells were not processed separately due to lower cell numbers.

Data from different patients were concatenated and processed without batch correction or integration, as performed previously (Wu, Al-Eryani, et al. 2021). For individual patient analysis, epithelial cells were subset by patient ID and processed with clustering resolution set to “RNA_snn_res.0.5” due to lower cell counts.

#### Differential gene expression analysis (DGEA)

DGEA was performed for each cell type-specific object using the MAST method (Finak et al. 2015) at the default clustering resolution (“RNA_snn_res.0.8”). For B-cells, the resolution was set to “RNA_snn_res.0.3” due to fewer cells. Upregulated differentially expressed genes (DEGs) were reported with *min.pct = 0.25* and *logfc.threshold = 0.25*, unless otherwise specified.

#### Manual refinement

After DGEA, QC metrics (“nFeature_RNA”, “nCount_RNA”, “pt.mito”) and the top 50 DEGs for each cluster were inspected. Clusters with low-quality cells (e.g., “nFeature_RNA” < 1000 and concurrent high median “pt.mito”) or heterotypic doublets (e.g., DEGs included both *CD3D* and *PDGFRA*) were labelled as “Low_QC” and “Doublets” cells, respectively, and filtered out. Cells misannotated by SCN (e.g., *Low QC* cells misannotated as *B-cells, PNS_glial* cells annotated as *rand*) were re-assigned to their correct lineages under “cell type major” (***major***; Supp. Fig. S23) based on DEGs at “RNA_snn_res.0.8” and identifying gene markers (listed in ST_3).

Cellular taxonomy was further defined at two additional tiers, as done in (Wu, Al-Eryani, et al. 2021): “cell type minor” (***minor***) and “cell type subset” (***subset***), The ***minor*** tier was determined by combining lower-resolution clustering (e.g., “RNA_snn_res.0.2”) with marker gene expression, whereas the ***subset*** tier reflected cell grouping at the default clustering resolution (“RNA_snn_res.0.8”). Nomenclature examples are provided in ST_4. Cell type-specific objects were merged and reprocessed as a single object (‘merged object’), as outlined above.

### Estimating Copy Number Variation (CNV) in scRNA-seq data

#### CopyKAT

CNV signal and ploidy of individual cells were estimated using *CopyKat* (Gao et al. 2021) on filtered counts matrices of individual samples. Non-epithelial cells (*Endothelial, T-cells, SMCs, B-cells, Myeloid, Plasma cells*) were used as CNV-free references, and epithelial cells were the queries. CNVs were inferred using ‘copykat(id.type=”S”, ngene.chr=5, win.size=25, KS.cut=0.1, distance=”euclidean”)’, and cells were classified (‘aneuploid/CNV^pos^’ vs. ‘diploid/CNV^neg^’) natively by *CopyKAT*.

The “CNA_value” (Fig. 3D) was computed from the average squared CNV signal at each genomic locus, within -1 to +1 range. An average CNV profile was generated by selecting the top 5% of cells with the highest genomic instability scores per tumour, and each cell’s correlation estimate (“cor.estimate”) was derived by correlating with this profile, as done previously (Wu, Al-Eryani, et al. 2021).

CNV heatmaps in the supplementary figures combine *CopyKAT*-derived cell classification and InferCNV-derived CNV estimates. These heatmaps are organised by cellular barcode ID to enhance clarity, leveraging the denoising steps implemented within the inferCNV pipeline.

#### inferCNV

*inferCNV* (Tirosh et al. 2016) was run on individual samples containing ≥ 50 Epithelial cells. The CNV signal for individual cells was estimated using a 100-gene sliding window. Non-epithelial cells (excluding ‘CAFs’ and ‘rand’) were used as reference profiles for CNV-free cells, with epithelial cells as queries.

### Assessing enrichment of gene signatures

Signature activity was evaluated using AUCell (v1.12.0) (Aibar et al. 2017) on the log-normalised scRNA-seq counts matrix. Unless otherwise indicated, AUCell was run using default parameters, assessing whether a subset of the input gene signature was enriched among the top 5% of expressed genes in each cell. AUC threshold values were obtained using ‘getThresholdSelected()’ to classify cells as “positive” or “negative” for each signature, where indicated.

For the identification of malignant cells, two PCa signatures “PCa_LIU_UP” and “PCa_WALLACE_UP” from MSigDB; RRID:SCR_016863) were applied to the merged object using AUCell, where ‘getThresholdSelected()’ was used to set and retrieve the AUC threshold values for each signature. Cells scoring above the assigned threshold AUC value of 0.15 for PCa_LIU_UP and 0.33 for PCa_WALLACE_UP was annotated as “PCa_liu_up_positive” and “PCa_wallace_up_positive”, respectively.

Where indicated, differences in means for AUCell activity scores were assessed using ‘stat_compare_means(method = “t.test”)’, while gene expression means were assessed using ‘stat_compare_means(method = “wilcox.test”)’.

The enrichment of pnCAF and Glial signatures in the Visium data was assessed using *scanpy* (v1.9.1), with ‘scanpy.tl.score_genes(adata = adata, ctrl_size=50, n_bins=25)’. Negative scores, indicating lower average expression compared to a random reference gene set, were filtered out. Spots scoring below 0.5 for the assessed gene sets were not included in the analysis. For Fig. 5E, adjacent spots with scores above the thresholds were grouped together if separated by no more than one spot. Spots separated by more than one spot were assigned to separate groups.

The enrichment of pnCAF, general CAF, and Glial cells signatures in the published Xenium dataset was assessed using Xenium Explorer (v3.0.0; RRID:SCR_025847) and visualised as summarised expression of cell type-specific transcripts, each represented by distinct colour codes.

### Gene ontology over-representation analysis (ORA)

DEGs were ranked by decreasing avg_logFC values, and the top *n* genes per cluster were selected for further functional annotation and over-representation analysis (ORA): 200 genes for Club cells, 200 for SMCs, and 150 for CAFs. Functional annotation and ORA were performed using *msigdbr* (v7.5.1; RRID:SCR_022870) for Club cells, and *clusterProfiler* (v4.2.0; RRID:SCR_016884) for SMCs and CAFs, with the ‘compareCluster(fun = “enrichGO”, OrgDb = “org.Hs.eg.db”, ont = “BP”)’.

### Gene-set enrichment analysis

DGEA was performed, with all thresholds removed, using ‘FindMarkers(test.use=”MAST”, logfc.threshold *= -Inf,* min.pct *= -Inf,* min.diff.pct*=-Inf*)’ on populations of interest (e.g., altered benign MSMB+ cells versus a malignant cluster). The DEGs were ranked by decreasing avg_logFC values. These, together with the hallmark signature library (“h.all.v7.2.symbols.gmt“), were used as input for gene-set enrichment analysis (GSEA) using ‘fgseaMultilevel()’ from the *fgsea* package (v1.20.0; RRID:SCR_020938). Gene sets with p.adj ≤ 0.05 were deemed significantly enriched.

### Cellular proportion analysis

Differences in cellular proportions between core types (adjacent-benign vs cancer) were evaluated using the ‘propeller()’ function from the *speckle* package (Phipson et al. 2022). The ‘propeller()’ function accounts for cell counts and calculates proportions directly from a Seurat object, with the option to apply either arcsin square root (transform = “asin”) or logit transformation (transform = “logit”) to the resulting proportions. Both transformations were applied, yielding consistently concordant results in terms of significance. Significant findings (FDR < 0.05) were reported as both “asin” and “logit” derived FDR values.

### Ligand-receptor analysis

Ligand-receptor (L-R) analysis was performed on log-normalised scRNA-seq data using CellChat (v2.1.2) (Jin et al. 2021), exploring all available options (“Secreted Signaling”, “Non-protein Signaling”, “Cell-Cell Contact”, and “ECM-Receptor”). Cells were grouped by ***major_temp*** annotations (CAFs at ***minor***, all other cell types at ***major*** level), and all analyses were performed with default parameters. Visualisation used native functions within CellChat. The signalling role of cell types was evaluated at the pathway level using ‘netAnalysis_signalingRole_network(slot.name = “netP”)’, and significant L-R interactions were identified for populations and pathways of interest using ‘extractEnrichedLR()’, with the significance threshold set at p.adj < 0.001 for visualisation.

### Spatial deconvolution using Cell2Location

Cell2Location (v0.1.3) (Kleshchevnikov et al. 2022) was used to estimate cell type numbers in each Visium spot. The model, trained on scRNA-seq data at ***major***, ***minor***, and ***major_temp*** levels, used the following parameters: ‘summary_name = ‘means’’ and ‘use_n_obs = 1000’. Cell type mapping employed default settings with these parameters: ‘N_cell_per_location = 10’, ‘detection_alpha = 20’, ‘batch_size = Nonè, ‘max_epochs = 30000’, ‘train_size = 1’, and ‘use_gpu = Truè. In Fig. 5B, cell types constituting <5% of a spot were filtered out.

A detailed list of pipelines and analytical packages is provided in ST_13.

### Identification of stably expressed genes using Cepo

The Seurat object was converted to a SingleCellExperiment object, and the count matrix was log2 normalised using ‘logNormCounts()’ from the *scater* (RRID:SCR_015954) package. Cepo (v1.10.2) (Kim et al. 2021) was applied to the normalised data, excluding genes expressed in <5% of any cell type. All other parameters were set to their default values. For each cell type, the top 100 genes with the highest cell type specificity scores were selected as cell type signatures.

### Accessing published data

Normal prostate scRNA-seq data (Henry et al. 2018) from GSE145843 were reprocessed as detailed in the scRNA-seq data pre-processing section. Xenium Prime 5K data for human PCa (FFPE) was downloaded from https://www.10xgenomics.com/datasets/xenium-prime-ffpe-human-prostate.

## Data availability

Raw scRNA-seq and genotyping data have been deposited with the European Genome-phenome Archive (EGA), hosted by the European Bioinformatics Institute and Centre for Genomic Regulation under accession number [XX]. Processed scRNA-seq, including cell type specific objects, and Visium data are available through the CellxGene Portal under accession number [XX]. A meta data dictionary for the columns in the processed objects is provided in ST_14.

## Code availability

Code is available on GitHub for the scRNA-seq [XX] and Visium [XX] analyses.

## Acknowledgements

This work was supported by a research grant from the St Vincent’s Clinic Foundation and supported by the generosity of J. McMurtrie, AM and D. McMurtrie, and the Petre Foundation. A.S. is the recipient of a Senior Leadership Fellowship from the NHMRC. E.A and S.vdL were supported by the Australian Government Research Training Program Scholarship. H.H. is supported by a CINSW ECF and the CanToo Foundation. L.A.S. is supported by a Principal Cancer Research Fellowship from the Cancer Council’s Beat Cancer project (PRF2919), the Flinder Foundations, and a Cancer Council NSW Project Grant (RG 23-04). The funders had no role in study design, data collection and analysis, decision to publish or preparation of the manuscript.

We thank the following people for providing their assistance with experiments: Anne-Maree Haynes, Anjali Balakrishnan, and Daniela Barreto for tissue and clinical metadata collection; Anaiis Zaratzian, Andrew de la Silva, and Michael Tayao from the Garvan Histopathology Facility for H&E staining, routine tissue processing, and Visium tissue preparation; Eric Lam, Mubarika Tyebally, Chia-Lin Chang, Joanna Warren, and Michael Geaghan from the Cellular Genomics and Data Science platforms for assistance with the scRNA-seq and Visium data generation and sequencing. We thank Kristijan Apostolov for the illustrations in Fig. 1A and Supp. Fig. 6.

