## Supplementary Figures 1-23 for "Profiling of epithelial functional states and fibroblast phenotypes in hormone therapy-naïve localised prostate cancer"

Supplementary Figure 1

A

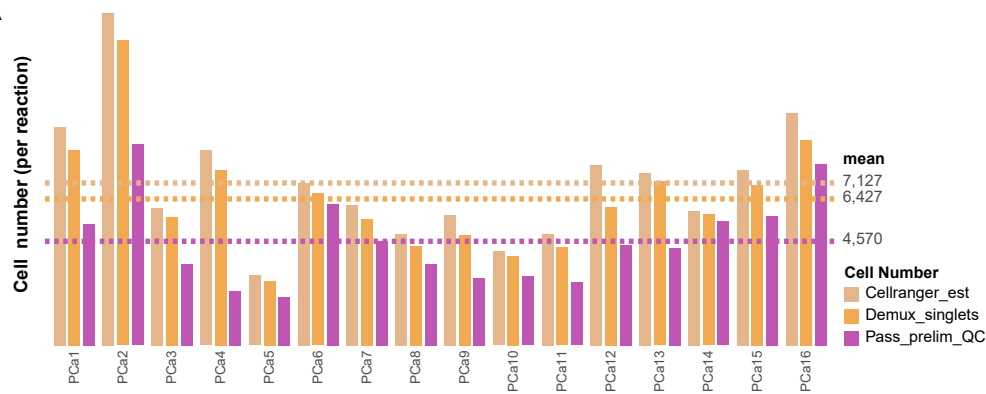

B

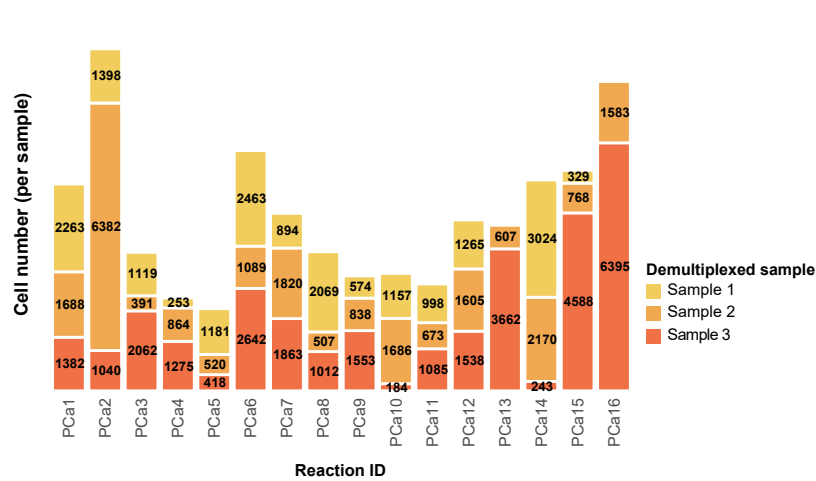

C

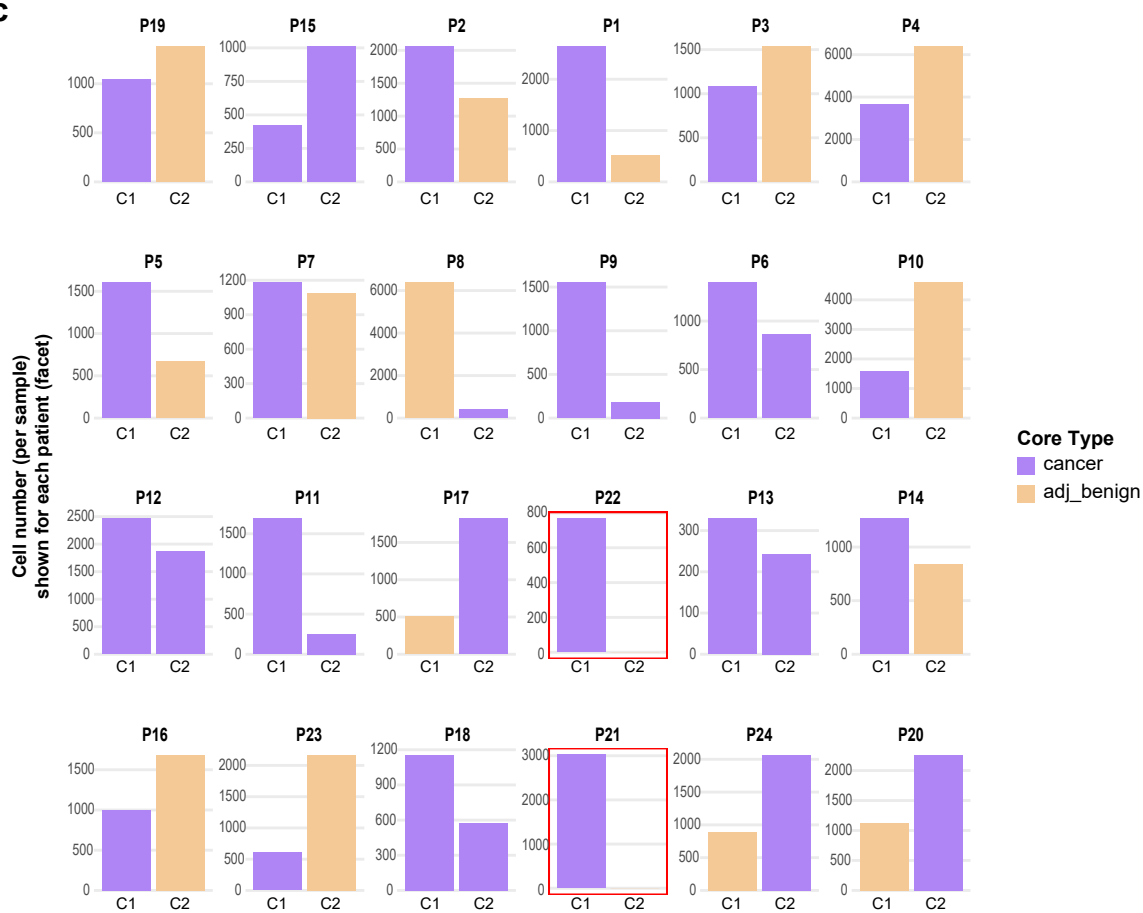

**Supplementary Figure 1:** Overview of cell numbers per reaction and sample

**A)** Cell counts per Reaction for CellRanger\_est, Demux\_singlets, and Passed\_prelim\_QC, with dashed lines indicating mean values (7,127; 6 427; 4,570, respectively). **B)** Cell counts per sample within each Reaction, coloured by the number of detected samples (sample 1, 2, and 3) after SNP-based demultiplexing. **C)** Side-by-side barplots of patient-matched samples (core 1 (C1) and core 2 (C2)), coloured by core type shown side-by-side; barplots are coloured by core type (cancer, adj-benign). Red rectangles highlight the two samples that were not detected post SNP-based demultiplexing (P21-2, P22-2).

Supplementary Figure 2

A

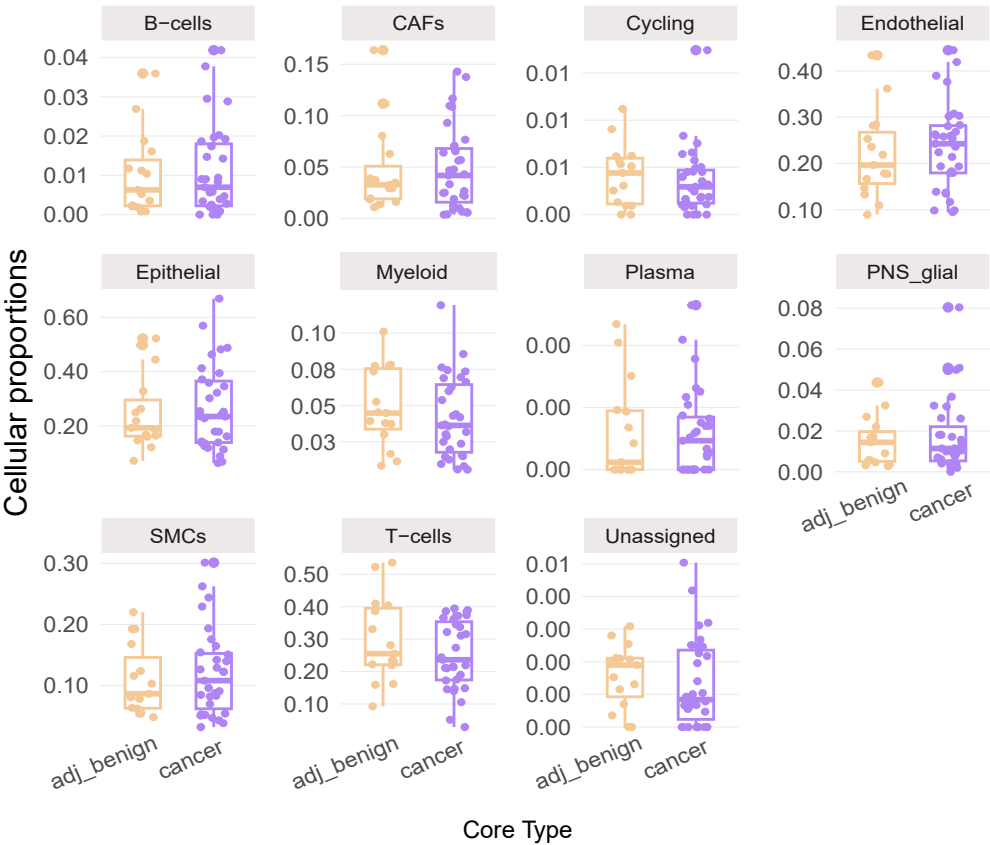

B

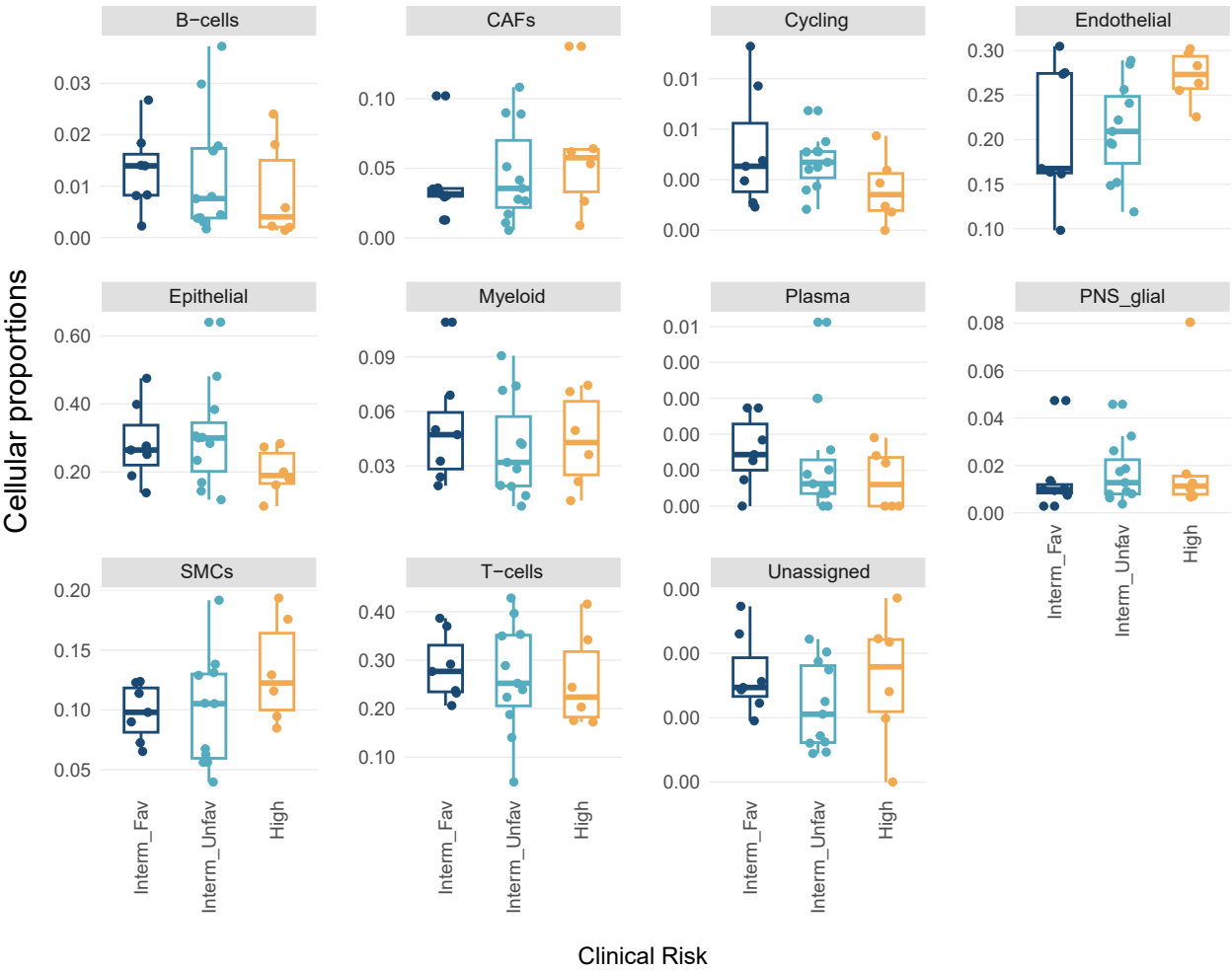

**Supplementary Figure 2: *major* cell type abundance**

**A) *major*** cell type abundance with respect to core types, with dots representing individual samples (n = 46). **B) *major*** cell type abundance across ISUP Clinical risk, with dots representing individual patients (n = 24). Statistical significance (FDR < 0.05) was not achieved (evaluated using speckle (Phipson et al. 2022)).

### Supplementary Figure 3

**A**

T-cells: *subset* annotation

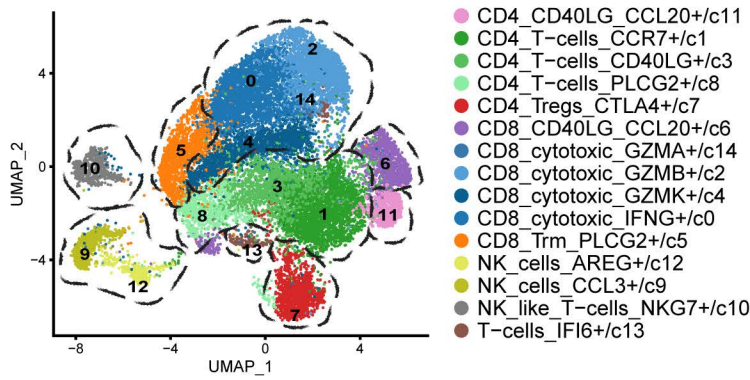

**B**

Cluster composition

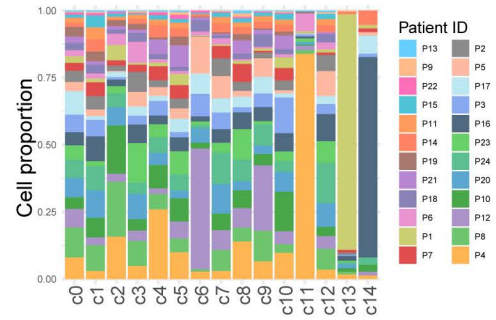

**C**

Top DEGs per *subset* category

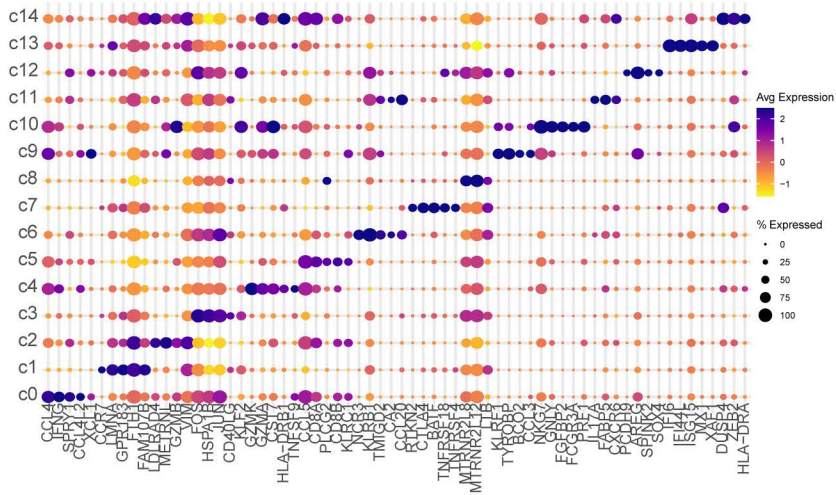

**D**

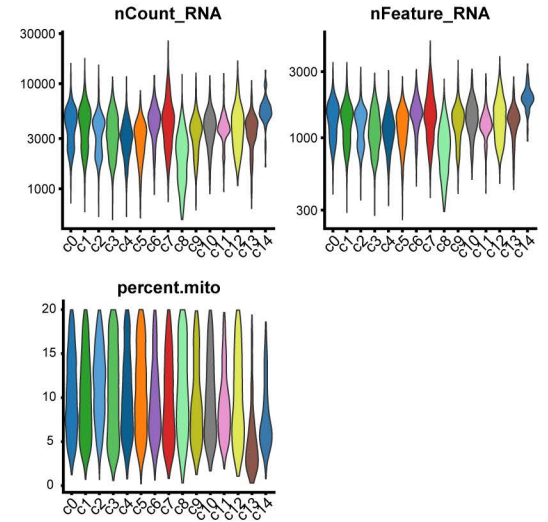

**E**

Myeloid: *subset* annotation

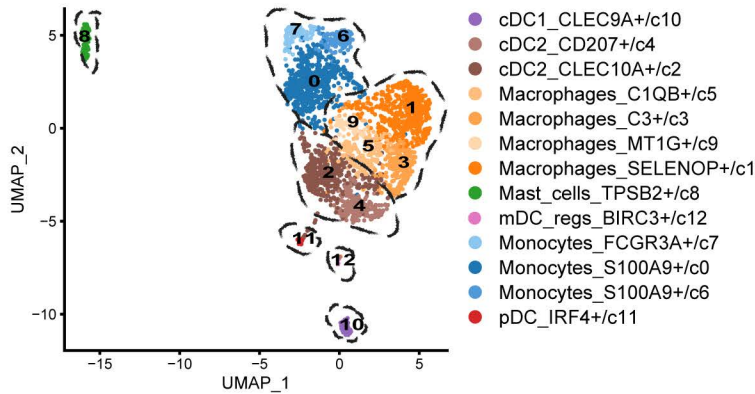

**F**

Cluster composition

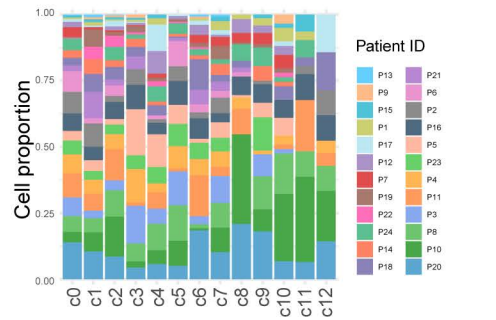

**G**

Top DEGs per *subset* category

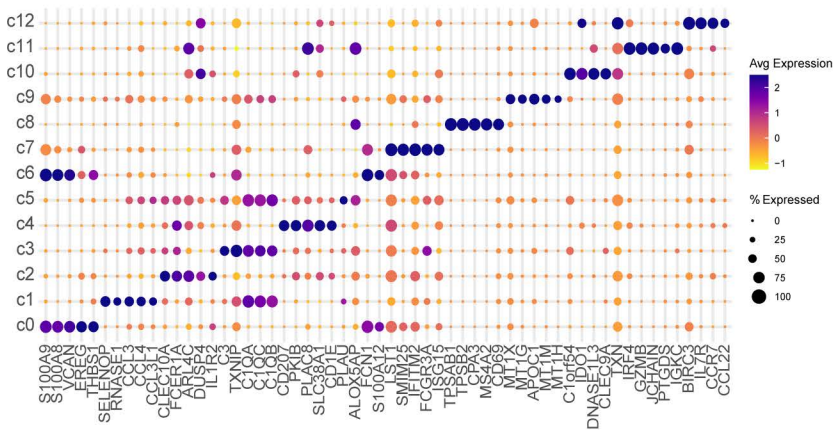

**H**

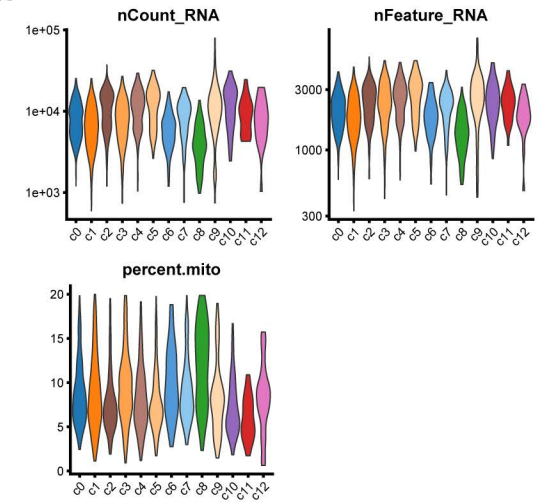

**Supplementary Figure 3:** Overview of T-cells and Myeloid cells in the PCa TME.

**A-D:** T-cells; **E-H:** Myeloid cells.

**A, E)** UMAP visualisation of **subset** annotation; dashed circles indicate **minor** lineages. **B, F)** **subset** cluster composition, filled by 24 unique patients (P1-P24). **C, G)** Average expression of the top 5 differentially expressed genes per **subset** category. **D, H)** Quality Control metrics ("nCount\_RNA", "nFeature\_RNA", "percent.mito"), with cells grouped by **subset** annotation. y-axis for "nCount\_RNA" and "nFeature\_RNA" shows log values.

### Supplementary Figure 4

**A**

Endothelial: *subset* annotation

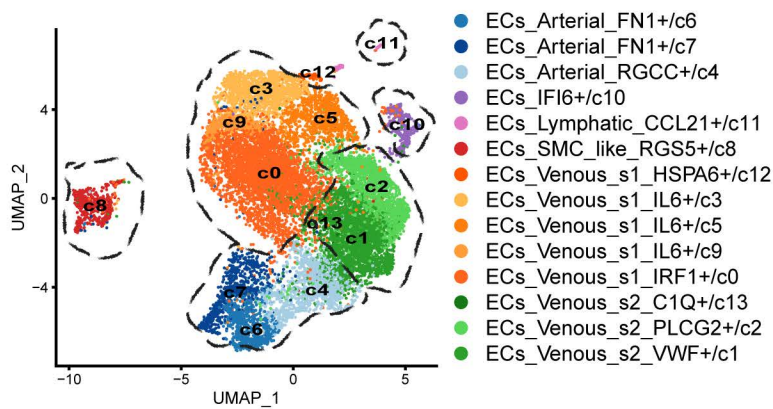

**B**

Cluster composition

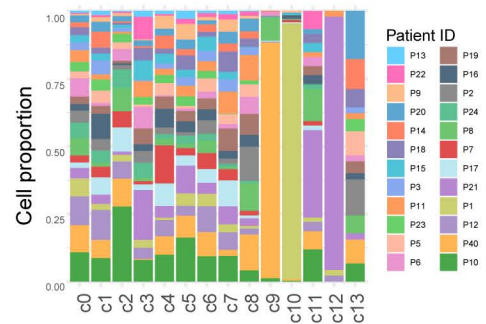

**C**

Top DEGs per *subset* category

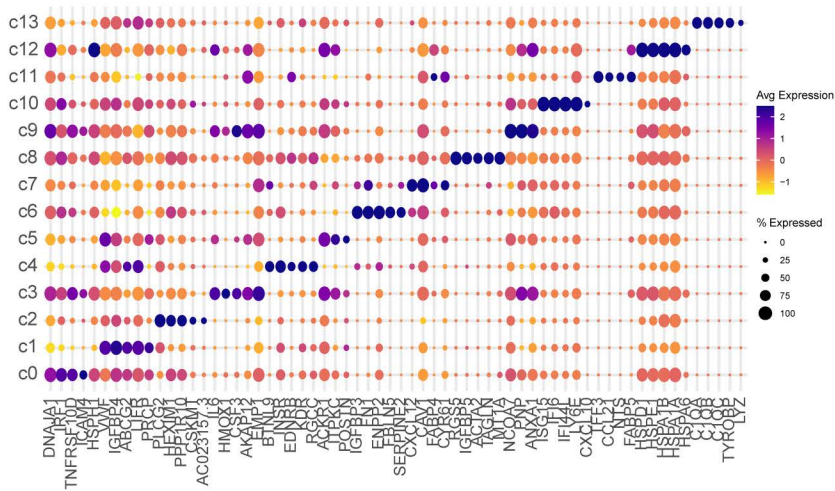

**D**

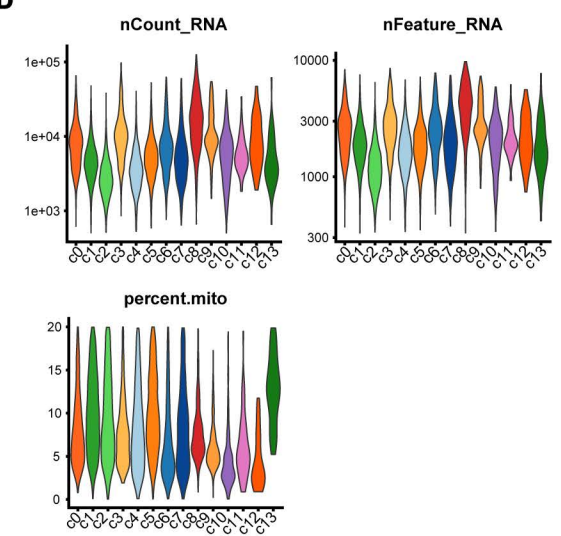

**E**

PNS\_glia: *subset* annotation

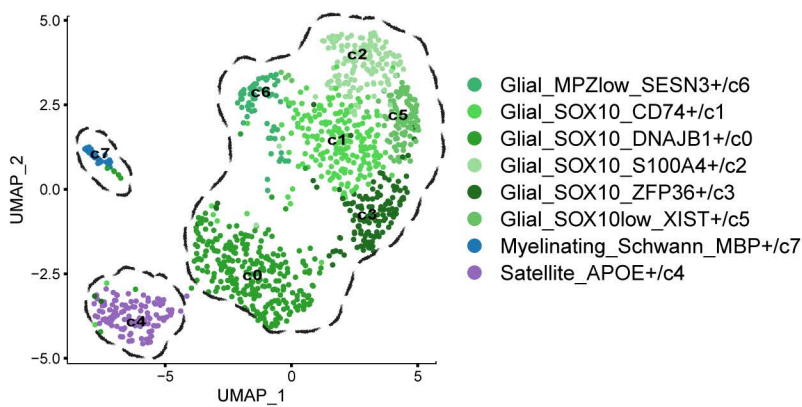

**F**

Cluster composition

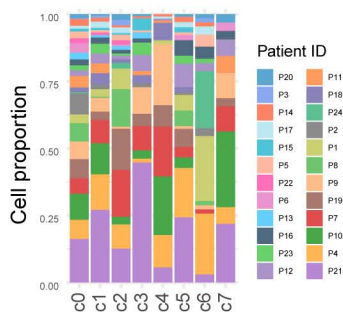

**G**

Top DEGs per *subset* category

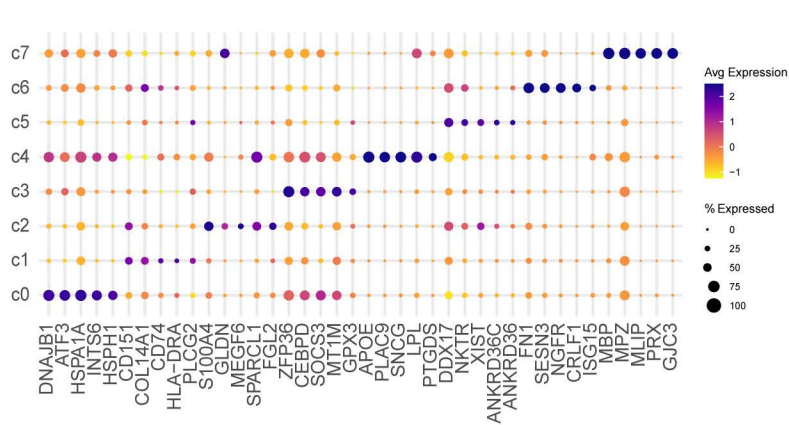

**H**

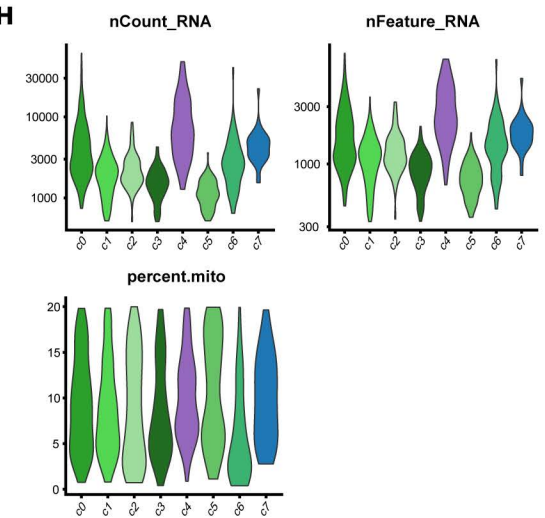

**Supplementary Figure 4:** Overview of Endothelial and PNS\_glial cells in the PCa TME.

**A-D:** Endothelial cells; **E-H:** PNS\_glial cells.

**A, E)** UMAP visualisation of **subset** annotation; dashed circles indicate **minor** lineages. **B, F)** **subset** cluster composition, filled by 24 unique patients (P1-P24). **C, G)** Average expression of the top 5 differentially expressed genes per **subset** category. **D, H)** Quality Control metrics ("nCount\_RNA", "nFeature\_RNA", "percent.mito"), with cells grouped by **subset** annotation. y-axis for "nCount\_RNA" and "nFeature\_RNA" shows log values.

Supplementary Figure 5

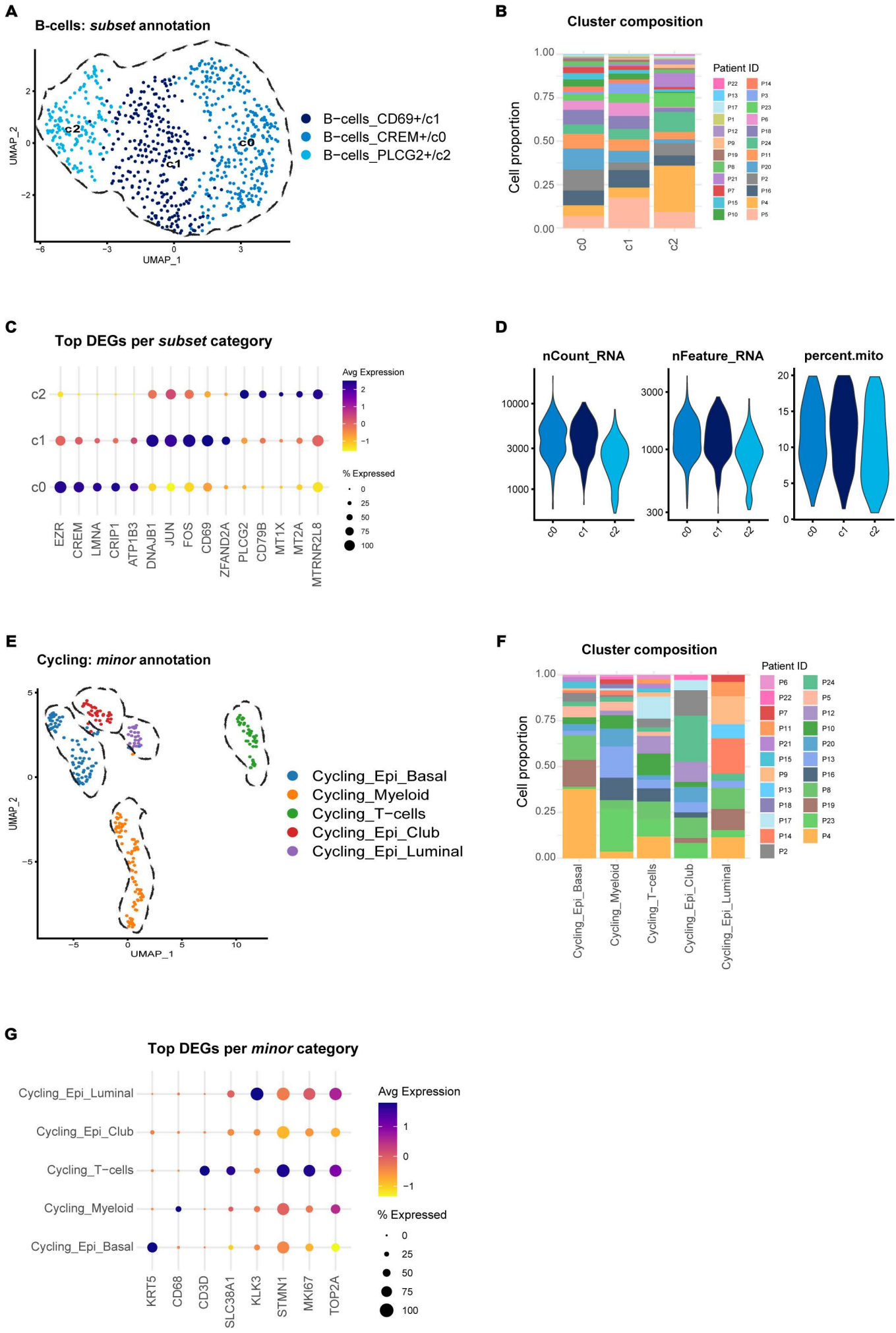

**Supplementary Figure 5:** Overview of B-cells and Cycling cells in the PCa TME.

**A-D:** B-cells; **E-G:** Cycling cells.

**A, E)** UMAP visualisation of **subset** annotation; dashed circles indicate **minor** lineages. For **E)**, Cycling cells were initially detected as part of the Epithelial and Immune subclustering analysis, then combined into a **major** “Cycling” category. **B, F)** **subset** cluster composition, filled by 24 unique patients (P1-P24). **C, G)** Average expression of the top 5 differentially expressed genes per **subset** category. **D, H)** Quality Control metrics (“nCount\_RNA”, “nFeature\_RNA”, “percent.mito”), with cells grouped by **subset** annotation. y-axis for “nCount\_RNA” and “nFeature\_RNA” shows log values.

#### Supplementary Figure 6

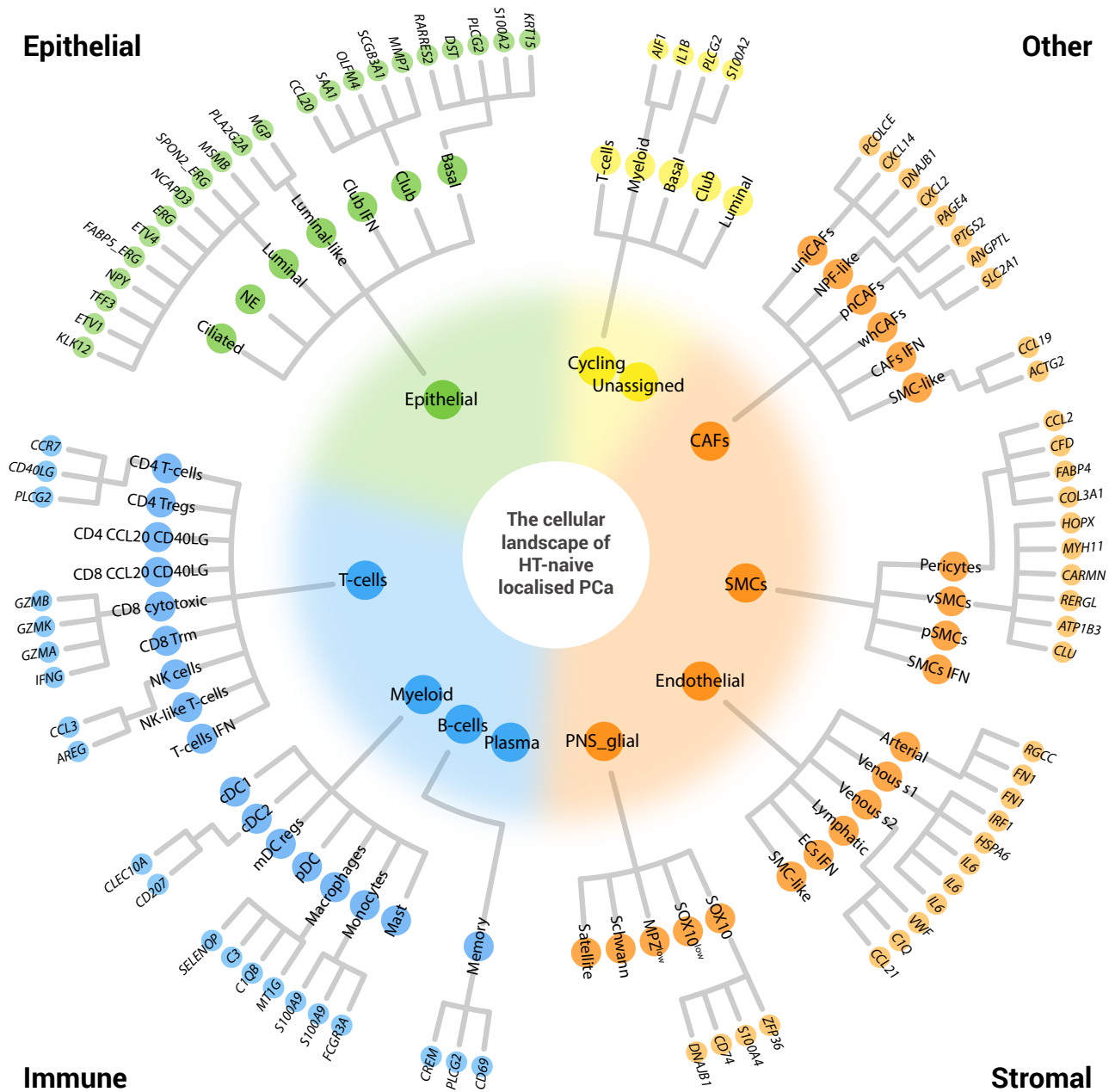

**Supplementary Figure 6:** A cellular taxonomy of hormone therapy (HT)-naive localised prostate cancer

Cell types are depicted across three levels: **major** (inner), **minor** (middle), and **subset** (outer). The **subset** level represents clusters and is labelled by the top differentially expressed gene in that cluster, when ranked by average logFC.

**IFN:** Interferon; **NE:** Neuroendocrine; **Tregs:** T-regulatory cells; **Trm:** T-resident memory cells; **NK:** Natural Killer; **cDC:** Conventional dendritic cells; **mDC regs:** mature dendritic regulatory cells; **pDC:** plasmacytoid dendritic cells; **CAFs:** cancer-associated fibroblasts; **uniCAFs:** universal CAFs; **NPF-like:** normal prostate fibroblast-like; **pnCAFs:** peri-neural CAFs; **whCAFs:** wound-healing CAFs; **SMCs:** smooth muscle cells; **vSMCs:** vascular SMCs; **pSMCs:** prostatic SMCs; **ECs:** endothelial cells; **s1:** state 1, **s2:** state 2. Some names are shortened for visualisation purposes. The full nomenclature is provided in ST\_15.

Supplementary Figure 7

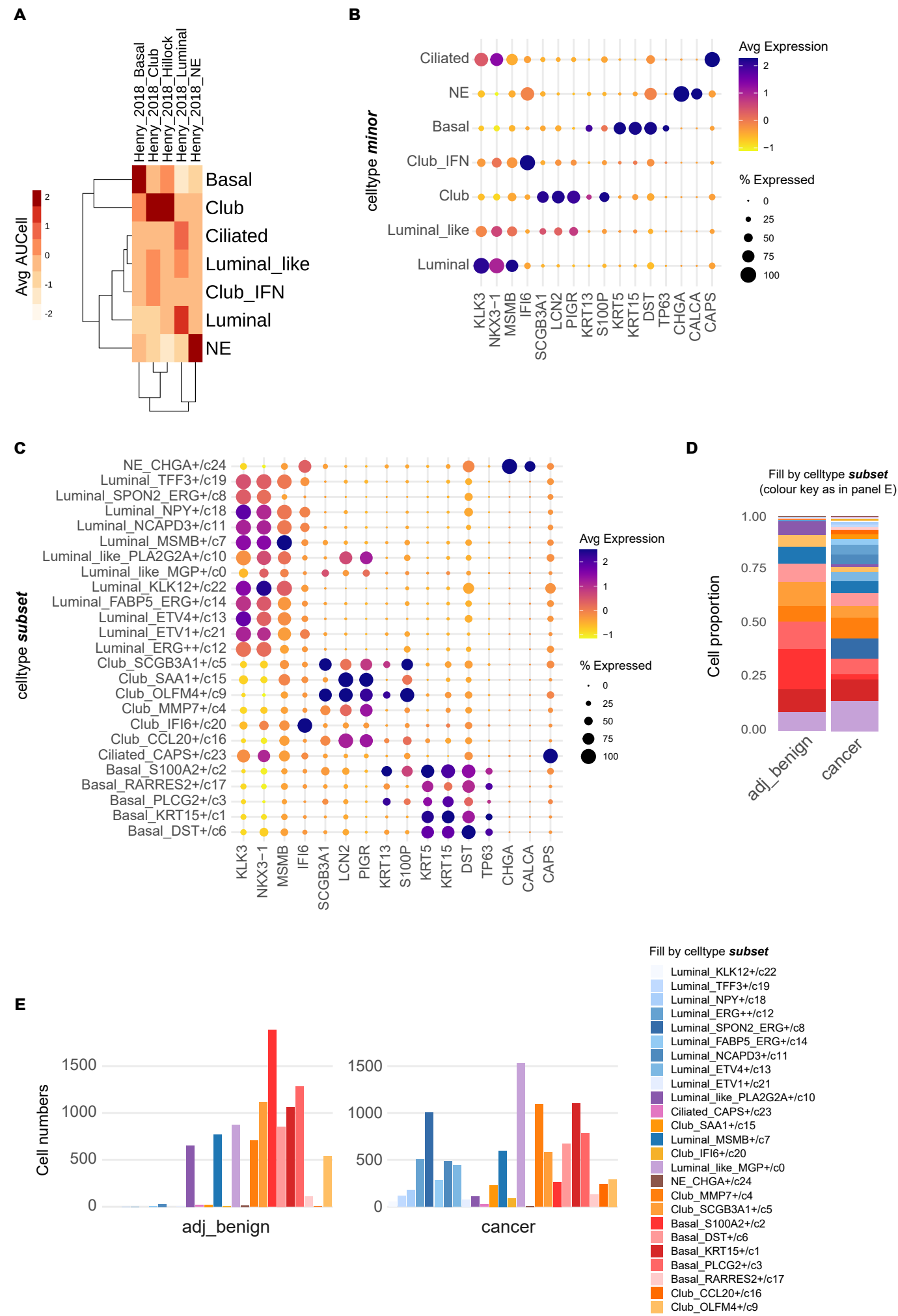

**Supplementary Figure 7:** Expression of canonical epithelial markers and cell type subset abundance in cancer and adjacent-benign cores

**A)** Average AUCell enrichment scores per **minor** category for signatures of prostatic epithelial lineages (Henry et al. 2018) and of TMPRSS2-ERG fusion (Setlur et al. 2008). **B)** Expression of Luminal (*KLK3*, *MSMB*, *NKX3-1*), Interferon (*IFI6*), Club (*SCGB3A1*, *LCN2*, *PIGR*), Hillock (*KRT13*, *S100P*), Basal (*KRT5*, *KRT15*, *DST*, *TP63*), Neuroendocrine (*CHGA*, *CALCA*) and Ciliated (*CAPS*) cell markers, with cells grouped by **minor** annotation. **C)** As in (B), with cells grouped by **subset** annotation. **D)** **subset** cell type abundance (%) by core type. **E)** **subset** cell type numbers by core type.

### Supplementary Figure 8

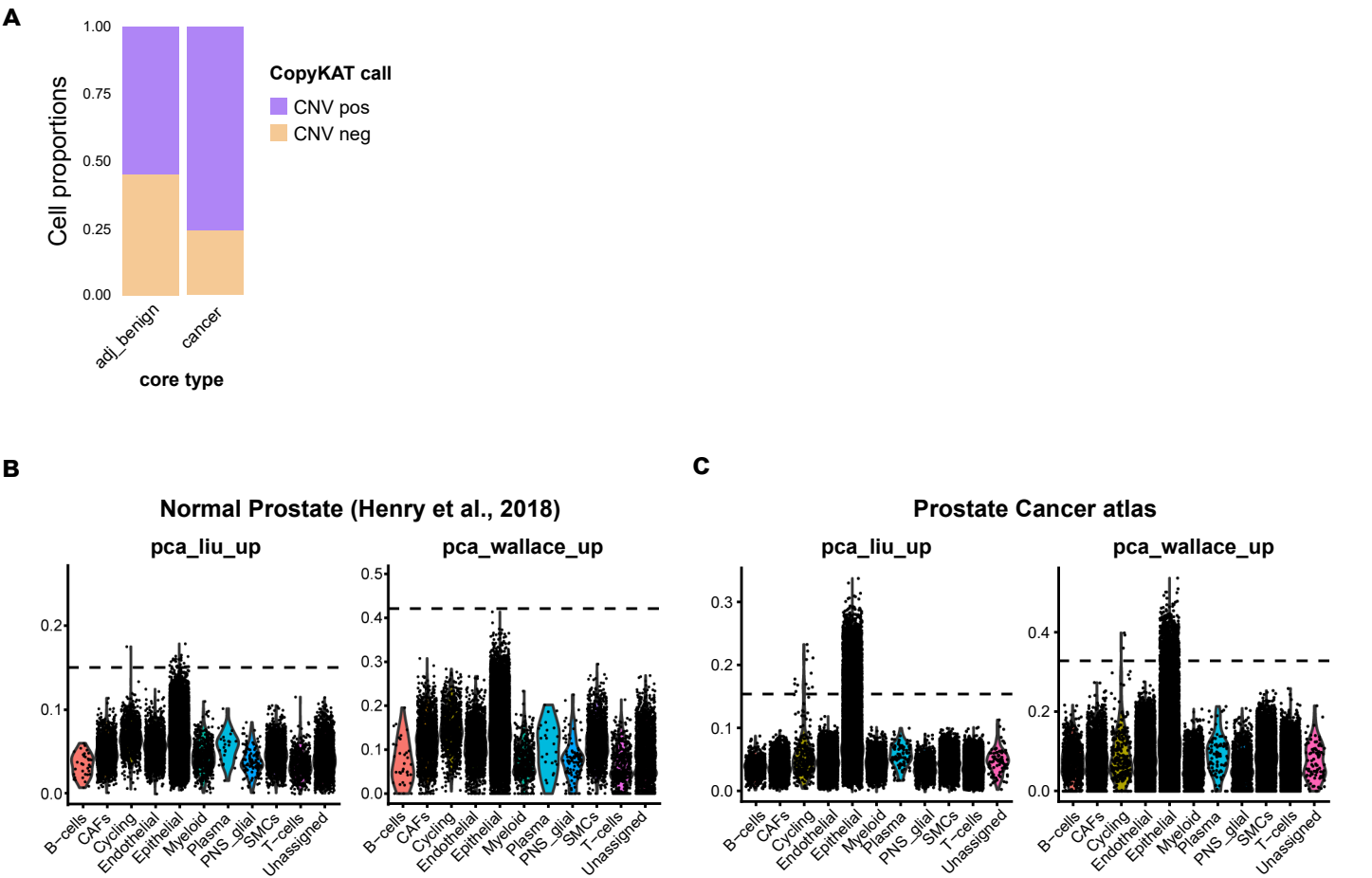

**Supplementary Figure 8:** Application of CopyKAT and published prostate cancer signatures to define the malignant spectrum

**A)** CopyKAT classifications of all Epithelial cells (CNV<sup>pos</sup>, CNV<sup>neg</sup>) using a normal-cell reference (all cells, excluding CAFs and unknown); split by core type. **B)** AUCell enrichment scores for “pca\_liu\_up” (left) and “pca\_wallace\_up” in Normal prostate scRNA-seq data (Henry et al., 2018). Dotted horizontal line represents the AUCell threshold for each signature. Cells are grouped by *major* annotation. **C)** As in (B), but for the PCa atlas data.

Supplementary Figure 9

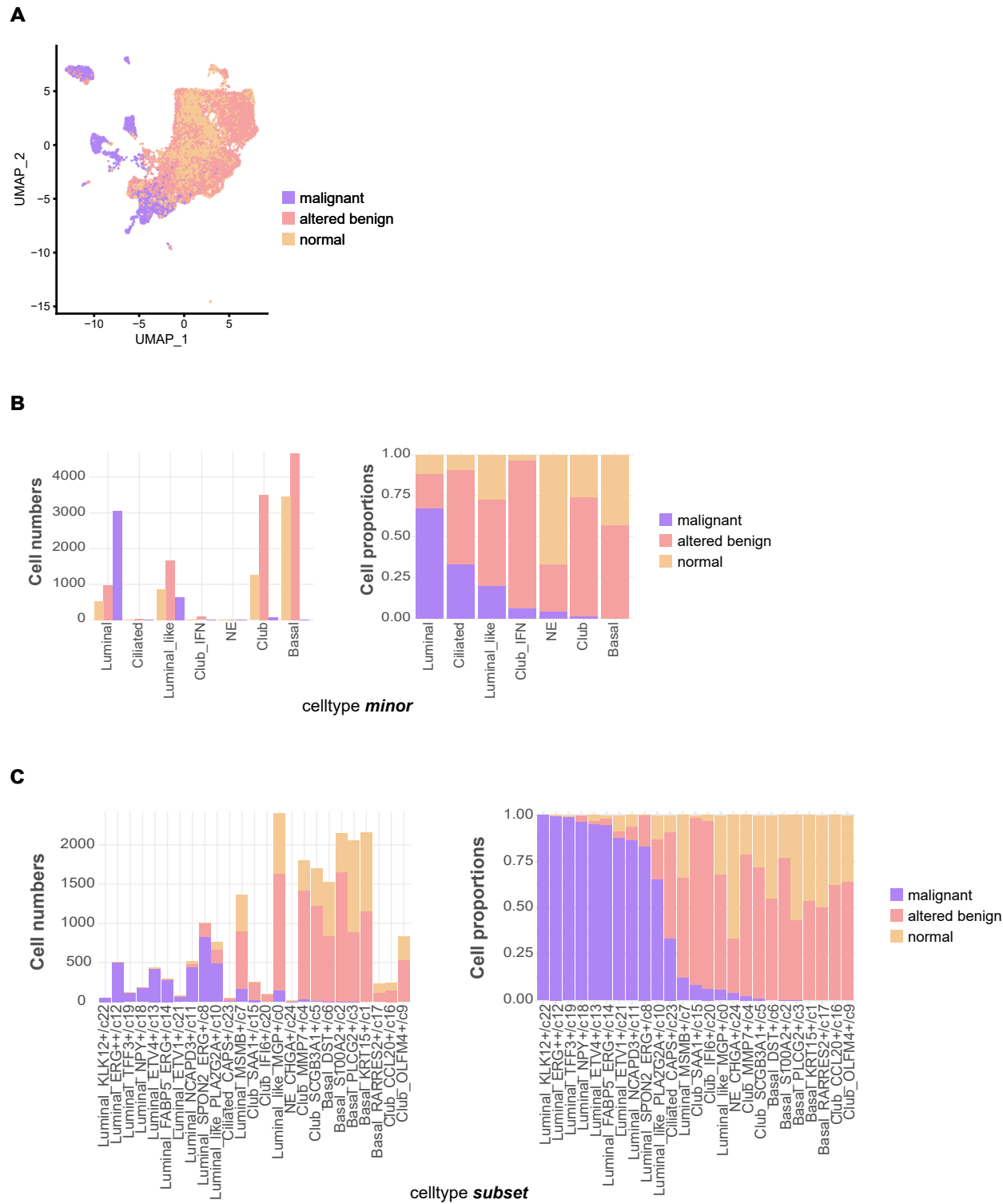

Supplementary Figure 9: Distribution of epithelial functional states across epithelial lineages and clusters

**A)** UMAP visualisation with malignant spectrum state annotation (normal, altered benign, malignant) projected. **B)** Number (left) and proportions (right) of cells, grouped by *minor* annotation. **C)** As in (B), with cells grouped by *subset* annotation. In B and C, the denominator is the total number of Epithelial cells, with colours indicating their malignant spectrum state. The cell clusters on the x-axis are ranked by proportion of malignant cells in each cluster.

Supplementary Figure 10

A Patient ID P4

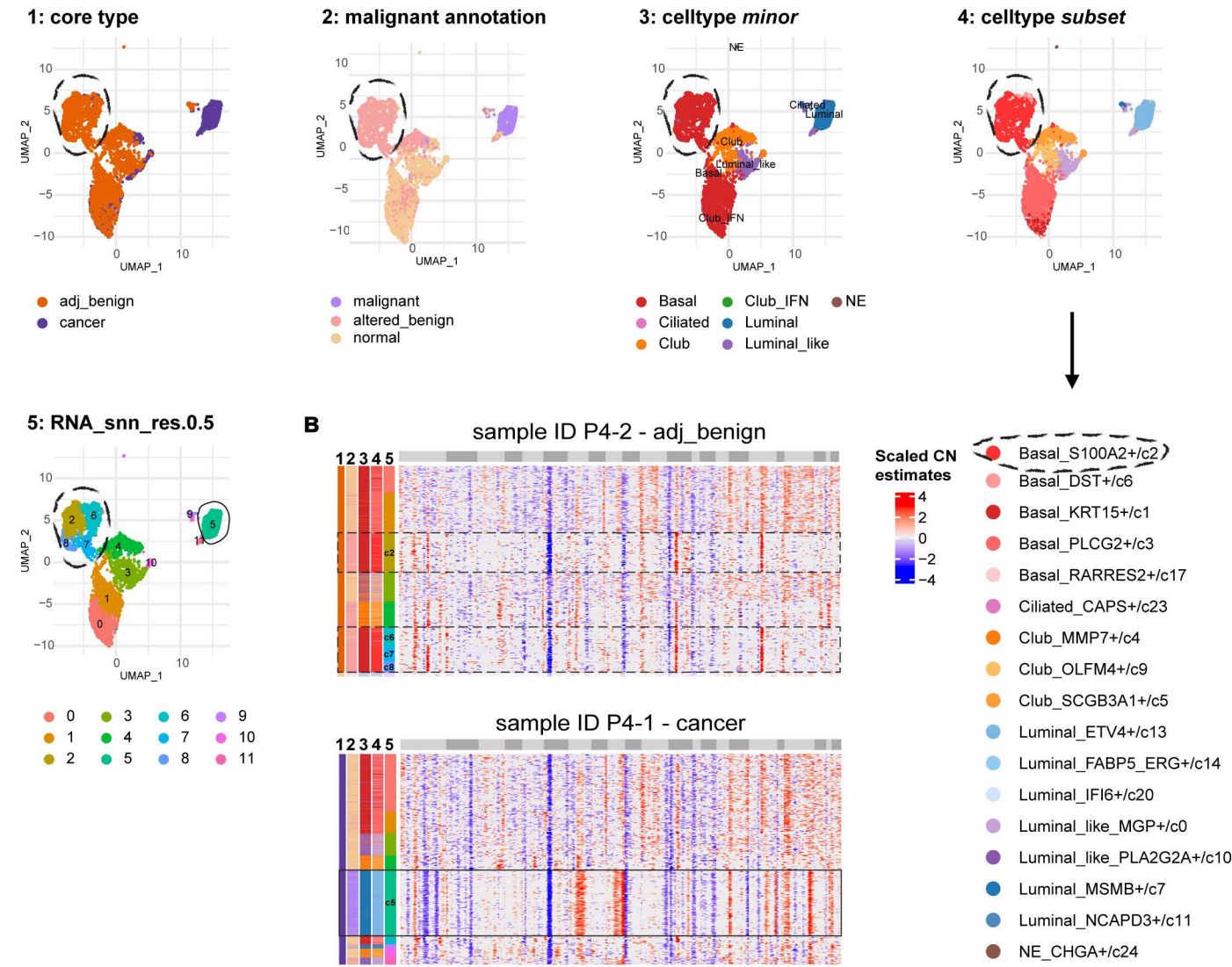

C Patient ID P10

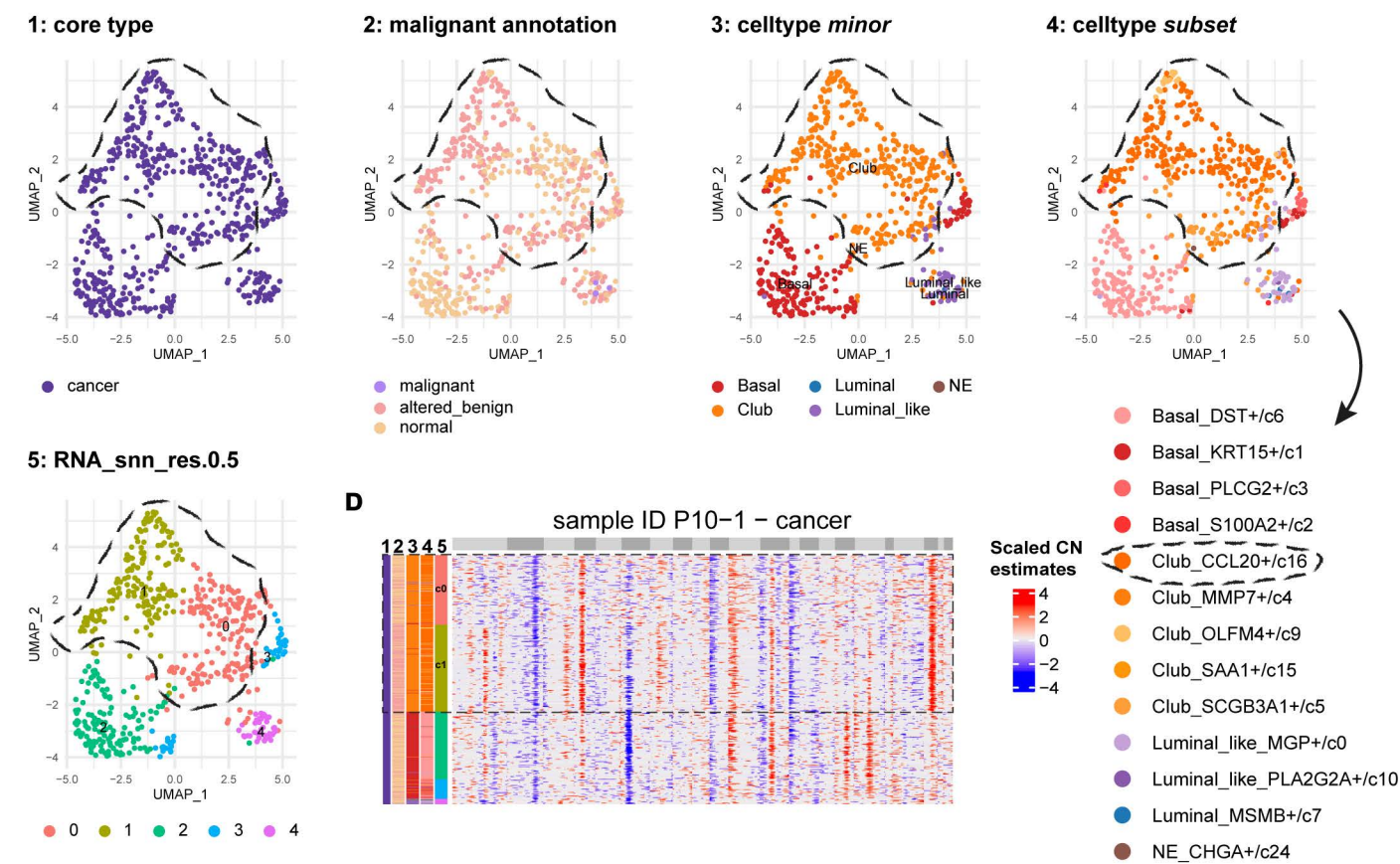

**Supplementary Figure 10:** Detection of copy number changes in Basal and Club cells (shown for two patients)

**A)** UMAP of reclustered epithelial cells in patient P4, showing (1) core type, (2) malignant, (3) *minor*, (4) *subset* annotation, and (5) Seurat clusters (RNA\_snn\_res.0.5). Clusters identified as "Basal\_S100A2+/c2" are circled with a dashed line. Malignant Luminal cluster (c5) is circled with a solid black line. **B)** Copy number estimates for samples P4-2 (adjacent-benign, top) and P4-1 (cancer, bottom). Sample P4-2 shows CNVs in "Basal\_S100A2+/c2" cells (clusters indicated with dashed lines). Row meta data show (1) core type, (2) malignant, (3) *minor*, (4) *subset* annotation, and (5) Seurat clusters. z-score scaled CNV signal is shown, with rows (cells) clustered by CNV signal. Row dendrograms are omitted. Chromosomes 1-22 are alternatively coloured (odd: light grey; even: dark grey). **C)** As in (A), but for patient P10. Clusters identified as "Club\_SAA1+/c16" are circled with a dashed line. **D)** As in (B), but for sample P10-1 (adjacent-benign).

Supplementary Figure 11

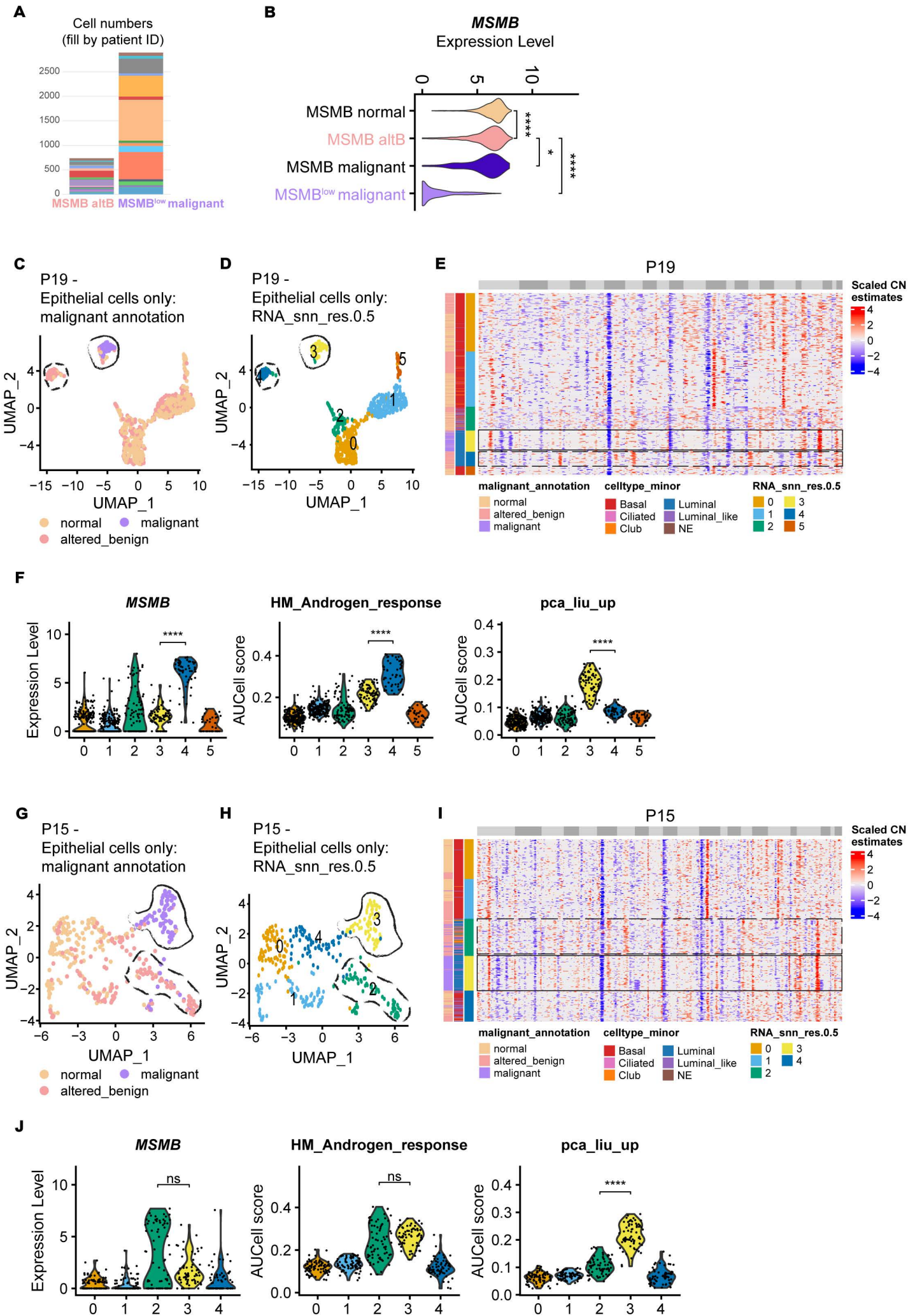

**Supplementary Figure 11:** Downregulation of androgen signalling in malignant cells compared to altB MSMB+ cells at the individual patient level (shown for two patients)

**A)** Cell numbers for altB MSMB+ cells and MSMB<sup>low</sup> malignant cells, coloured by patient ID (right). **B)** *MSMB* expression across MSMB<sup>low</sup> malignant cells and the three MSMB+ states. **C-F)** Results for patient P19. UMAP of reclustered epithelial cells, showing **C)** malignant spectrum annotation and **D)** Seurat clusters (c0-c5; RNA\_snn\_res.0.5). Dashed lines highlight altB MSMB+ (c4) and solid lines highlight MSMB<sup>low</sup> malignant (c3) cells. **E)** Copy number estimates for patient P19 (both cores), showing CNVs in altB MSMB+ cells. Row meta data show (1) malignant spectrum, (2) *minor* and (3) Seurat cluster annotation for each cell. z-score scaled CNV signal is shown, with rows (cells) ordered by Seurat clusters. Chromosomes 1-22 are alternatively coloured (odd: light grey; even: dark grey). Relevant gene expression clusters are indicated with dashed (altB MSMB+) and solid (MSMB<sup>low</sup> malignant) line rectangles. **F)** High *MSMB* expression in c4 and enrichment of the 'pca\_liu\_up' signature in c3. Means compared with Wilcoxon test (unadjusted p-values; $p \leq 0.001$ (\*\*\*\*)). **G-J)** As in (C-F), but for patient P15, comparing altB MSMB+ (c2) and MSMB<sup>low</sup> malignant cells (c3).

#### Supplementary Figure 12

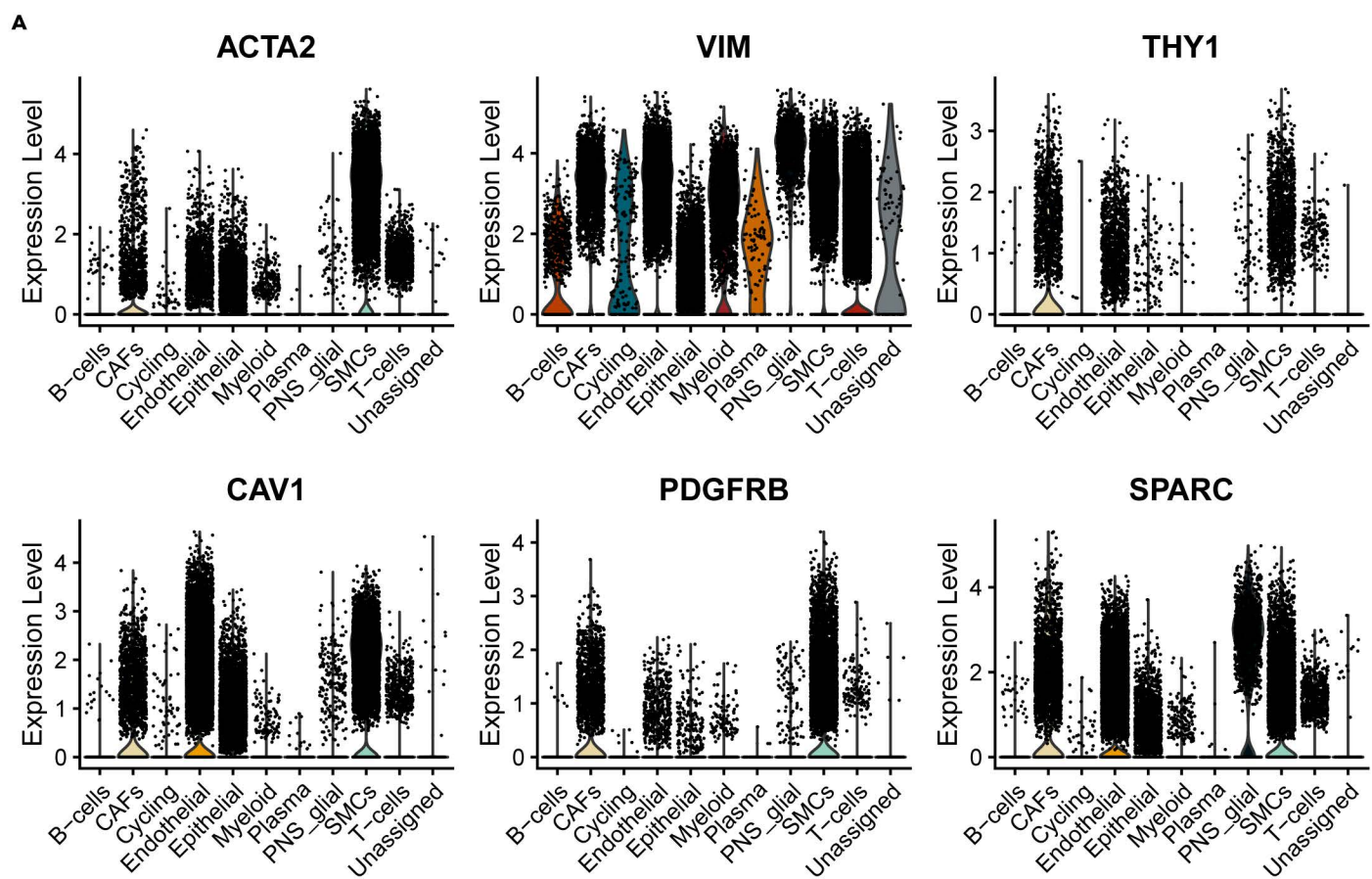

**Supplementary Figure 12:** Genes canonically used for the identification of CAFs are expressed by various stromal cell types

**A)** Expression of *ACTA2*, *VIM*, *THY1*, *CAV1*, *PDGFRB* and *SPARC*. Cells are grouped by **major** annotation

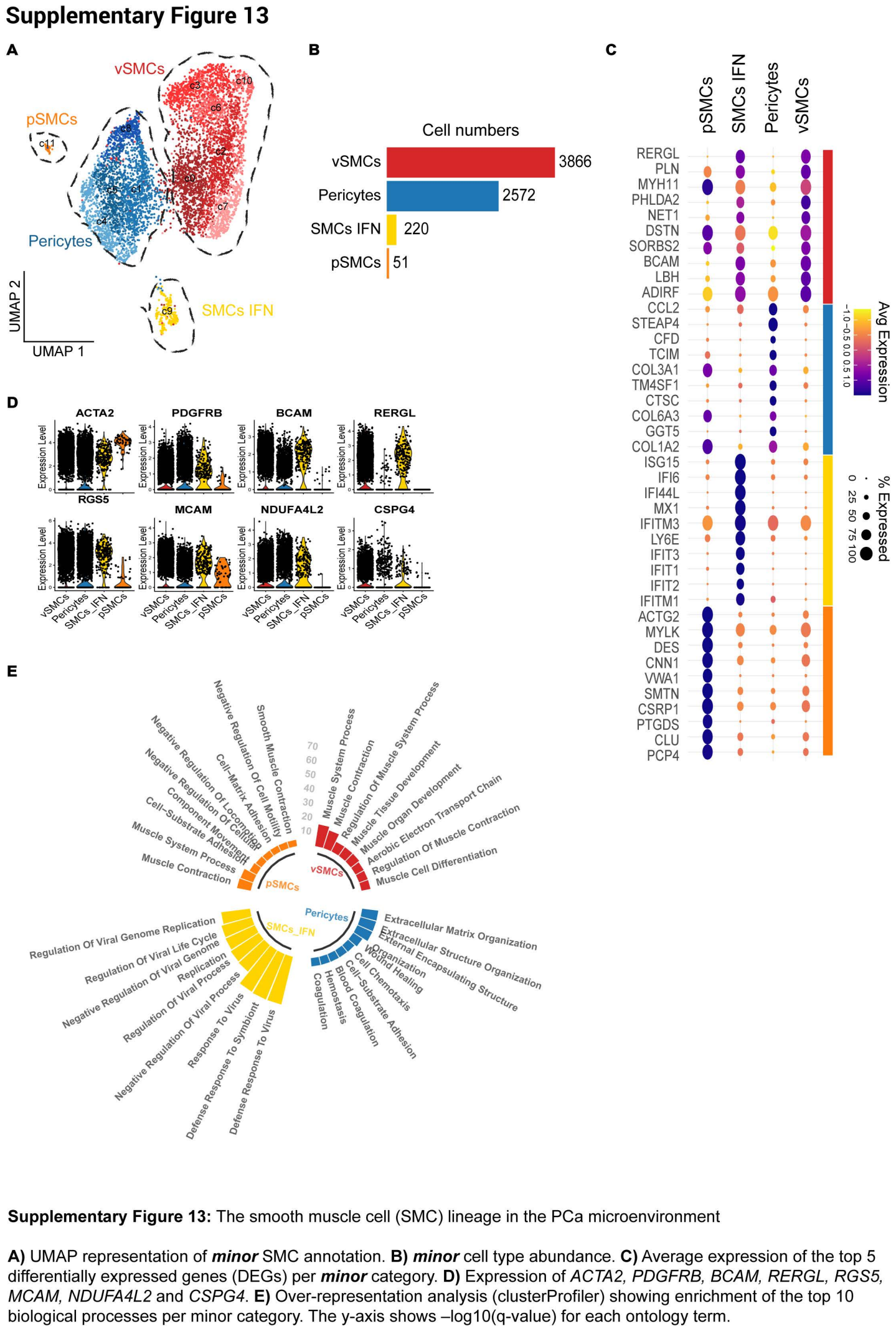

### Supplementary Figure 14

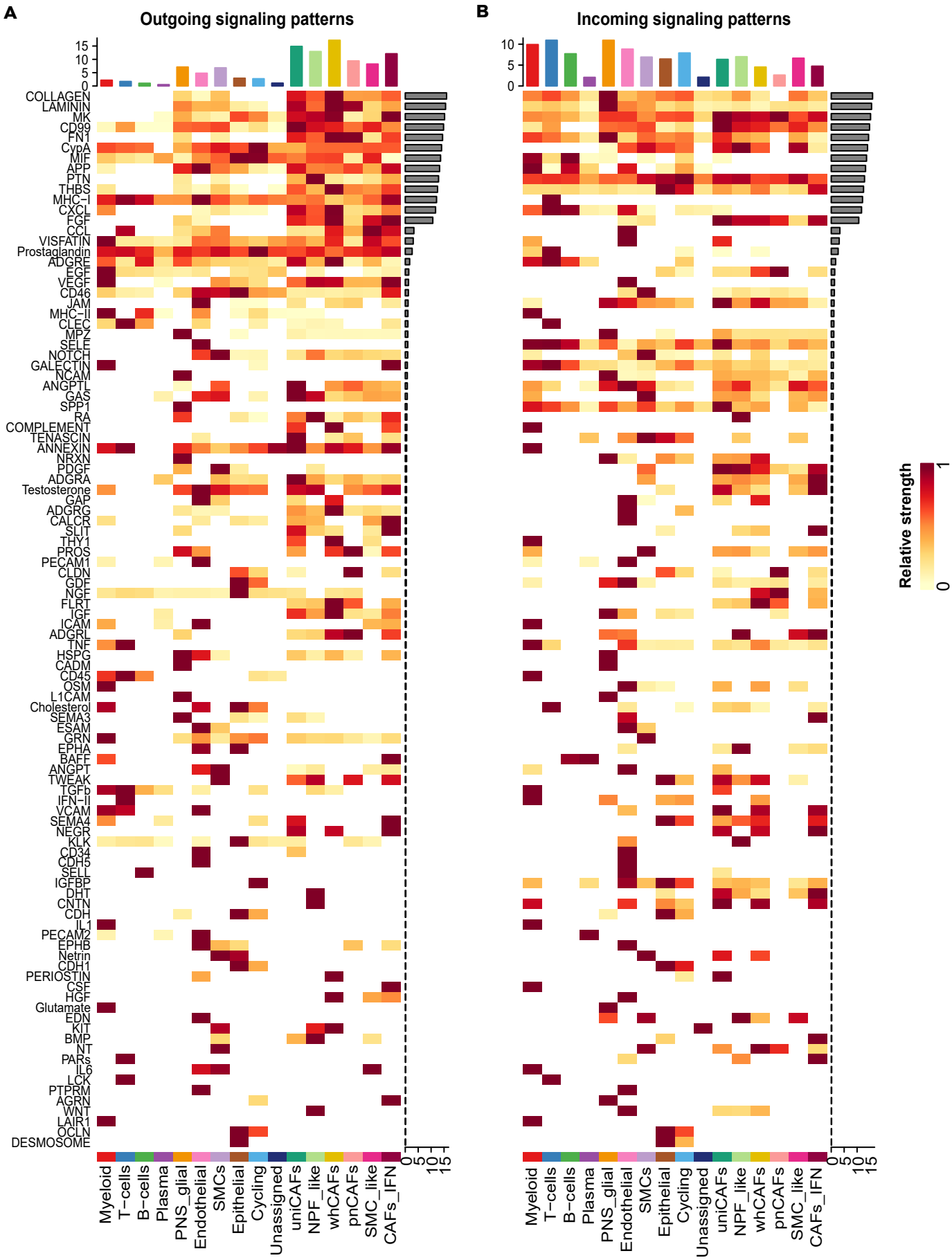

**Supplementary Figure 14:** CellChat outgoing and incoming signals for all signalling pathways

CellChat heatmaps displaying selected outgoing (**A**, cells are source) and incoming (**B**, cells are targets) signals (x-axis) across all cell types. Cell types are grouped by **major** category, except for CAFs, which are grouped by **minor** category (x-axis). Colour values indicate relative strength of each signalling pathway, scaled row-wise per heatmap. Grey barplot (right indicates overall signalling strength for each pathway; coloured barplot (top) shows total signalling strength per cell type (all pathways summarised). A selected list of signalling pathways is shown in Fig. 4G, with the panels in a different orientation.

Supplementary Figure 15

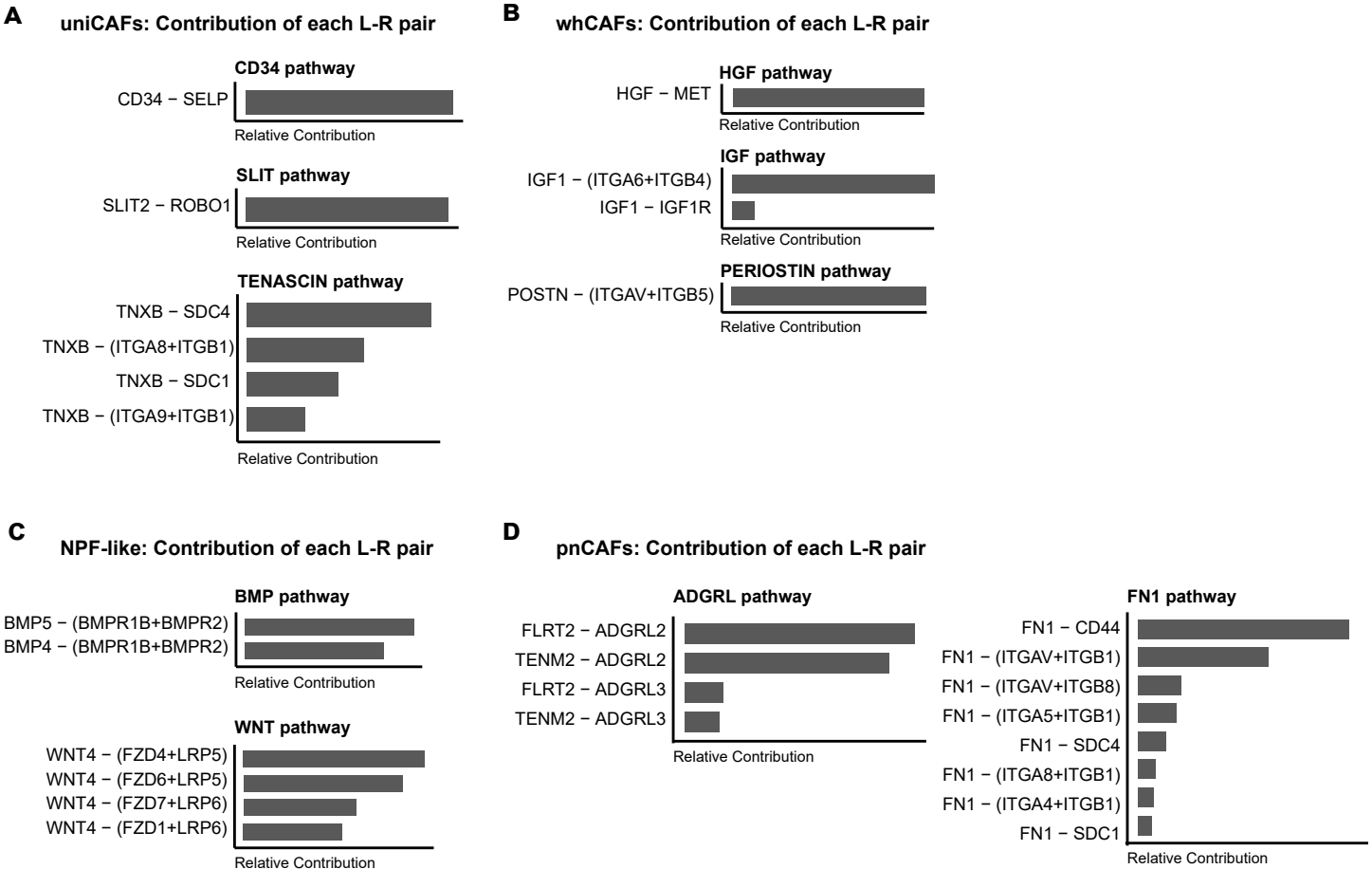

**Supplementary Figure 15:** Contribution of ligand-receptor (L-R) molecules driving pathway enrichment in CAF subtypes

Plots showing the relative contribution of ligand-receptor (LR) pairs, with CAF subtypes as ligand sources, and all other cell types as receptor targets. L-R analysis performed using CellChat, grouping cell types by *major* category, except CAFs by *minor* category. Only L-R pairs with p.value ≤ 0.05 are shown.

**A)** CD34, SLIT, and TENASCIN pathways, with uniCAFs as ligand source. **B)** HGF, IGF, and PERIOSTIN pathways, with whCAFs as ligand source. **C)** BMP and WNT pathways, with NPF-like cells as ligand source. **D)** ADGRL and FN1 pathways, with pnCAFs as ligand source.

Supplementary Figure 16

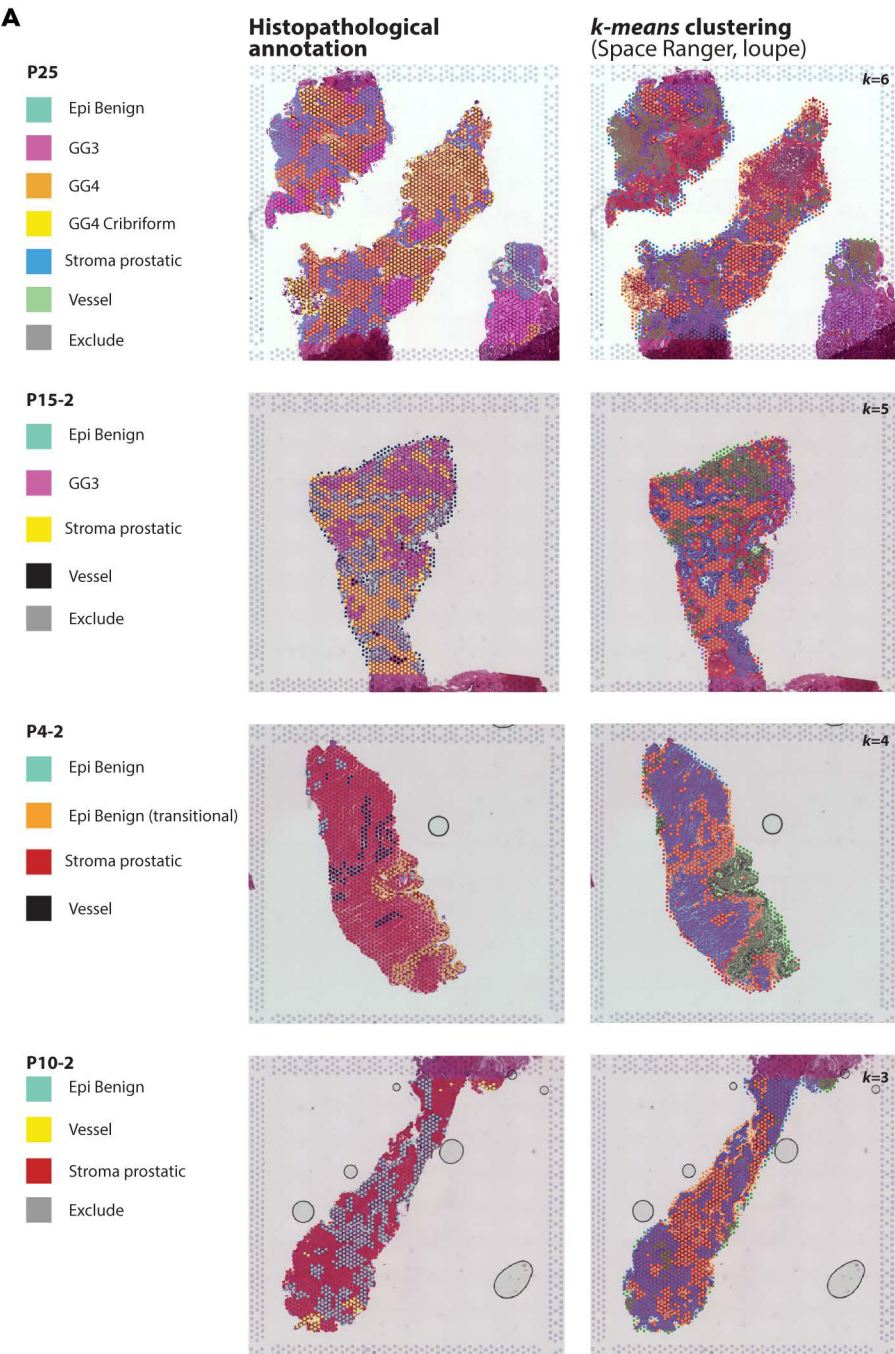

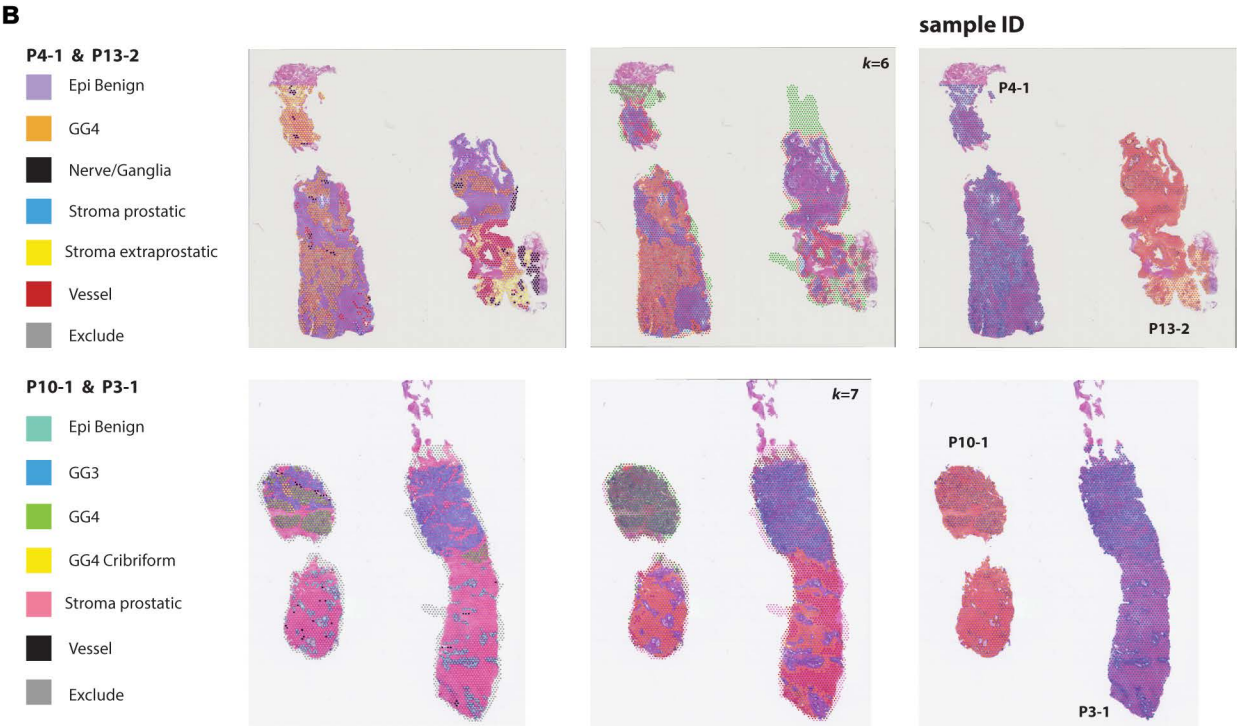

**Supplementary Figure 16:** Spatial plots showing histopathological annotations and k-means clustering for additional 8 Visium samples (addition to Fig. 5A).

**A)** Visium FFPE V1 samples (n=4), 6.5 x 6.5mm reactions. **B)** Visium FFPE V2 samples (n=4), 11 x 11mm reactions. Supplementary Figure 16: Spatial plots showing histopathological annotations and k-means clustering for additional 8 Visium samples (addition to Fig. 5A).

Histological features are indicated by the legends on the left. Colours are mismatched between patients to best reflect visual differences between annotations. *k*-means clusters, derived from default Space Ranger analysis, are visualised using the Loupe browser. Different colours indicate different clusters; cluster numbers are not indicated. In Panel B, the third column indicates two different sample IDs per Visium reaction.

### Supplementary Figure 17

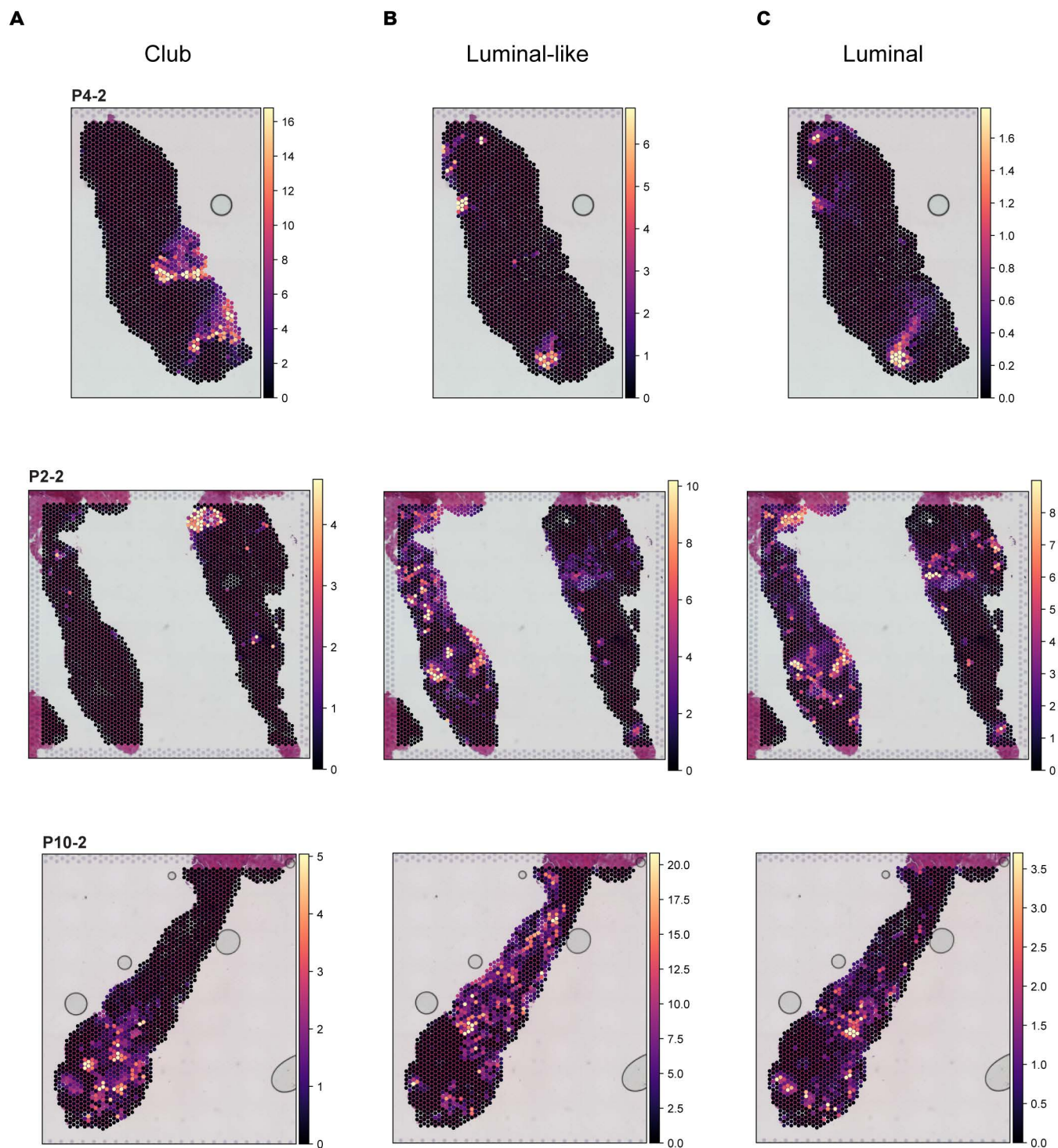

**Supplementary Figure 17:** Cell2Location results for Club, Luminal-like and Luminal cells in adjacent-benign cores

Per-spot deconvolution values for **A**) Club, **B**) Luminal-like, and **C**) Luminal cells. Each row is a new sample (P4-2, P2-2, P10-2). Histopathological annotation for each sample is shown in Supp. Fig. S16.

Supplementary Figure 18

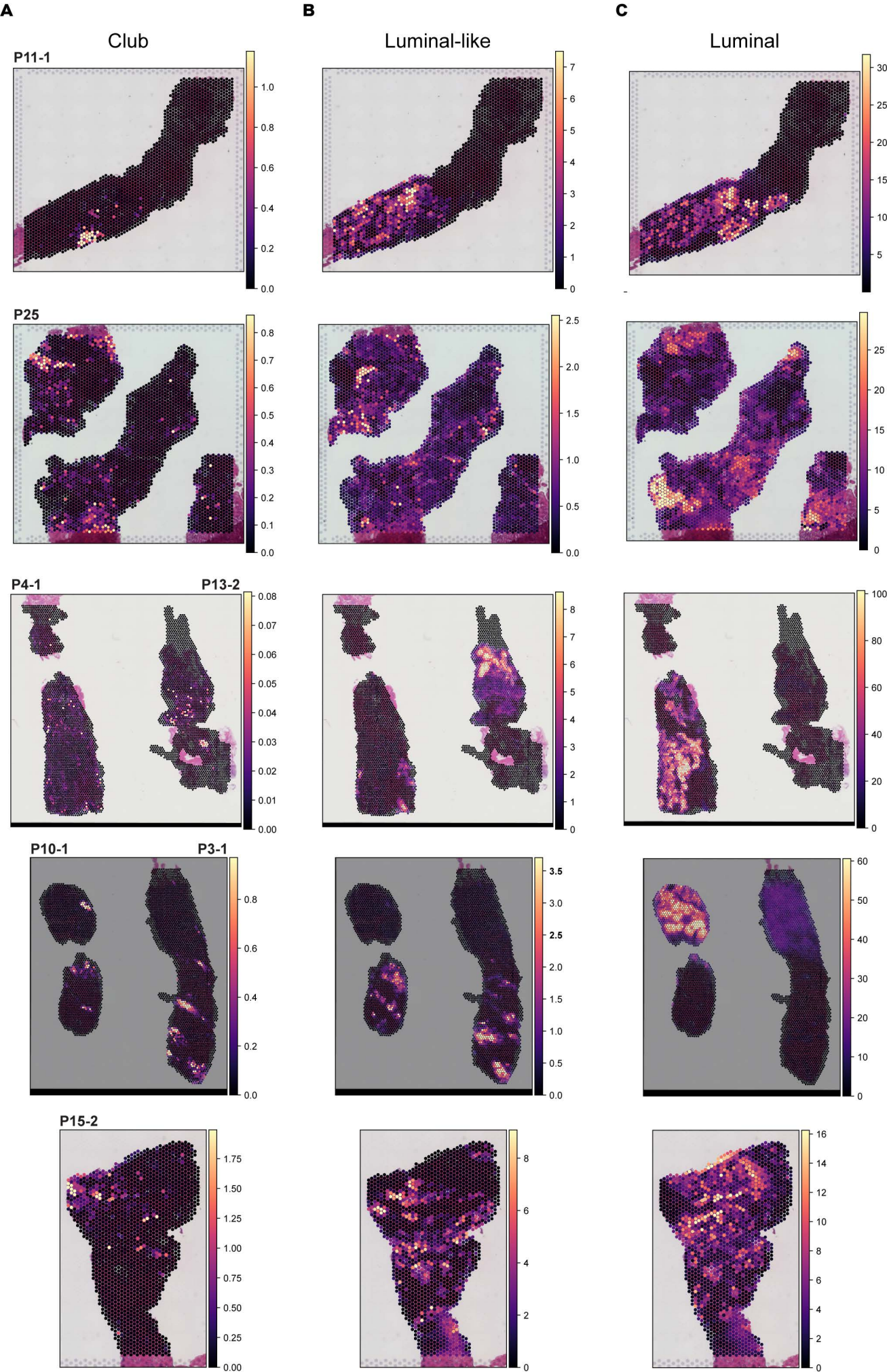

**Supplementary Figure 18:** Cell2Location results for Club, Luminal-like and Luminal cells in cancer cores

Per-spot deconvolution values for **A)** Club, **B)** Luminal-like, and **C)** Luminal cells. Each row is a new sample (P11-1, P25, P4-1 & P13-2, P10-1 & P3-1, P15-2). Histopathological annotation for each sample is shown in Supp Fig. S15.

Supplementary Figure 19

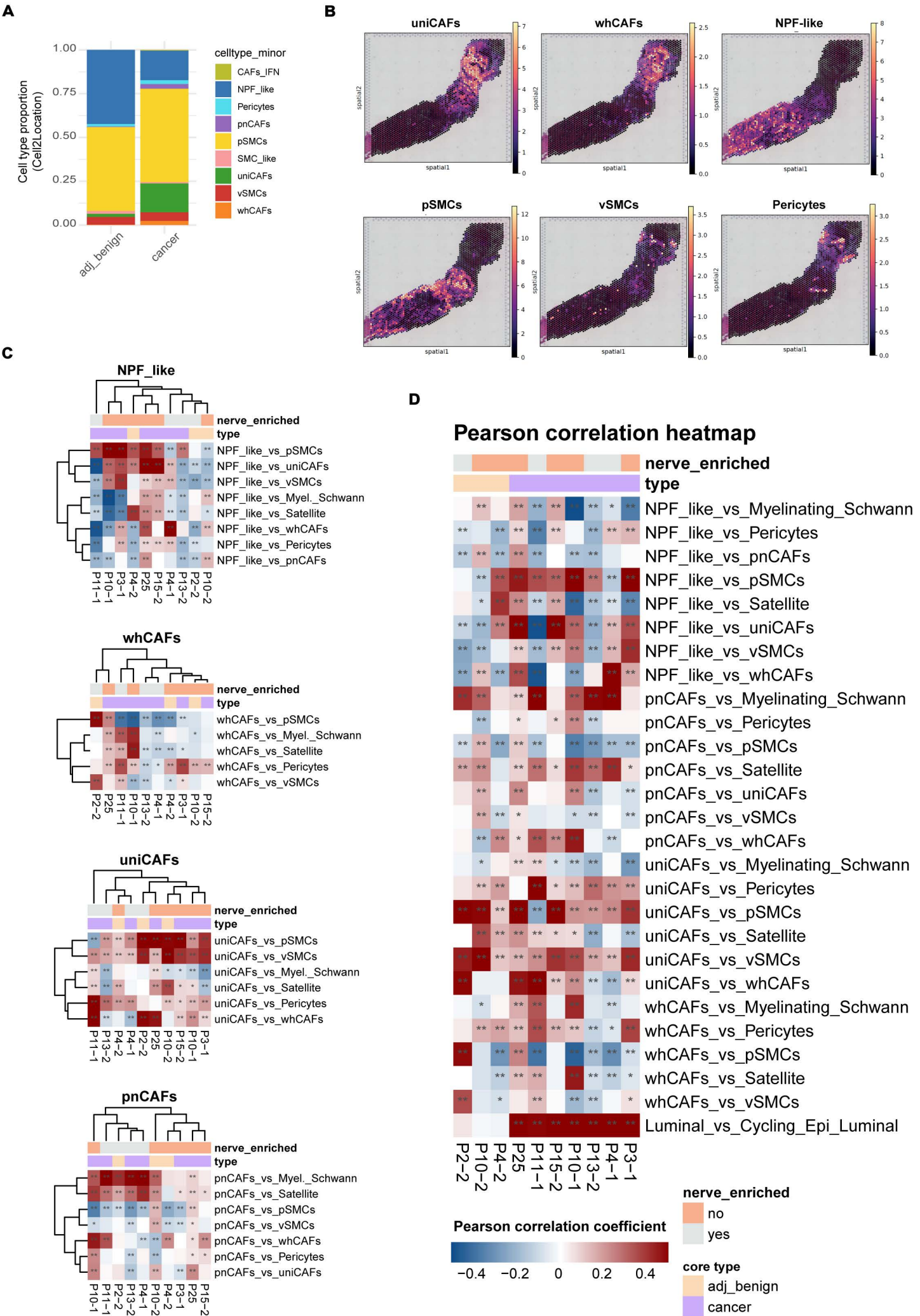

##### Supplementary Figure 19: Cell2Location deconvolution and correlation analysis

**A)** Cell2Location deconvolution proportions for CAF and SMC *minor* cell types per core type (x-axis) across 10 samples. Cell types constituting <5% of a spot are filtered out. **B)** Cell2Location per-spot deconvolution values for uniCAFs, whCAFs, NPF-like cells, pSMCs, vSMCs, and Pericytes; shown for sample P11-1. **C)** Clustered heatmap of Pearson correlation values between *minor* CAF subtypes, SMC, and PNS\_glial subtypes across 10 samples (3 adjacent-benign, 7 cancer). Shown per CAF *minor* subtype (NPF-like cells, whCAFs, uniCAFs, pnCAFs). Nerve-containing samples are indicated as 'nerve-enriched'. Asterisks denote Benjamini-Hochberg adjusted p-values: \* < 0.01, \*\* < 0.001. **D)** Similar to C), but showing all CAF cell types combined and without clustering rows and columns, representing the full comparison shown in Fig. 5C.

**A**

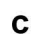

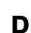

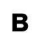

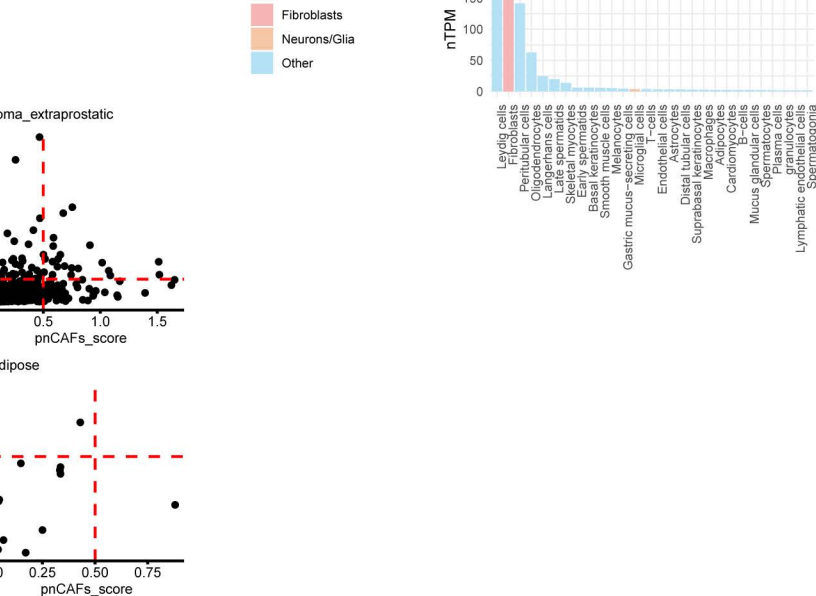

#### Supplementary Figure 20: Cepo-derived pnCAF gene signature

**A)** Average expression for nine Cepo-derived pnCAF genes (used in Scanpy scoring of Visium data) in the scRNA-seq dataset, with cells grouped by cell type **major temp** (CAFs at **minor**, everything else at **major**). Dot size represents the percentage of cells expressing each gene. **B)** Expression (nTPM) of nine pnCAF genes (used in Scanpy scoring of Visium data) in Human Protein Atlas RNA single-cell type data. Only cell types with nTPM > 1 per gene are shown on the x-axis. Bars are coloured by cell type category (red = fibroblasts, yellow = neurons or glia, blue = other cell types). **C)** Per-spot Pearson correlation between Cepo-derived PNS\_glial and pnCAFs gene sets across 10 Visium samples. Scores are calculated using ``scanpy.tl.score_genes()``, representing the difference between the average expression of each gene set and a randomly sampled reference gene set. Negative scores, indicating lower average expression compared to the reference, are filtered out. Dashed lines indicate score thresholds of 0.5. Each spot is coloured by whether it derives from a nerve-enriched sample. **D)** Unique histologies in nerve-enriched samples only (4/10), with thresholds for each signature indicated by dashed lines (0.5, arbitrary). Nerves are highlighted with a peach rectangle.

### Supplementary Figure 21

A

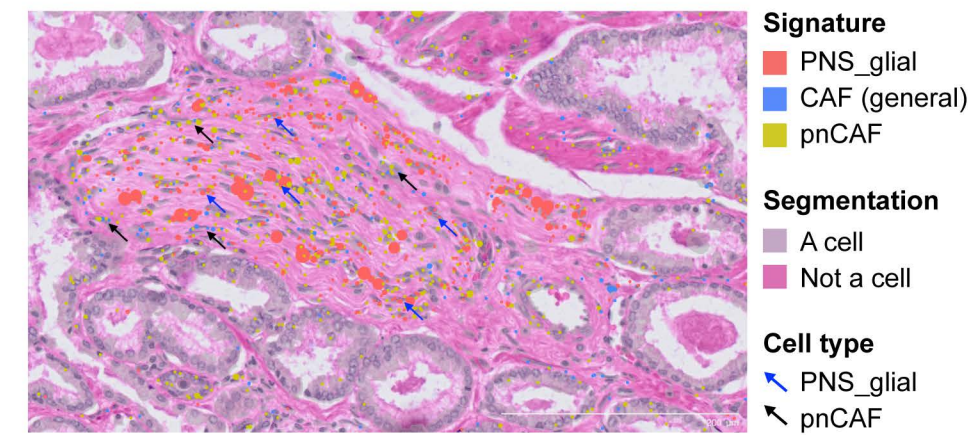

**Supplementary Figure 21:** PNS\_glial and pnCAF signatures are enriched in distinct cell types within nerve bundles in published Xenium spatial transcriptomics data from a prostate adenocarcinoma sample.

**A)** Zoomed-in view of a nerve bundle. Distinct colours represent summarised transcript expression for specific cell types, using Cepo-derived signatures for PNS\_glial and pnCAFs, and a manually curated signature for CAF (general). PNS\_glial and pnCAF cells are marked with blue and black arrows, respectively. Scale bar = 200 μm.

Supplementary Figure 22

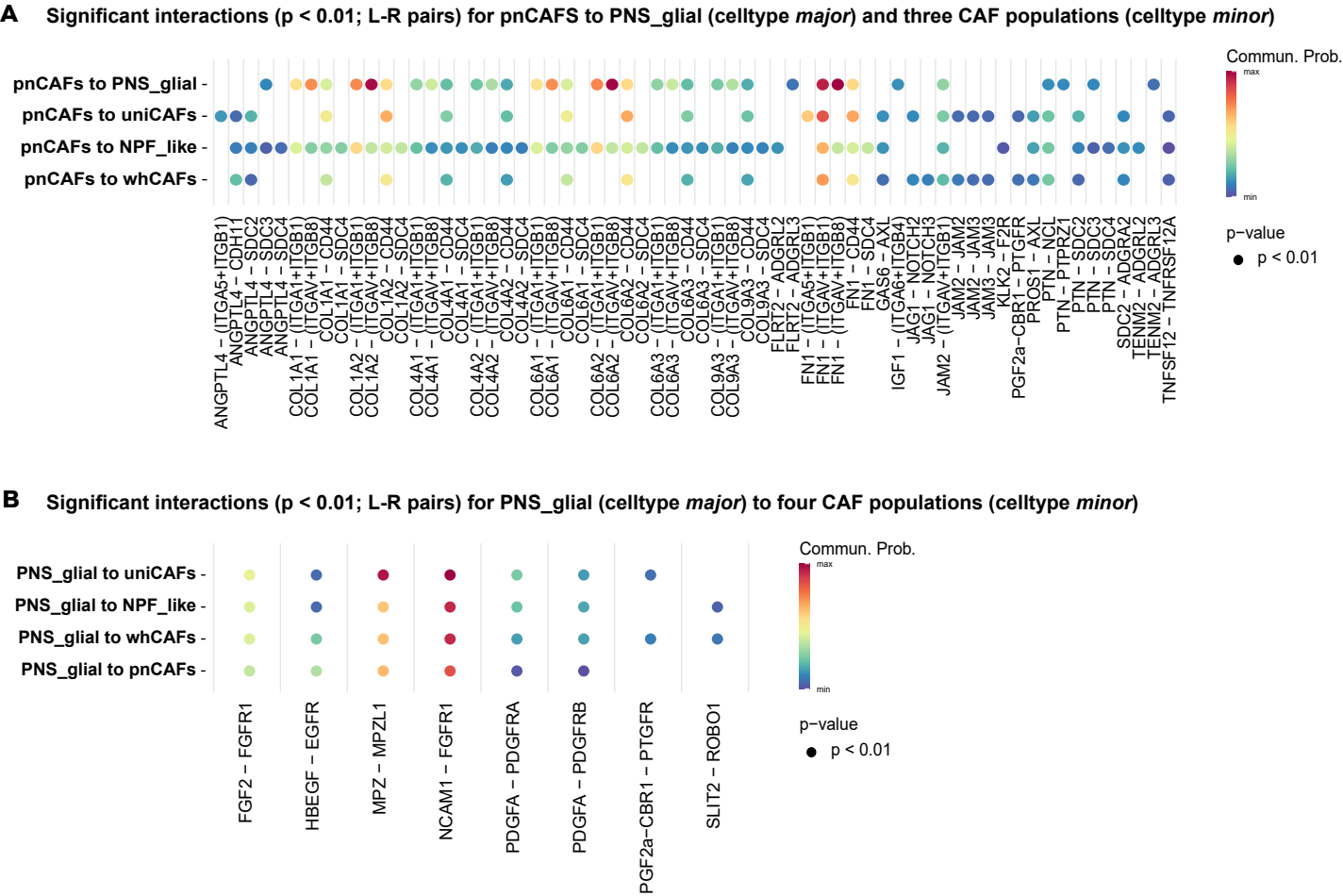

**Supplementary Figure 22:** Significant interactions between pnCAFs, PNS\_glia cells, and other CAF subtypes

**A)** Significant ligand-receptor interactions with pnCAFs as the ligand source, targeting PNS\_glia cells and the remaining CAF subtypes. **B)** Significant ligand-receptor interactions with PNS\_glia cells as the ligand source, targeting pnCAFs and the remaining CAF subtypes.

Only selected ligand-receptor pairs, with a $p\text{-value} \leq 0.01$, are shown.

Supplementary Figure 23

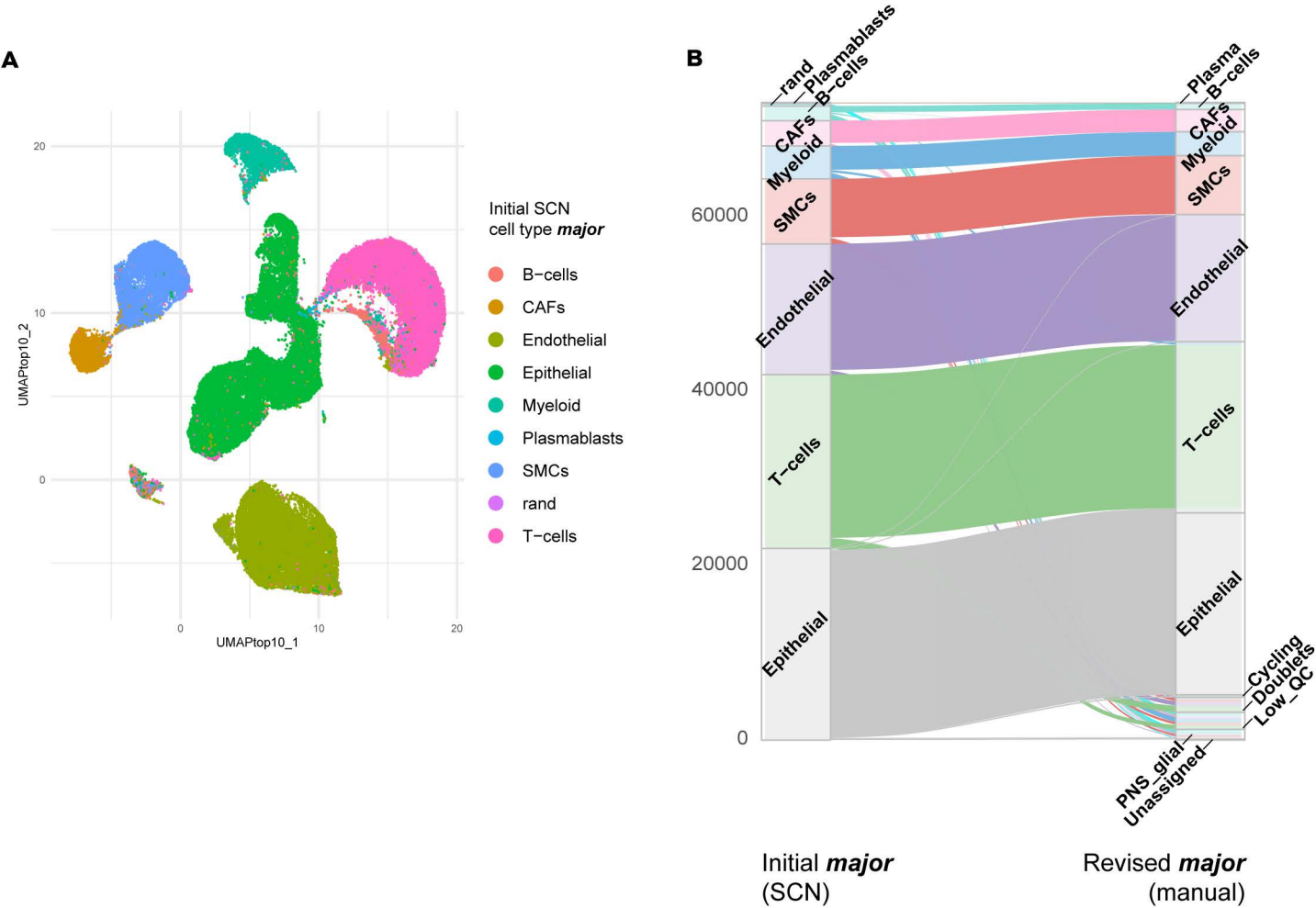

**Supplementary Figure 23:** Preliminary annotation of the PCa atlas dataset using SingleCellNet (SCN)

**A)** UMAP of the PCa atlas dataset, showing cells with temporary SCN annotation, trained on a breast cancer scRNA-seq dataset (Wu et al. 2021). **B)** Changes in cell annotation from initial SCN annotation to final manual annotation for *major* cell types; y-axis = cell count; Low\_QC and Doublets cells were filtered out.
